# Automatic error control during forward flux sampling of rare events in master equation models

**DOI:** 10.1101/254896

**Authors:** Max C. Klein, Elijah Roberts

## Abstract

Enhanced sampling methods, such as forward flux sampling (FFS), have great capacity for accelerating stochastic simulations of nonequilibrium biochemical systems involving rare events. However, the description of the tradeoffs between simulation efficiency and error in FFS remains incomplete. We present a novel and mathematically rigorous analysis of the errors in FFS that, for the first time, covers the contribution of every phase of the simulation. We derive a closed form expression for the optimally efficient count of samples to take in each FFS phase in terms of a fixed constraint on sampling error. We introduce a new method, forward flux pilot sampling (FFPilot), that is designed to take full advantage of our optimizing equation without prior information or assumptions about the phase weights and costs along the transition path. In simulations of both single- and multi-dimensional gene regulatory networks, FFPilot is able to completely control sampling error. Higher dimensional systems have additional sources of error and we show that this extra error can be traced to correlations between phases due to roughness on the probability landscape. Finally, we show that in sets of simulations with matched error, FFPilot is on the order of tens-to-hundreds of times faster than direct sampling, in a fashion that scales with the rarity of the events.

## I. INTRODUCTION

Understanding how inanimate biochemical molecules come together and interact to form a living cell is one of the fundamental goals of biology^1^. The cell’s state can be described as a point in a high dimensional space, where each dimension corresponds to the concentration of a different molecule. Complex, nonequilibrium regulatory and signaling networks connect these molecules through positive and negative interactions, naturally resulting in a large number of metastable states within the space^2^, representing different cellular phenotypes. Projected into two dimensions, the phase space appears as a rough landscape, sometimes referred to as a quasipotential^2^, epigenetic^3^, or phenotypic^4^ landscape. Stochastic fluctuations cause the cell’s state to move along the landscape and occasionally jump between metastable states.

Metastable systems, because they depend upon random fluctuations, are typically modeled using a formulation of the chemical master equation^5, 6^ or using stochastic differential equations^7^. In the former case, the models are often numerically studied using the stochastic simulation algorithm^8, 9^ (SSA) or one of its many varieties^10–12^. Such simulations have provided insight into diverse biological processes, including: the lysis/lysogeny decision in bacteriophage *λ*^13^, the *lac* operon in *Escherichia coli* ^14^, check-pointing during the cell cycle^15^, differentiation of stem cells^16^, the binding of an intrinsically disordered peptide to a protein^17^, macrophage regulation^18^, and gradient detection during yeast mating^19^.

Transitions between metastable states in stochastic biochemical systems are infrequent in that one must wait a long time to observe a large fluctuation that causes the system to switch states^20^. The time spent in the transition region is also very short relative to the waiting time^21^. Such dynamics are known as rare events, and are expensive to simulate as most of the computational effort is spent simply simulating the waiting state. Numerical methods for improving the efficiency of simulating rare events, generally known as enhanced sampling (ES) techniques, have a long history. The earliest work in the field is typically credited to Kahn *et al.*^22^, but has since been applied to the study of many systems in molecular mechanics. The key assumption is that metastable systems exhibit a large barrier separating the states on some free energy landscape. By biasing the simulation toward the transition path between the states one can more efficiently recover its free energy profile and, thus, the transition rates. For example: umbrella sampling^23, 24^ uses overlapping biasing potentials to confine multiple simulations to narrow windows along the path; metadynamics^25, 26^ gradually adds a repulsive potential to low points on the free energy surface causing the system to explore low probability regions of phase space and to eventually cross the barrier; transition path sampling^27–30^ generates a statistically correct set of transition paths starting from an initial trial path using acceptance and rejection criteria; weighted ensemble^31, 32^ runs multiple independent trajectories while dynamically splitting and merging them, with careful accounting of the trajectory weights, to balance simulations along the transition path. In general, a final unbiasing and/or recombination step is always needed to calculate unbiased statistics.

Although stochastic biochemical systems are described by quasipotential rather than free energy landscapes, the underlying physics is compatible with ES. Consequently, many varieties of ES can be applied to stochastic biochemical systems with rare event dynamics. Three of the most well known candidates are forward flux sampling (FFS)^33–36^, nonequilibrium umbrella sampling^37–39^, and weighted ensemble^40–44^. Although much discussion has taken place regarding the strengths and weaknesses of each of these methods^45, 46^ there is no consensus as to an optimal approach. In the case of a transition between only two metastable states along a single order parameter, FFS appears to be a reasonable choice and is the focus of this work.

The FFS method can be used to calculate both the transition rate, *i.e.*, the inverse of the mean first passage time (*MFPT*), between two metastable states and the probability distribution along the transition path^35^. Although it can be applied to non-stationary processes^47^, here we investigate only stationary processes. The fundamental operation of FFS is to partition the phase space along an order parameter using a series of non-intersecting interfaces that track progress over the transition barrier. Many trial trajectories are started successively from each interface in order to compute the probability of advancing to the next interface versus returning to the initial state. The product of the advancement probabilities and the probability flux out of the initial state gives the mean transition rate.

The generation of many trial trajectories at each of the interfaces is a large part of the computational work in FFS. It is not surprising, then, that a number of authors have tried to optimize performance of FFS by studying the interplay between the number of trajectories sampled and the statistical error. Allen *et al.*^48^ first introduced a framework for studying the relationship between error in the estimated transition rate constant and error in the estimated interface trial probabilities. They also studied the computational efficiency of FFS, taken to be the inverse of cost times error, and showed that it was relatively insensitive to the choice of interface parameters. Borrero and Escobedo^49^ presented two techniques to minimize FFS error for a fixed cost either by optimizing the number of trial runs at each interface or by iteratively refining the placement of the interfaces. Kratzer *et al.*^50^ extended this idea with an algorithm to automatically define interface number and position on-the-fly by constraining the number of successful trials to be the same for each interface. However, none of these previous works included a systematic treatment of the error arising from the estimate of the flux out of the initial state and they all assume that computational cost per unit distance along the order parameter is fixed. Jian *et al.*^51^ later showed that interface placement along a complex transition landscape, *e.g.* one with a metastable intermediate state, cannot be correctly optimized by methods that assume cost is proportional to interface distance.

Here we present a method to optimally perform an FFS simulation of a nonequilibrium stationary process at a given margin of error and confidence interval. The key advance of our method is to estimate all of the interface probabilities and costs from a short pilot simulation and then use those parameters to optimize the configuration of a full FFS simulation. We minimize the total computational cost to achieve a user specified statistical error, which greatly reduces the number of choices that need to be made by the user. Our method includes a new formal treatment of the sampling error arising in the calculation of the flux out of the initial state, which is shown to have a significant influence on total cost. Unlike previous efforts, the cost in our method varies by interface, which enables it to account for changes in computational efficiency along the order parameter. We evaluate the capability of our method to control sampling error on three different models exhibiting rare event dynamics. In two one-dimensional models, sampling error dominates and is controlled precisely while in a multidimensional model landscape error becomes significant and requires oversampling. Finally, we derive an expression for the speedup of FFS relative to direct simulation for the same level of error and show that the advantage of FFS is substantial and increases with the rarity of the event.

The remainder of this work is organized as follows: In Sec II we introduce our theoretical framework for estimating the sampling error in a FFS simulation. In Secs III A and III B we derive the optimizing equation used by our method to control sampling error and then describe the forward flux pilot sampling (FFPilot) algorithm. In Secs III C-III E we evaluate the accuracy and performance of our method using three increasingly complex models. In Secs III F and III G we analyze sources of error outside of sampling error. Finally, in Secs III I and III J we evaluate the theoretical efficiency and measure the performance of FFPilot.

## II. THEORY AND METHODS

### A. Simulation of Rare Events in Stochastic Processes

The standard stochastic simulation protocol, here referred to as direct sampling (DS), proceeds in two iterated steps: 1) increment the system time with the time of the next stochastic event, 2) update the system state according to the event. These two steps are repeated until some predetermined stopping condition (simulation steps, simulation time, etc.) is reached, at which point the simulation is terminated. Repeated DS simulation produces a data set from which ensemble average quantities can be estimated by straightforward averaging. Running more DS replicates leads to increased accuracy.

As discussed above, DS has the disadvantage that it requires a long simulation time to sample each rare event. Therefore, calculating rare event statistics is computationally expensive. Enhanced sampling (ES) methods use a combination of constraints on simulation trajectories and statistical unbiasing methods in order to enrich the sampling of rare events. Forward flux sampling (FFS) is a popular ES method that was initially proposed by Allen and coworkers^33^. We implemented the FFS algorithm as follows:

1. Find the steady states of the model system, here termed 𝓐 and 𝓑. These will serve as an initial and a final state for the simulation.
2. Choose a one dimensional parameter 𝓞 for which 𝓞 𝓐 < 𝓞 𝓑.
3. Choose a set {λ_0_, · · ·, λ_N_ } of interface values of 𝓞 such that 𝓞 𝓐 *< λ_i_ < 𝓞* 𝓑 for every *λ_i_*. We say that a trajectory has fluxed forward with respect to a *λ_i_* when it crosses the interface traveling in the direction of increasing 𝓞, and that it has fluxed backward when it crosses traveling in the direction of decreasing 𝓞.
4. Begin FFS phase 0:
  a. Execute a DS trajectory initialized at *A* with a reflecting barrier at the midpoint between 𝓞 𝓐 and 𝓞 𝓑. Multiple phase 0 trajectories can be executed in parallel.
  b. Each time the trajectory fluxes forward across *λ*_0_, record the elapsed simulation time *τ* since the previous crossing event, and also record the state 𝓧.
  c. Once *n*_0_ samples of *τ* have been collected, terminate the trajectories and move to the next phase.
5. Begin FFS phase *i >* 0:
  a. Randomly choose a state *X* from the collection of states at which any trajectory in the previous phase crossed *λ_i−_*_1_.
  b. Execute a DS trajectory initialized at 𝓧. Allow the trajectory to run until it either crosses *λ*_0_ back into 𝓐 or moves forward across *λ_i_*. Terminate the trajectory and record a trajectory outcome value, either 0 or 1, depending on whether the trajectory moved backward or forward, respectively. If the trajectory moved forward, add the endpoint, which will lie along *λ_i_*, to the set of states that will be used to initialize trajectories during the next phase.
  c. Repeat steps 5a-b.
  d. Once *n_i_* trajectory outcomes have been collected terminate the phase. Every phase *i* trajectory can be executed in parallel.
6. The procedure for FFS phase *i* is then repeated for phase *i* + 1 (during which trajectories are launched from *λ_i_*) until the final phase, phase *N*, is reached. Trajectories in phase *N* begin at *λ_N−_*_1_ and may move forward across *λ_N_*and into 𝓑.

The overall aim of FFS is to ratchet a simulation from 𝓐 *→* 𝓑 across state space. The immediate goal of each phase is to estimate a phase weight. During phase *i*, samples are taken from an observable 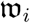 that behaves as a random variable. w_0_ is the waiting time in between forward flux events across *λ*_0_, and each 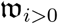 is the final outcome (0 for fall back or 1 for flux forward) of each trajectory launched in that *i*. The phase weight *w_i_* is defined as the true expected value of the random variable 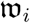:

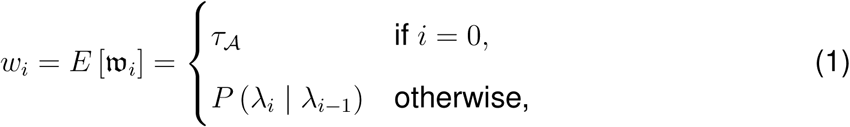

where *τ_A_* is the expected waiting time in between *λ*_0_ crossing events, and *P* (*λ_i_ | λ_i−_*_1_) is the probability that a trajectory launched from *λ_i−_*_1_ (*i.e.* launched during phase *i*) crosses forward past *λ_i_* before it falls back behind the starting interface *λ*_0_. The phase weights can be used to reweight the output of an FFS simulation so as to calculate unbiased statistics. Note that 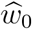 has different units that 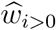, but we choose to discuss them all in terms of a

### B. Estimating Values of Interest from DS and ES Stochastic Simulations

As mentioned in the previous section, estimating values of interest from DS simulations is straightforward. One valid estimator of any ensemble average quantity is simply the arithmetic mean taken across an appropriate set of direct observations. For example, in order to calculate the *MFPT* of the switching process that takes some multistate system from 𝓐 *→* 𝓑, *n* DS simulations are initialized in 𝓐. Each of these *n* replicate simulations is allowed to run until the first time it enters 𝓑, at which point it is terminated. The time that each simulation *i* ran for is then a single sample *FPT_i_*of the first passage time of the switching process. The mean of these first passage time samples is an estimate of *MFPT* ^52^:

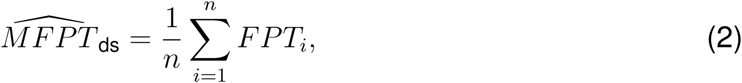

where [inline is the estimator (*i.e.* estimation function) of *MFPT* specific to DS simulation. Throughout this paper, we place a^ above a symbol to indicate that we are referring to an estimator of a value rather than to the value itself.

Equivalent ensemble average estimators can be calculated using FFS. Although the results of FFS simulations are biased and partitioned, statistical protocols allow FFS results to be recombined and reweighted so as to recapitulate the results of the unbiased ensemble. In order to perform these reweightings, we need estimates of the phase weights. During phase *i* of FFS, *n_i_* samples 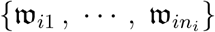 of an underlying random process 𝔴*_i_* are taken. The mean of this sample can be used as an estimator 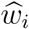 of phase weight *i*:

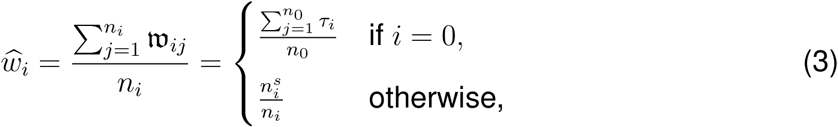

where each *τ_i_* is a sample of the waiting time in between phase 0 forward flux events and 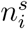 is the total count of trajectories that successfully fluxed forward during phase *i >* 0.

In an idealized situation in which it were possible to know the exact values of the phase weights *w_i_*, the exact value of the *MFPT* could be found via a simple combination *W* of the phase weights:

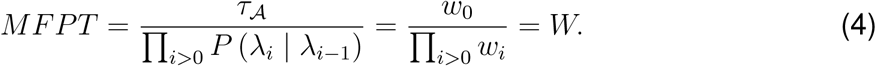

In reality, we only know the phase weight estimators 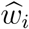, so we must instead estimate the value of the *MFPT* by way of the combination of estimators *W*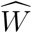

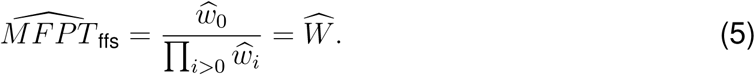

where 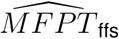 is the estimator of *MFPT* specific to FFS simulation. Other quantities of interest, such as the stationary PDF, can also be calculated using the phase weights^35^.

### C. Predicting Simulation Error in Terms of Margin of Error

Stochastic simulation results are not single valued, but are instead estimators, random variables with associated distributions. For example, when attempting to calculate *MFPT*, each complete round of simulations can be thought of as performing a single draw from the distribution of possible 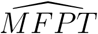 values. A prediction about the likely level of error in any given set of simulations can be made based on the characteristics of this estimate distribution. One way to quantify confidence in this prediction is in terms of the margin of error. The margin of error of a random variable *X* is defined as the ratio of half the width of a specified confidence interval and its expected value:

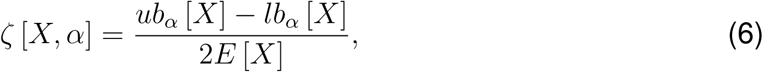

where *ζ* [*X, α*] is the margin of error of *X* at confidence level *α*, *E* [*X*] is the expected value of *X*, and *lb_α_* [*X*] and *ub_α_* [*X*] are the lower and upper confidence bounds, respectively.

For many values of interest *ζ* [*X, α*] is dependent on simulation parameters that are set by the user. For example, when calculating *MFPT*, the margin of error 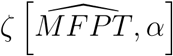 is dependent upon the total number of replicate simulations (when using DS), or upon the count of trajectories run in each phase (when using FFS).

### D. Determining Margin of Error from Simulation Parameters

It is straightforward to determine the margin of error of an estimator that depends on a single underlying observable. For example, consider the margin of error of an *MFPT* estimate as determined via DS simulation, 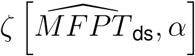. It has been shown that observations of first passage times between two well separated states follow an exponential distribution during DS simulations^21^. Given this distribution of first passage times, the central limit theorem^53^ gives the distribution of *MFPT* estimates in the limit of large sample size:

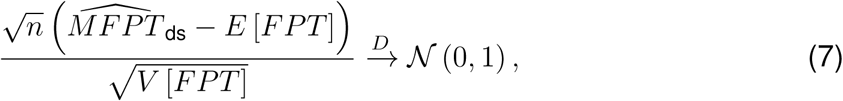

where *n* is the count of first passage time observations, *E* [*FPT*] = *MFPT* is the expected value of the first passage time, *V* [*FPT*] is the variance, 𝓝 (0, 1) is the standard normal distribution (*i.e.* mean 0 and variance 1), and 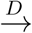 signifies that the distribution on the left converges to the one on the right. Eq 7 implies^53^ that an estimate of *MFPT* calculated using *n* observations will follow a normal distribution such that:

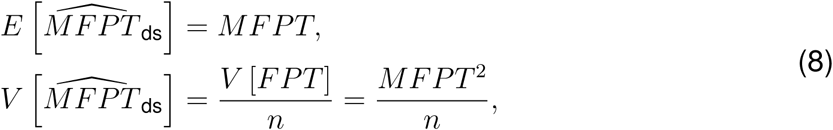

where the fact that *V* = *E*^2^ for an exponential distribution was used to factor out *V* [*FPT*]. The lower and upper bounds of the confidence interval of any normally distributed random variable *X* can be found using a standard formula^54^ that depends on the first two moments of *X*:

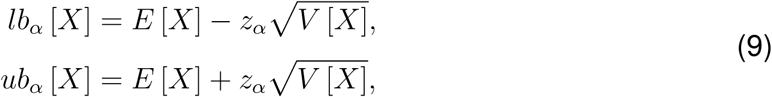

where *z_α_* is the z score associated with the confidence level *α* (*e.g. z.*_95_ *≈* 1.96)^55^. Plugging Eqs 8 and 9 into Eq 6 yields the margin of error of the DS *MFPT* estimator:

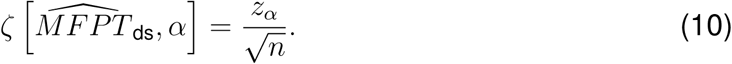

Thus, if say, 10^5^ replicate trajectories are produced during a DS simulation, the resultant estimate of *MFPT* will have no more than value 95% of the time. 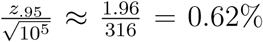 difference from the true

### E. Minimizing the Computational Cost Required to Achieve a Desired Error Goal

We define the computational cost 𝓒 of a simulation to be equivalent to the average in-simulation time required to complete it (alternatively, one may use the count of simulation steps). When estimating *MFPT* using DS simulation, the computational cost is:

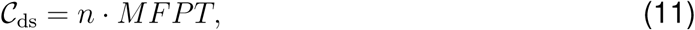

where *n* is the total number of replicate trajectories. Eq 10 illustrates the direct trade-off between computational cost and simulation accuracy. In order to find the minimum number of replicate trajectories that are required to achieve a particular error goal in DS simulations, Eq 10 can be solved for n:

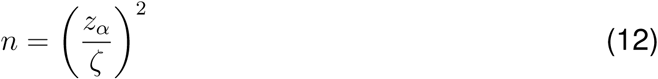

## III. RESULTS

### A. Derivation of the FFPilot optimizing equation

In this subsection we find the number of trajectories to launch in each FFS phase (which we will also refer to as sample count or *n_i_*) that will minimize simulation run time while fixing the margin of error *ζ* of the estimator of *MFPT*. We will refer to the *MFPT* estimator in this subsection as 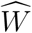. The derivation of the optimal choice of *n_i_* starts with the characterization of the moments and distribution of 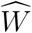. We then use the moments and distribution of 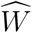 to find *ζ* as a function of the per-phase sample counts *n_i_*. We then use the method of Lagrange multipliers to find the optimal choice of *n_i_* that will keep *ζ* fixed while minimizing simulation run time.

#### 1. The Moments and Distribution of the FFS *MFPT* Estimator 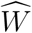

Each individual FFS phase weight *w_i_* can be thought of as the first moment (*i.e.* the mean) of an observable 𝔴*_i_* of a random process that is specific to phase *i*. By the end of every FFS phase *i*, *n_i_* samples have been drawn from 𝔴*_i_* (see Sec II A for a more concrete description), at which point the phase weight *w_i_* is estimated as:

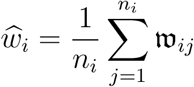

Given the above form of 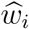, and given that the observable w meets certain regularity conditions^56^, the asymptotic distribution of the individual 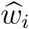 terms can be determined from the central limit theorem:

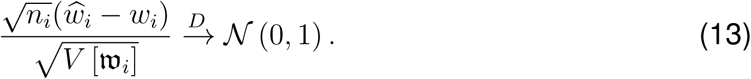

The moments of each 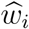 can be determined by the appropriate interpretation of Eq 13:

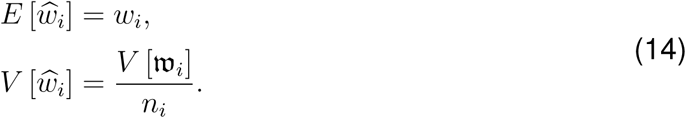

An estimator with this type of convergence behavior is said to be 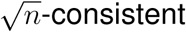.

Given that the *MFPT* estimator 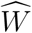 is defined in Eq 5 as a function 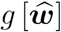 of a vector of 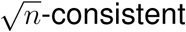 estimators 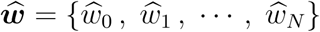, we can apply the multivariate delta method^57^ to determine the moments and distribution of 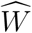. From the multivariate delta method, we know that:

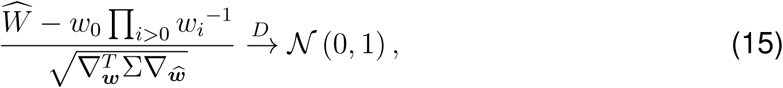

where ∇***_w_*** is the gradient of 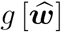 evaluated at 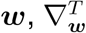 is the transpose of ∇***_w_***, and Σ is the covariance matrix of the phase weight estimators 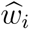. We can determine the moments of 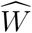 by interpreting Eq 15:

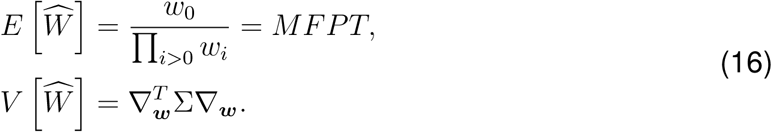

A simpler form of 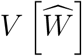 can be found. To begin with, we find ∇***_w_***. In column vector form it is:

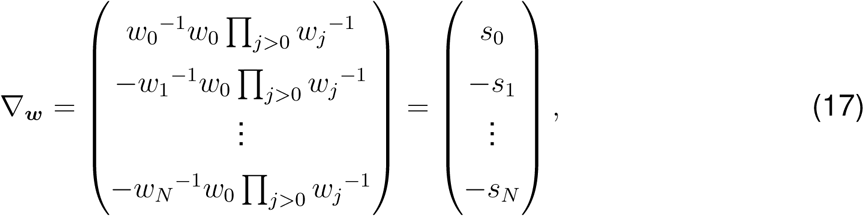

where we have substituted *s_i_* for 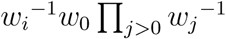 for the sake of brevity. If the covariance of 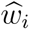 and 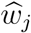 is written as *σ_ij_*, then the covariance matrix Σ is:

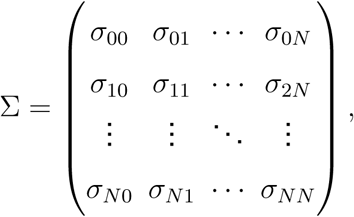

and the variance of 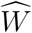 can be written as:

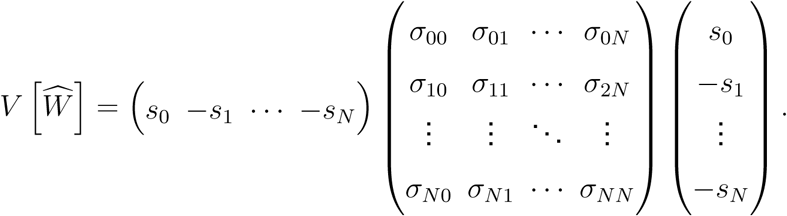

If we impose the assumption of independence on all of the phase weight estimators 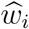, then only the diagonal elements of the covariance are non-zero:

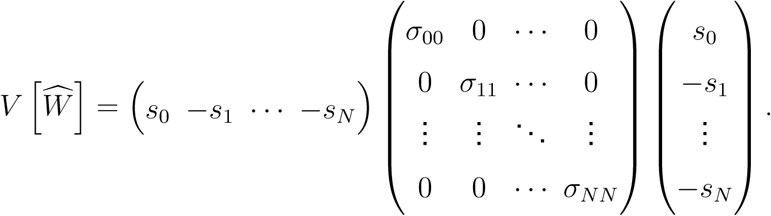

Under this condition of independence the form of 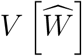 can be simplified considerably:

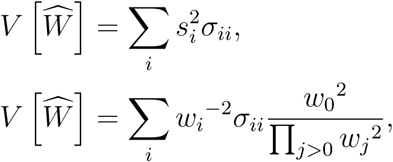

and since 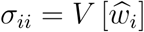:

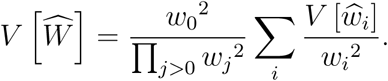

Finally, plugging in substitutions from Eq 4 and from Eq 14, the variance of 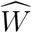 is:

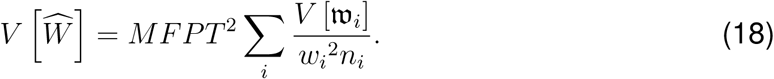

Alternatively, the variance of 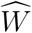 can be derived from the formula for the variance of a product of random variables (see supplemental text). The expressions derived from each technique agree in the high sample count limit.

The result in Eq 18 agrees with and is similar to the established result of Allen *et al.*^48^ concerning the variance of estimates produced by a complete FFS simulation. Unlike earlier work, however, we have imposed no particular form on 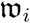, the random process underlying each phase *i*, *i.e.*, we do not assume here that it is a Bernoulli process. As will be seen in Sec III A 4, this generalization allows us to study the contributions of phase 0 to the overall error of an FFS simulation for the first time.

#### 2. Margin of Error of 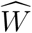

Now we derive a formula for 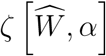, the margin of error of the *MFPT* estimator 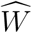. From Eq 15 we know that 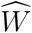 follows a normal distribution. The lower and upper confidence bounds of 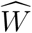 are found by plugging the moments of 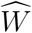 (given by Eqs 16 and 18) into the bounds formulas for a normally distributed random variable (given by Eq 9):

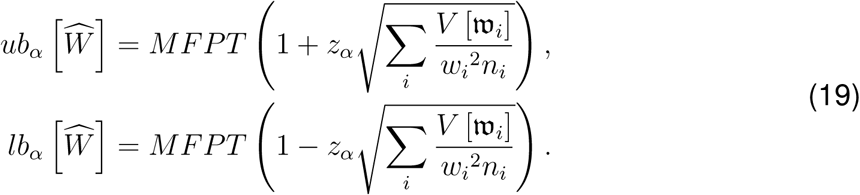

Plugging Eq 19 into the margin of error definition (given by Eq 6) yields the desired margin of error:

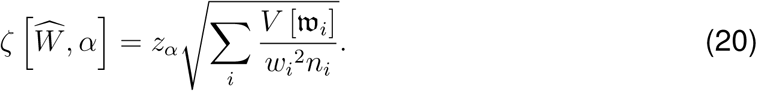

#### 3. Derivation of the General Optimizing Equation

As we can see from Eq 20, there are many different choices of *n_i_* that will give the same value of 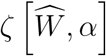. What we really want to find is the optimal choice of *n_i_* that will minimize simulation run time while keeping 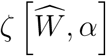 fixed. We can find a formula for this optimal choice using the method of Lagrange multipliers^58^.

For the method of Lagrange multipliers, we need a function to minimize, the target function *f* [*x*], and a function to hold constant, the constraint equation *g*[*x*]. We use the total computational cost 𝓒 as the target function, which for FFS is:

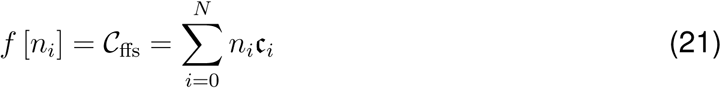

where c*_i_* is the average computational cost per sample. For the constraint equation, we square both sides of Eq 20 and set it equal to zero:

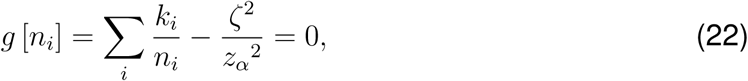

where

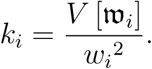

Now that we’ve chosen a target and a constraint function, the next step of the method is to combine Eqs 21 and 22 in order to write out the Lagrangian:

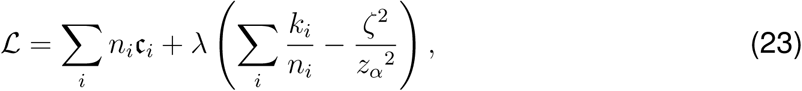

and then find its gradient ∇𝓛:

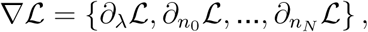

We find the values of each of the partial derivatives individually. Calculating *∂_λ_*𝓛 is trivial, and when finding each separate *∂_n_i__* 𝓛 we can eliminate all but one term from both sums, since terms that don’t depend on *n_i_* will vanish:

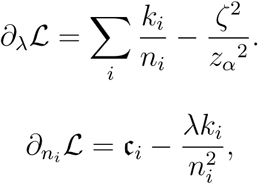

Now we can write out the actual gradient

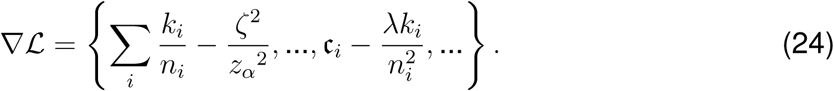

The next step is to set each component of the gradient Eq 24 equal to zero and solve the resulting set of *N* + 2 equations for *λ* and each *n_i_*. We begin by solving *∂_ni_* 𝓛 = 0 for *n_i_* in terms of *λ*:

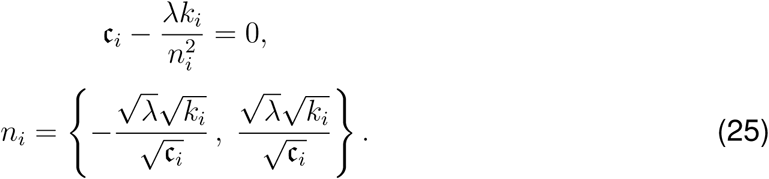

Next, we solve *∂_λ_*𝓛 = 0 for 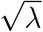 by substituting in the positive expression for *n_i_* found in Eq 25:

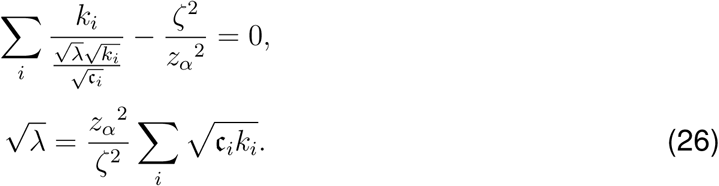

Now we eliminate *λ* from our expression for *n_i_* in Eq 25 by substituting in the expression for 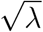 we found in Eq 26:

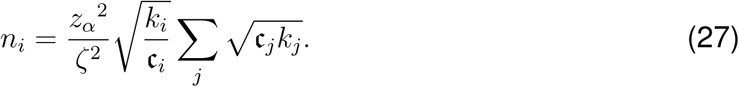

#### 4. The Optimizing Equation for FFS

A form of the general optimizing equation given in Eq 27 that is more specific to FFS can be found by considering the properties of the observable 𝔴*_i_* in each phase. During a phase *i >* 0, each trajectory launched and finished is equivalent to a single sample taken from 𝔴*_i>_*_0_. The *j*th trajectory of a phase will either succeed (*i.e.* cross forward to the next interface) with probability *p_ij_*, or it will fail (*i.e.* fall back into its starting basin), with probability 1 *− p_ij_*. The sample taken from 𝔴*_i>_*_0_ is 1 If the trajectory succeeds, and 0 otherwise. Thus, the outcome of the *j*th trajectory of phase *i* is a Bernoulli random variable with probability parameter *p_ij_*.

For multidimensional systems, *p_ij_* is dependent upon the starting point of a trajectory. Since the starting point of each trajectory is chosen at random, *p_ij_* in general varies from trajectory to trajectory. Thus, each 𝔴*_i>_*_0_ is technically a mixture of Bernoulli random variables. However, for the purposes of the optimizing equation, it can be shown that no accuracy is lost if each 𝔴*_i>_*_0_ is treated as a single Bernoulli random variable (see Appendix A) with moments:

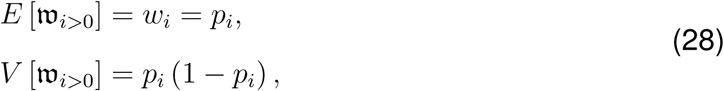

where *p_i_* is the crossing probability in phase i (*i.e. P* (*λ_i_ | λ_i−_*_1_)).

The *k_i_* terms in Eq 27 can be expanded to yield:

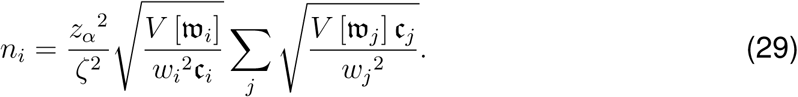

The moments from Eq 28 can then be plugged into Eq 29 to yield the FFS specific form of the optimizing equation:

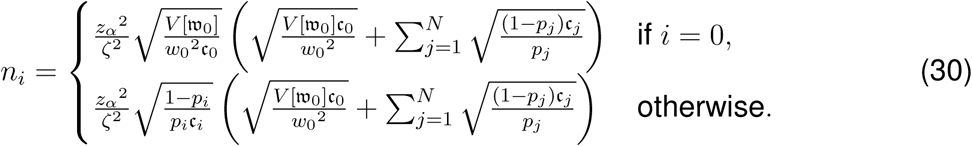

We call this form the FFPilot optimizing equation.

In regards to phase 0, the precise forms of 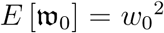 and 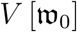 are unknown. We have found that 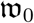, the waiting time in between phase 0 forward flux events, does not in general follow an exponential distribution (see supplemental Fig S2). In fact, the distribution of 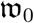 seems to be highly model dependent. Thus, in order to avoid any assumptions about w_0_, we leave the ratio 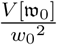 unexpanded in Eq 30.

Of the assumptions made in deriving Eq 30, two of the most significant are the assumption of large sample size, and the assumption of the uncorrelatedness of the phases during an FFS simulation. The large sample size assumption underlies the validity of Eqs 14 and 15. In general this assumption can be satisfied by setting a minimum floor on the number of samples *n_i_* taken in each phase (an implementation of this sample size floor is discussed in the next section).

Ensuring that the uncorrelatedness assumption is satisfied is altogether trickier. Most of the preexisting FFS literature takes the uncorrelatedness of the phases as a given^47–49^, but in practice we have found this to not always be the case. This issue is discussed in detail in Sec III F. In brief, systems with complex, high dimensional state spaces tend to have correlations between the outcomes of trajectories across the different phases. This results in non-zero covariance between the different phase weights, which is effectively like adding an extra term to Eq 18. In other words, when the phases are correlated, our approach will somewhat underestimate the actual variance, and Eq 30 will somewhat underestimate the number of samples required to achieve a particular error goal.

##### B. FFPilot: A Sampling Algorithm Designed to Take Advantage of the Optimization Equation

We wanted to be able to apply the optimizing equation to real biochemical networks, but to do so we need some prior knowledge of the system under study. As shown in Eq 30, two global and 2 *·* (*N* + 1) phase-specific parameters are required in order to apply the optimization equation and thereby calculate the optimal value of *n_i_*. The two global parameters, the margin of error *ζ* and the confidence interval z score *z_α_*, are independent of the model being studied and are set according to the desired error goal. The other parameters, the ratio 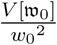 (from phase 0), the successful crossing probabilities *p_i_* (from phases *i >* 0), and the per-sample computational costs c*_i_*, are model dependent and vary for each different combination of model, order parameter, and interface placement.

In general the exact values of the model-dependent parameters are unknown and estimates must be used instead. Rather than simplifying using assumptions such as constant cost^48, 49^, we produce rough but conservative estimates of the necessary parameters using a pilot simulation. By conservative, we mean that the estimates, when plugged into the optimization equation, will be likely to give values of *n_i_* that are at least as large as the true values. This condition ensures that simulations run with *n_i_* trajectories per phase will produce results that are at least as accurate as the specified error goal.

We call our new enhanced sampling protocol FFPilot. The basic concept of FFPilot is to break an FFS simulation up into two stages. First a pilot stage is executed (see supplemental Fig S1), from which the parameters required for the optimization equation are estimated. Then, based on the results of the optimization equation, a production stage is planned and executed, from which the actual simulation output is calculated.

The FFPilot algorithm proceeds as follows:

1. Specify an error goal in terms of a target margin of error. Optionally, the confidence interval (which defaults to 95%) can be specified as well. As with standard FFS, the user must specify an order parameter and interface placements.
2. Begin FFPilot pilot stage:
  a. Set the pilot stage sample count *n*_pilot_ to a single fixed value. Throughout this paper we used a value of *n*_pilot_ = 10^4^.
  b. Run a complete FFS simulation, following the algorithm described in Sec II A. Unlike standard FFS, the number of samples to collect in each phase is determined by a blind optimization method.
    i. Phase 0 proceeds the same as in standard FFS, using *n*_0_ = *n*_pilot_ as the sample count.
    ii. In phases *i >* 0, trajectories are run until *n*_pilot_ successful forward flux events are observed. It can be shown that, for a relatively modest number of successes, this method constrains error to within 2% when estimating the individual phase weights (see Appendix B for complete details).
3. When the pilot stage is finished, estimate the values required for the optimization equation from the results of the pilot simulation. Use confidence intervals to form conservative estimates that, when plugged into the optimization equation, are likely to yield values of *n_i_* that are as large or larger than those required for the error goal.
4. Begin FFPilot production stage:
  a. Determine *n_i_*, the number of samples to collect in each phase, based on the error goal and Eq 30, the FFPilot optimizing equation (as parameterized in step 3).
  b. Run another complete FFS simulation using using the values of *n_i_* calculated in step 4a.
5. Collect results from the production stage simulation and use/analyze them in the same way as would be done for standard FFS simulation. The results from the pilot stage are ignored for the purposes of calculating the final simulation results as sampling differences in states at the interfaces would introduce additional error.

In terms of the error in the final simulation results, the optimization equation is inaccurate in the low sample count limit. Therefore, we enforce a minimum floor (10^3^) on the count of samples taken in each phase of the production stage.

The effectiveness of the pilot stage blind optimization method can be related to earlier findings of Glasserman *et al.*^59^ and Borerro *et al.*^49^. They showed that a fixed quantity of computational effort is optimally spent during an FFS simulation when the interfaces and the trajectory counts per phase are arranged in such a way that each interface encounters an equal flux of trajectories crossing them. Although we do not constrain the computational effort spent during our blind optimization, our approach produces equal flux across each interface as well.

##### C. Rare Event Model

We began our testing of FFPilot with a toy model of a barrier crossing process, which we refer to as the rare event model (REM). REM models a particle in a discrete potential field in which there are two metastable states, 𝓐 and 𝓑, connected by a transition path (see top of Fig 1). Particles in 𝓐 have a constant propensity to initiate a transition by entering the transition path. Because of the constant propensity the waiting times in between transition attempts are exponentially distributed.

**FIG. 1.**
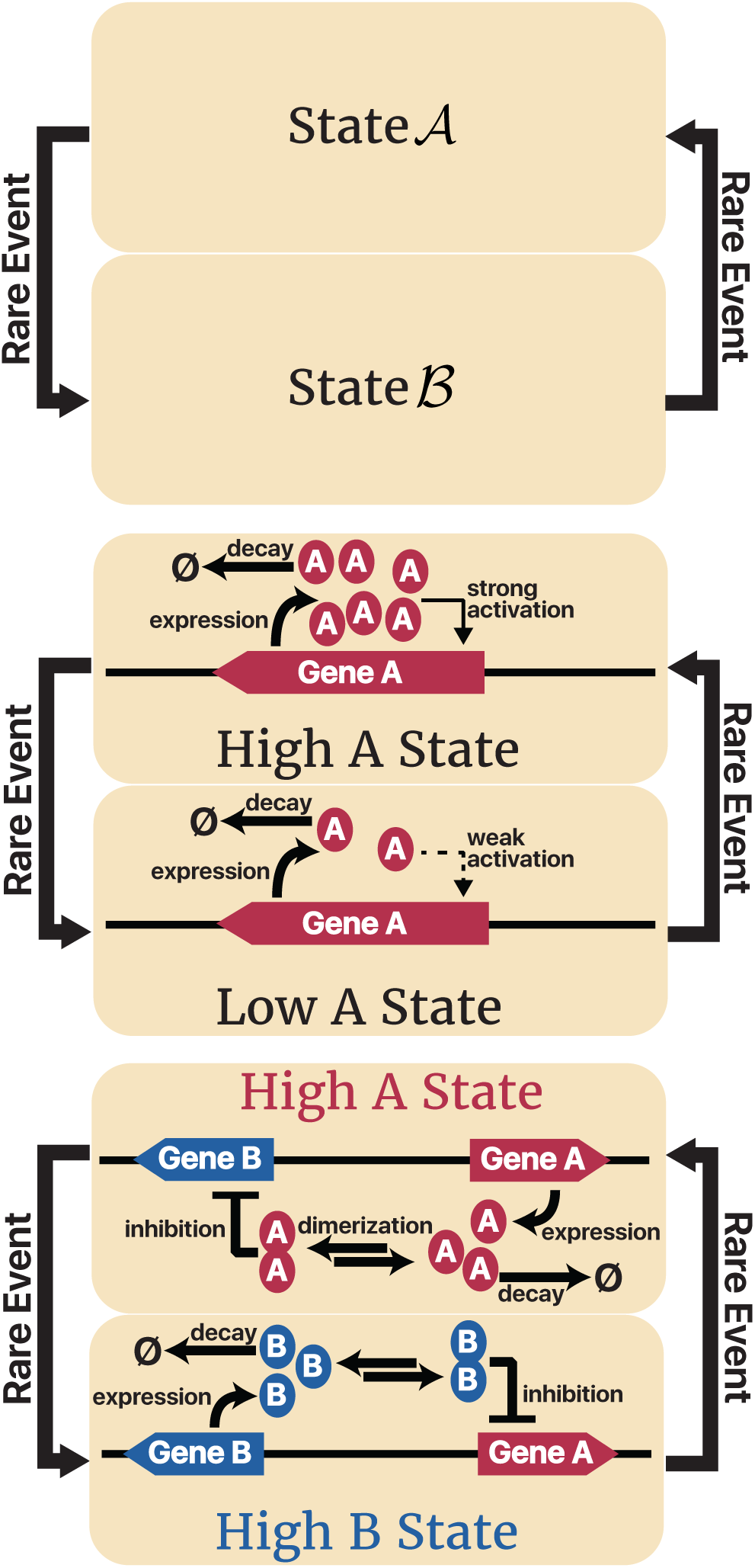
Schematics of the model systems investigated. (top) The Rare Event Model (REM, see Sec III C and Table I for complete details). (middle) Self Regulating Gene (SRG, see Sec III D and Tables II and III). (bottom) Genetic Toggle Switch (GTS, see Sec III E and supplemental Table S1.

**TABLE I.** Parameterization and other simulation data from our simplified Rare Event Model (REM). The phase weights wi were copied from those of GTSθ=1 (see Table V). All sample costs c were set to 1.

| phase | <sup>a</sup>$w_i$ | <sup>b</sup>$\mathbf{c}_i^f$ | <sup>c</sup>$\mathbf{c}_i^s$ | <sup>d</sup>$\mathbf{c}_i$ | <sup>e</sup>$n_i \times 10^{-3}$ |
| --- | --- | --- | --- | --- | --- |
| 0 | 45.3 |  |  | 1 | 1834 |
| 1 | .092 | 1 | 1 | 1 | 2020 |
| 2 | .274 | 1 | 1 | 1 | 1047 |
| 3 | .136 | 1 | 1 | 1 | 1621 |
| 4 | .150 | 1 | 1 | 1 | 1527 |
| 5 | .242 | 1 | 1 | 1 | 1138 |
| 6 | .636 | 1 | 1 | 1 | 486 |
| 7 | .711 | 1 | 1 | 1 | 410 |
| 8 | .858 | 1 | 1 | 1 | 261 |
| 9 | .913 | 1 | 1 | 1 | 199 |
| 10 | .973 | 1 | 1 | 1 | 107 |
| 11 | .989 | 1 | 1 | 1 | 67 |
| 12 | .998 | 1 | 1 | 1 | 30 |
| MFPT | $1.074 \cdot 10^6$ | | | | |

<sup>a</sup> The phase weight of phase $i$ .
<sup>b</sup> The expected cost of a failed phase $i$ trajectory.
<sup>c</sup> The expected cost of a successful phase $i$ trajectory.
<sup>d</sup> The expected cost of a phase $i$ sample.
<sup>e</sup> The phase $i$ sample count (in thousands) for a 1% error goal with 95% confidence.

**TABLE II.** Reaction scheme for our Self Regulating Gene (SRG) models.

| Reactions | Rates | Description |
| --- | --- | --- |
| $\emptyset \longrightarrow \mathbf{A}$ | $k_{low} + (k_{high} - k_{low}) \frac{\mathbf{A}^h}{k_{50}^h + \mathbf{A}^h}$ | expression |
| $\mathbf{A} \longrightarrow \emptyset$ | $\beta \mathbf{A}$ | decay |

**TABLE III.** Parameterizations and other simulation data from our SRG models. Per-phase weights, costs, and sample counts were estimated from the average results of 1000 FFPilot simulations (1% error goal). *MFPT* values were estimated from the results of 10^5^ DS simulations, plus or minus the 95% confidence interval bounds.

| parameter | SRG<sub>h=2.4</sub> |  |  |  |  |  | SRG<sub>h=2.3</sub> |  |  |  |  |  | SRG<sub>h=2.2</sub> |  |  |  |  |  |
| --- | --- | --- | --- | --- | --- | --- | --- | --- | --- | --- | --- | --- | --- | --- | --- | --- | --- | --- |
| $h$ | 2.4 | | | | | | 2.3 | | | | | | 2.2 | | | | | |
| $k_{low}$ | 10 | | | | | | 10 | | | | | | 10 | | | | | |
| $k_{high}$ | 200 | | | | | | 200 | | | | | | 200 | | | | | |
| $k_{50}$ | 91 | | | | | | 92.5 | | | | | | 94 | | | | | |
| $\beta$ | 1 | | | | | | 1 | | | | | | 1 | | | | | |
| $\mathcal{A}$ | 12 | | | | | | 12 | | | | | | 11 | | | | | |
| $\mathcal{B}$ | 155 | | | | | | 156 | | | | | | 157 | | | | | |
| phase | <sup>a</sup>$\lambda_i$ | <sup>b</sup>$w_i$ | <sup>c</sup>$c_i^f$ | <sup>d</sup>$c_i^s$ | <sup>e</sup>$c_i$ | <sup>f</sup>$n_i \times 10^{-3}$ | $\lambda_i$ | $w_i$ | $c_i^f$ | $c_i^s$ | $c_i$ | $n_i \times 10^{-3}$ | $\lambda_i$ | $w_i$ | $c_i^f$ | $c_i^s$ | $c_i$ | $n_i \times 10^{-3}$ |
| 0 | 23.0 | 9.66 |  |  | 9.66 | 641 | 23.0 | 6.89 |  |  | 6.89 | 703 | 23.0 | 4.78 |  |  | 4.78 | 762 |
| 1 | 33.6 | .007 | .130 | .913 | .135 | 26395 | 34.1 | .010 | .137 | .936 | .150 | 18975 | 34.6 | .018 | .138 | .846 | .151 | 12067 |
| 2 | 44.2 | .071 | 1.56 | 1.21 | 1.53 | 2383 | 45.2 | .112 | 1.65 | 1.20 | 1.60 | 1620 | 46.2 | .140 | 1.64 | 1.30 | 1.60 | 1253 |
| 3 | 54.8 | .277 | 2.89 | 1.14 | 2.41 | 850 | 56.2 | .296 | 2.99 | 1.24 | 2.47 | 714 | 57.8 | .354 | 3.05 | 1.23 | 2.41 | 556 |
| 4 | 65.3 | .565 | 4.22 | 1.30 | 2.57 | 447 | 67.3 | .580 | 4.35 | 1.28 | 2.57 | 387 | 69.3 | .572 | 4.45 | 1.39 | 2.70 | 337 |
| 5 | 75.9 | .855 | 5.45 | 1.12 | 1.75 | 254 | 78.4 | .825 | 5.63 | 1.23 | 2.00 | 237 | 80.9 | .814 | 5.78 | 1.23 | 2.08 | 212 |
| 6 | 86.5 | .962 | 6.48 | 1.03 | 1.24 | 147 | 89.5 | .946 | 6.75 | 1.08 | 1.39 | 147 | 92.5 | .924 | 7.01 | 1.24 | 1.67 | 141 |
| 7 | 97.1 | .993 | 7.35 | .844 | .890 | 73 | 101 | .986 | 7.71 | .926 | 1.02 | 85 | 104 | .977 | 8.12 | 1.09 | 1.25 | 88 |
| 8 | 108 | .999 | 8.11 | .700 | .709 | 34 | 112 | .997 | 8.56 | .836 | .862 | 46 | 116 | .993 | 9.07 | .905 | .959 | 53 |
| 9 | 118 | 1.00 | 8.83 | .767 | .769 | 16 | 123 | .999 | 9.42 | .825 | .833 | 25 | 127 | .997 | 10.0 | .981 | 1.00 | 32 |
| 10 | 129 | 1.00 |  | .779 | .779 | 8 | 134 | 1.00 | 10.4 | .894 | .897 | 14 | 139 | .999 | 11.0 | .955 | .965 | 20 |
| 11 | 139 | 1.00 |  | 1.05 | 1.05 | 5 | 145 | 1.00 |  | 1.07 | 1.07 | 9 | 150 | .999 | 12.3 | 1.24 | 1.25 | 14 |
| 12 | 150 | 1.00 |  | 1.33 | 1.33 | 3 | 156 | 1.00 |  | 1.44 | 1.44 | 6 | 162 | 1.00 | 13.8 | 1.48 | 1.49 | 10 |
| MFPT | $(1.521 \pm .0094) \cdot 10^5$ | | | | | | $(4.661 \pm .029) \cdot 10^4$ | | | | | | $(1.271 \pm .0079) \cdot 10^4$ | | | | | |

<sup>a</sup> The position of interface $i$ in terms of order parameter $A$ .
<sup>b</sup> The phase weight of phase $i$ .
<sup>c</sup> The expected cost of a failed phase $i$ trajectory.
<sup>d</sup> The expected cost of a successful phase $i$ trajectory.
<sup>e</sup> The expected cost of a phase $i$ sample.
<sup>f</sup> The phase $i$ sample count (in thousands) for a 1% error goal with 95% confidence.

The transition path itself is composed of a sequence of *N* barriers; particles enter the path before barrier 1. At each successive barrier, a particle will instantaneously either proceed to the next barrier with probability *p_i_*, or fall back into 𝓐 with probability 1 *− p_i_*. If the particle successfully passes the final barrier it enters 𝓑. In effect, the particle’s fate once it enters the transition path can be thought of as the outcome of *N* weighted coin flips. If all *N* coins land heads up, the particle completes the transition 𝓐 *→* 𝓑. Otherwise, the particle falls back into *A*.

We designed REM to map precisely onto FFPilot sampling in order to directly test the validity of the assumptions and simplifications that were made in the derivation of the FFPilot optimizing equation (Eq 27). Simulations of REM can be carried out using either DS or FFPilot. To simulate a single replicate of a particle starting in 𝓐 using DS, first the time until the particle leaves 𝓐 is randomly selected from an exponential random variable according to the propensity of entering the transition path. The particle’s behavior at each barrier is then randomly chosen, either falling back to 𝓐 or proceeding to the next barrier according to the appropriate *p_i_*. If the particle falls back into 𝓐 the process is repeated until it successfully passes to 𝓑. The total time the particle took to transition to 𝓑 is the 𝓐 *→* 𝓑 first passage time for that trajectory.

To simulate a single replicate of a particle starting in 𝓐 using FFPilot, first an order parameter and the interface positions must be chosen (see section Sec II A). We chose the barrier number *i* as the order parameter, and placed the interfaces between each barrier *i*. The phase 0 weight is calculated by first drawing many samples of the *A* leaving time according to the propensity, and then taking the mean of those samples. The remaining phase weights are determined by repeatedly starting a particle at barrier *i*, randomly selecting if it continues on to the next barrier according to *p_i_*, and then calculating the average probability of success from the observations. The pilot stage of FFPilot is accomplished by first running the phase 0 weight calculation *n*_pilot_ times, then running each phase *i >* 0 weight calculation until *n*_pilot_ success events are observed. The outcome of the pilot stage is then fed into the FFPilot optimizing equation Eq 30, and the results are used to determine how many samples to take during the FFPilot production stage. For the purposes of parameterizing Eq 30, the phase 0 relative variance and the per-phase costs c*_i_*are all set equal to 1 (see Table I for complete parameters). *MFPT* is then estimated as the product of the production stage phase weights.

We first studied the distribution of the *MFPT* estimators. Taken together, Eqs 15, 16, and 18 describe the normal distribution that repeated estimation of *MFPT* is expected to produce. In our derivation we have assumed that we are working in the high sampling limit for values of *w_i_* and *n_i_* of interest. To test this assumption, we performed 1.6 *·* 10^5^ independent simulations of REM using both DS and FFPilot using a 1% error goal. Fig 2 shows the distributions from our simulations. The black line in each figure is the normal distribution with mean and variance given by Eqs 16 and 18. The binned *MFPT* estimates from both the DS and the FFPilot REM simulations are in excellent agreement with the predicted normal distribution.

**FIG. 2.**
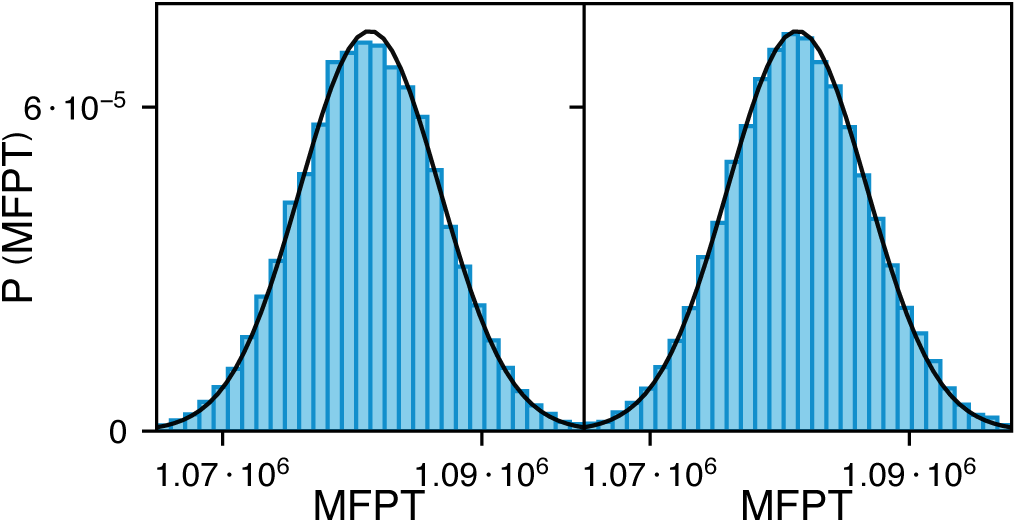
*MFPT* estimates for REM calculated using (left) DS and (right) FFPilot. Shown are (blue bars) binned *MFPT* estimates from 1.6 *·* 10^5^ simulations with a 1% error goal and (black lines) *MFPT* estimate distributions from Eq 7 and Eq 15, respectively.

Next, we tested how well the FFPilot approach was able to control sampling error in simulations of REM. If the method works as expected, 95% of simulations should have errors at or below the FFPilot error goal. We executed 1000 FFPilot simulations of REM at 3 different error goals (10%, 3.2%, and 1%). We used the full FFPilot algorithm to determine how many trajectories to start at each interface.

The percent errors of the *MFPT* calculated in each of these simulations are shown in Fig 3. The percent errors were calculated relative to the analytically determined *MFPT*. As can be seen, the 95th error percentiles (marked by the red lines) are located along *x* = *y*, indicating that overall error in the *MFPT* estimates was constrained to the error goal. REM has no source of error aside from sampling error, and under these conditions FFPilot precisely controls the total simulation error.

**FIG. 3.**
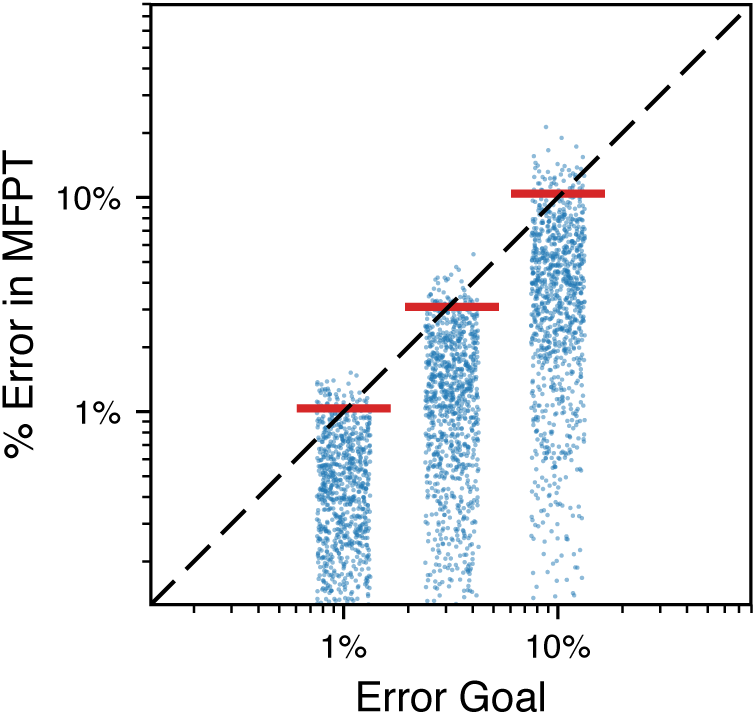
Actual error vs error goal of FFPilot simulations of the REM. Each strip shows the *MFPT* estimation error from (blue dots) 1000 independent FFPilot simulations and (red lines) the 95th percentiles of the errors. Jitter was added to the x-position of the dots for visualization.

##### D. Self Regulating Gene Model

We next tested FFPilot with a relatively simple biochemical network, the self regulating gene model (SRG)^6^. SRG models expression of a single protein *A*. *A* is produced though autocatalysis, and decays via a first order process (see Fig 1).

In the deterministic formulation of SRG, the rate of change in the quantity of protein *A* is:

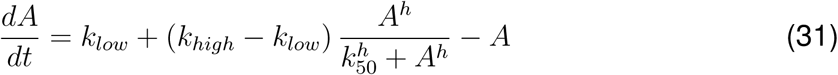

For a given set of parameters, the fixed points of the state space of SRG can be found by setting Eq 31 equal to 0 and solving for *A*. For all of the parameters we used in our simulations there are three fixed points, two stable and one unstable. One of the stable fixed points corresponds to a state with a low count of *A*, and the other corresponds to a state with high count of *A*.

To formulate SRG as a stochastic model, we use the chemical master equation (CME) (see Table II for complete reaction list). The CME models the probability for the system to be in any state. Additionally, fluctuations due to population noise can cause the system to transition back and forth between the low and high states. The *MFPT* between the states is related to the entropic barrier separating them.

We wanted to study how the height of the barrier between the low *A* and high *A* states affects the accuracy of FFPilot. Towards this end, we parameterized 3 different variants of SRG with different barrier heights, and thus different *MFPT* values. We call these three variants SRG_h=2.4_, SRG_h=2.3_, and SRG_h=2.2_, after the Hill coefficient used in the protein *A* production rate law. We tuned the other parameters in the model in order to approximately balance the occupancy of the low *A* and high *A* states in each of the variants (see Table III for parameter values).

For all FFPilot simulations of SRG we used the count of protein *A* as the order parameter. We determined the positions of the interfaces by first placing *λ*_0_ a quarter of the distance (in terms of the order parameter) from the lower fixed point to the intermediate fixed point, then *λ_N_* three quarters of the distance from the lower fixed point to the upper fixed point. We then placed 11 more interfaces spaced evenly between *λ*_0_ and *λ_N_*.

Unlike REM, trajectories in the low *A* state do not cross *λ*_0_ and enter the transition pathway with a fixed propensity. Instead, the propensity changes dynamically with the system state, giving rise to a complex distribution of waiting times in between crossing events. In deriving the FFPilot optimizing equation (more specifically, when deriving Eq 18), we assumed that the first two central moments of the phase 0 waiting time distribution (*w*_0_ and 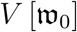) exist. In order to test this assumption we executed simulations in which we only performed phase 0, collecting 10^6^ crossing events for *λ*_0_.

Fig 4 shows the phase 0 inter-event time distributions. The tail of each distribution is fit well by a single exponential distribution, but the distribution near 0 is not. We estimated the value of the relative variance, 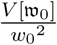, used in the phase 0 terms of the FFPilot optimizing equation (Eq 30) to be 6.80, 6.43, and 5.96 for SRG_h=2.4_, SRG_h=2.3_, and SRG_h=2.2_, respectively.

Next, we examined how well the FFPilot approach was able to control sampling error with respect to *MFPT*. We executed 1000 FFPilot simulations of SRG_h=2.4_, SRG_h=2.3_, and SRG_h=2.2_ using error goals 1%, 3.2%, and 10%. We estimated *MFPT* from each simulation, then found the percent error relative to the value estimated from a DS simulation executed with an error goal of 0.62%.

The errors are shown in Fig 5. The 95th error percentiles are again located precisely along *x* = *y*. The accuracy of the estimates show that FFPilot is able to control error in both phase 0 (regardless of the exact distribution of the inter-event times) and the remaining phases for SRG.

**FIG. 4.**
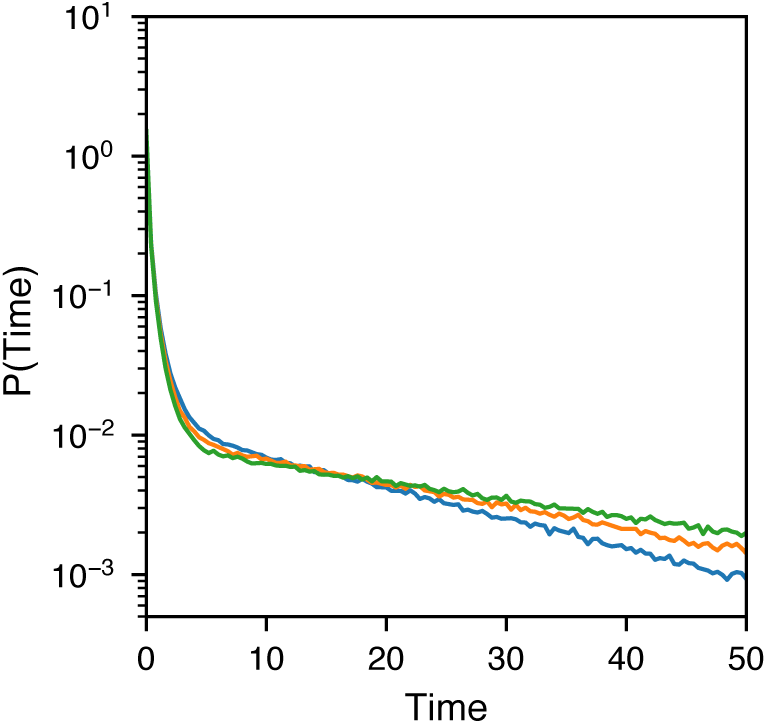
Waiting times between forward crossing events during phase 0 of forward flux simulations of the SRG models. Each line shows a distribution calculated from 10^6^ samples from a single phase 0 trajectory of (blue) SRG_h=2.4_, (orange) SRG_h=2.3_, and (green) SRG_h=2.2_.

**FIG. 5.**
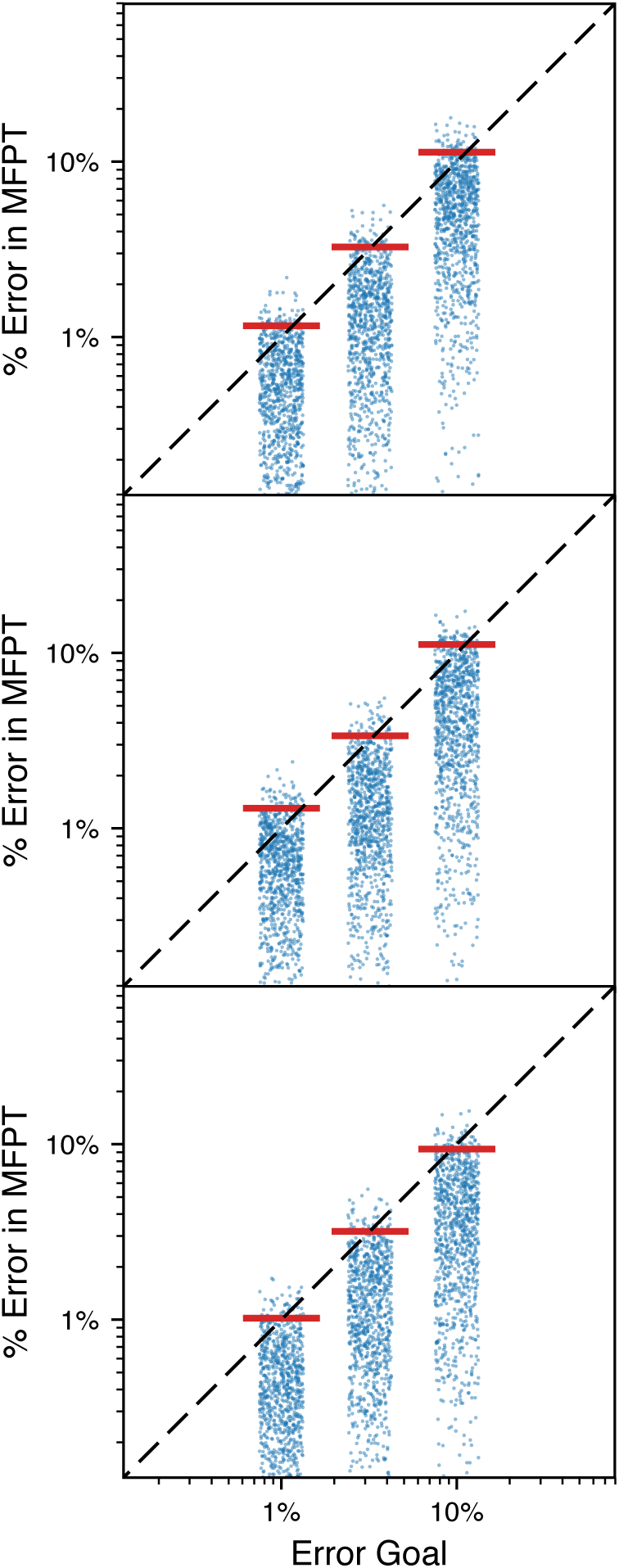
Actual error vs error goal of FFPilot simulations for (top) SRG_h=2.4_, (middle) SRG_h=2.3_, and (bottom) SRG_h=2.2_. Each strip shows (blue dots) 1000 independent FFPilot simulations and (red lines) the 95th percentiles of the errors.

We also looked at the contribution of phase 0 to the overall cost of the pilot stage. Applying Eq 21 to values from Table III, we found that for all variants of SRG phase 0 required around *∼*35% of the total simulation time. This finding is in contrast to the long-standing assumption in the FFS literature that phase 0 does not significantly contribute to the cost of a simulation and should therefore be extensively sampled^48^.

##### E. Genetic Toggle Switch Model

The last model we investigated using FFPilot was a more complex gene regulatory network, one of a family of extensively studied systems commonly referred to as a genetic toggle switch (GTS)^60–65^. Our GTS has seven species that interact with one another via fourteen reactions, all of which are first or second order (see Table IV). GTS consists of a single piece of operator DNA, *O*. When *O* is not bound to anything it can produce either of two proteins, *A* and *B*. *A* and *B* can both decay, they can both form homodimers, and those dimers can both bind back to *O*. Only one dimer can bind to *O* at any given time. When *O* is bound to a dimer of either protein, it can only produce more of that same protein (see Fig 1).

**TABLE IV.** Reaction scheme for our Genetic Toggle Switch (GTS) models.

| <u>Reactions</u> |  |  |  | <u>Rate Constants</u> |  | <u>Description</u> |
| --- | --- | --- | --- | --- | --- | --- |
| *Protein A* |  | *Protein B* |  | *forward* | *reverse* |  |
| $O$ | $\longrightarrow O + A$ | $O$ | $\longrightarrow O + B$ | $\theta$ | | expression |
| $A$ | $\longrightarrow \emptyset$ | $B$ | $\longrightarrow \emptyset$ | $\theta/4$ | | decay |
| $2A$ | $\rightleftharpoons A_2$ | $2B$ | $\rightleftharpoons B_2$ | 5 | 5 | dimerization |
| $O + A_2$ | $\rightleftharpoons OA_2$ | $O + B_2$ | $\rightleftharpoons OB_2$ | 5 | 1 | operator binding |
| $OA_2$ | $\longrightarrow OA_2 + A$ | $OB_2$ | $\longrightarrow OB_2 + B$ | $\theta$ | | bound expression |

**TABLE V.** Parameterizations and other simulation data from our GTS models. Per-phase weights, costs, and sample counts were estimated from the average results of 1000 FFPilot simulations (1% error goal). *MFPT* values were estimated from the results of 10^5^ DS simulations, plus or minus the 95% confidence interval bounds.

| parameter | GTS$_{\theta=.1}$ | | | | | GTS$_{\theta=1}$ | | | | | GTS$_{\theta=10}$ | | | | |
| --- | --- | --- | --- | --- | --- | --- | --- | --- | --- | --- | --- | --- | --- | --- | --- |
| $\theta$ | .1 | | | | | 1 | | | | | 10 | | | | |
| $\mathcal{A}$ | $O = 0, OA_2 = 1, OB_2 = 0, A = 4, B = 0, A_2 = 16, B_2 = 0$ | | | | | | | | | | | | | | |
| $\mathcal{B}$ | $O = 0, OA_2 = 0, OB_2 = 1, A = 0, B = 4, A_2 = 0, B_2 = 16$ | | | | | | | | | | | | | | |
| phase $^a \lambda_i$ | $^b w_i$ | $^c c_i^f$ | $^d c_i^s$ | $^e c_i$ | $^f n_i \times 10^{-3}$ | $w_i$ | $c_i^f$ | $c_i^s$ | $c_i$ | $n_i \times 10^{-3}$ | $w_i$ | $c_i^f$ | $c_i^s$ | $c_i$ | $n_i \times 10^{-3}$ |
| 0 -27.0 | 453 |  |  | 453 | 1664 | 45.3 |  |  | 45.3 | 1337 | 4.52 |  |  | 4.52 | 985 |
| 1 -22.5 | .092 | 19.7 | 63.5 | 23.7 | 8010 | .092 | 1.97 | 6.35 | 2.37 | 6436 | .090 | .197 | .643 | .238 | 4698 |
| 2 -18.0 | .271 | 148 | 113 | 138 | 1728 | .274 | 14.8 | 11.2 | 13.8 | 1382 | .281 | 1.51 | 1.12 | 1.40 | 977 |
| 3 -13.5 | .128 | 307 | 176 | 290 | 1897 | .136 | 30.7 | 17.1 | 28.8 | 1482 | .170 | 3.13 | 1.46 | 2.85 | 946 |
| 4 -9.0 | .111 | 442 | 120 | 406 | 1738 | .150 | 44.0 | 10.5 | 38.9 | 1201 | .339 | 4.46 | .597 | 3.15 | 568 |
| 5 -4.5 | .082 | 536 | 131 | 503 | 1849 | .242 | 52.6 | 10.1 | 42.3 | 859 | .676 | 4.98 | .529 | 1.97 | 356 |
| 6 0.0 | .437 | 659 | 118 | 423 | 684 | .636 | 61.4 | 8.93 | 28.0 | 451 | .872 | 5.14 | .463 | 1.06 | 268 |
| 7 4.5 | .632 | 830 | 167 | 411 | 467 | .711 | 73.6 | 12.7 | 30.3 | 365 | .873 | 5.83 | .657 | 1.32 | 240 |
| 8 9.0 | .861 | 994 | 140 | 259 | 310 | .858 | 85.1 | 11.1 | 21.6 | 276 | .918 | 6.43 | .599 | 1.08 | 208 |
| 9 13.5 | .940 | 1153 | 165 | 225 | 209 | .913 | 97.3 | 14.4 | 21.6 | 210 | .925 | 7.08 | .861 | 1.33 | 178 |
| 10 18.0 | .988 | 1283 | 122 | 135 | 116 | .973 | 108 | 11.5 | 14.1 | 140 | .960 | 7.58 | .809 | 1.08 | 141 |
| 11 22.5 | .997 | 1397 | 156 | 159 | 53 | .989 | 117 | 15.4 | 16.5 | 81 | .972 | 8.14 | 1.27 | 1.46 | 102 |
| 12 27.0 | 1.00 | 1493 | 150 | 150 | 19 | .998 | 126 | 14.9 | 15.1 | 37 | .989 | 8.61 | 1.36 | 1.44 | 63 |
| MFPT | $(7.015 \pm .043) \cdot 10^7$ | | | | | $(1.074 \pm .0067) \cdot 10^6$ | | | | | $(7.701 \pm .048) \cdot 10^3$ | | | | |

<sup>a</sup> The position of interface $i$ in terms of order parameter $\Delta$ .
<sup>b</sup> The phase weight of phase $i$ .
<sup>c</sup> The expected cost of a failed phase $i$ trajectory.
<sup>d</sup> The expected cost of a successful phase $i$ trajectory.
<sup>e</sup> The expected cost of a phase $i$ sample.
<sup>f</sup> The phase $i$ sample count (in thousands) predicted by FFPilot to achieve a 1% error goal, 95% confidence.

The combination of positive feedback (of monomer production on dimer/operator binding) and negative feedback (of dimer/operator binding on production of the competing monomer) gives GTS bistable dynamics^66^. The system as a whole switches between a state with high levels of the various forms of *A* and low levels of *B*, and a state with low levels of *A* and high levels of *B*.

For GTS we defined three order parameters. One, which we called Δ, is the difference between the total count of protein *B* and the total count of protein *A*. Another, which we called Σ, is the sum of the total count of protein *A* and the total count of protein *B*. The last, Ω, takes a value from [-1,1] based solely on the state of the single operator. In terms of the underlying species counts, the order parameters can be written as:

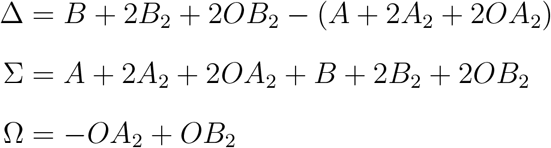

where *A* and *B* are the monomer counts, *A*_2_ and *B*_2_ are the dimer counts, and *OA*_2_ and *OB*_2_ are the dimer-operator complex counts. Equivalently, Ω can be said to have one of three categorical values:

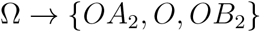

Although all of our GTS simulations are based on a stochastic master equation formulation of the system, it is helpful to consider the more straightforward deterministic formulation when trying to understand the system’s overall behavior (see supplemental Table S1 for the deterministic rate equations). For a given set of parameters, the fixed points of the deterministic formulation can be found. For all of the parameter sets we used in our simulations there are three fixed points in terms of Δ, two stable fixed points and one unstable fixed point. One of the stable fixed points corresponds to the state with a high level of *A* and a low level of *B*, and the other stable fixed point corresponds to the state with a low level of *A* and a high level of *B*. We refer to these two states as *A* and *B*, respectively.

We wanted to be able to tune the rarity of the 𝒜 → ℬ event without disrupting the overall dynamics of GTS. To do so, we added a relative protein turnover parameter *θ*. The rate constants of all of the expression and decay reactions for both *A* and *B* are multiplied by *θ*. Since *θ* does not affect the birth/death ratio of each protein, the steady state levels of both *A* and *B* are constant with respect to *θ*. However, *θ* does have a large effect on the rate of 𝒜 → ℬ switching. We used three different variants of GTS in our simulations, GTS*_θ_*_=.1_, GTS*_θ_*_=1_, and GTS*_θ_*_=10_, see Table V for complete parameters.

For all FFPilot simulations we used Δ as the order parameter. We tiled the state space in terms of Δ by placing *λ*_0_ at Δ = *−*27, *λ_N_* at Δ = 27, and then placing 11 more interfaces evenly spaced in between.

As with SRG, we were interested in the distribution of phase 0 inter-event times of GTS in order to establish the validity of Eq 30 with respect to GTS. Fig 6 shows the results of our phase 0 inter-event time distribution simulations. The phase 0 distributions of the different GTS variants differ a great deal, but interestingly their 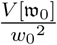 values (the ratio of moments required for Eq 30) are very similar. 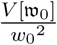 was found to be 8.15, 8.15, and 8.41 for GTS*_θ_*_=.1_, GTS*_θ_*_=1_, and GTS*_θ_*_=10_, respectively.

**FIG. 6.**
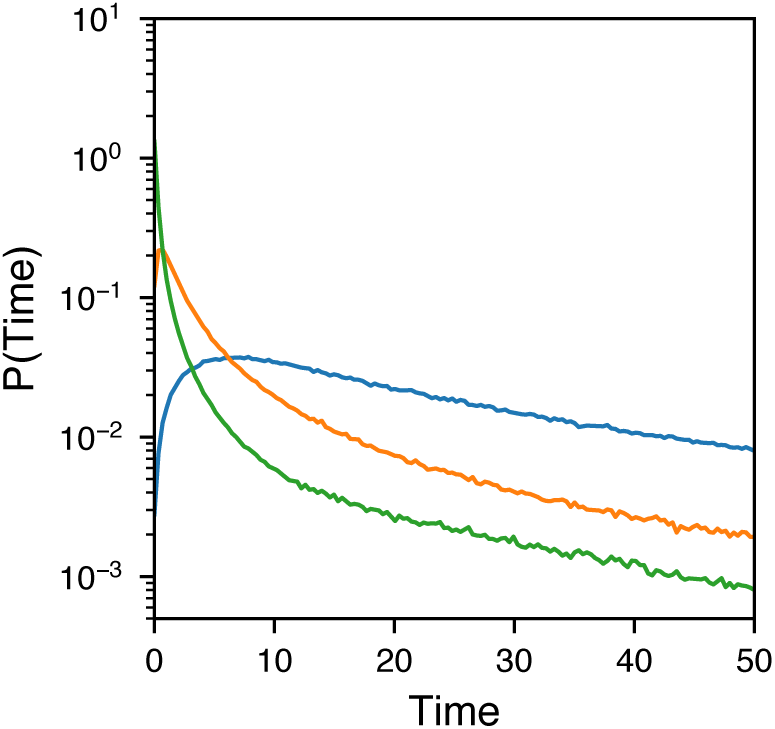
Waiting times between forward crossing events during phase 0 of FFS simulations of the GTS models. Each line shows a distribution calculated from 10^6^ samples from a single phase 0 trajectory of (blue) GTS*_θ_*_=.1_, (orange) GTS*_θ_*_=1_, and (green) GTS*_θ_*_=10_.

The phase weights from the FFPilot simulation can also be used to reconstruct the stationary probability density function (PDF) of the system. We used the output of a 10% error goal simulation from each of GTS*_θ_*_=.1_, GTS*_θ_*_=1_, and GTS*_θ_*_=10_ to generate the PDFs shown in Figs. 7 and S19. As can been seen from a comparison with the PDFs generated from extensive direct sampling, FFPilot reproduces a highly accurate PDF, especially in the low probability transition region, and with much lower computational cost than direct sampling. In the PDF of the GTS, we see the two stable fixed points we expect and a transition path between them with a barrier that increases in height as *θ* decreases. Unlike other GTS models where authors have observed three stable state^61, 62, 65^, our model has by design only two states in the region of parameter space in which we are working.

We next ran a test to examine how well the full FFPilot protocol was able to control sampling error in estimations of *MFPT* of the 𝓐 *→* 𝓑 switching process. We executed 1000 FFPilot simulations of GTS*_θ_*_=.1_, GTS*_θ_*_=1_, and GTS*_θ_*_=10_ using error goals 1%, 3.2%, and 10%. We estimated *MFPT* for each simulation, then found the percent errors relative to the results from high accuracy DS simulations of equivalent models, which were run with a 0.62% error goal.

The percent errors of the *MFPT* estimates are shown in Fig 8. As can be seen in the figure, the 95th error percentiles (marked by the red lines) are located somewhat above *x* = *y*, indicating that the overall errors in the estimated *MFPT* values are above the desired errors. The 95th percentile lines do decrease along with error goal, implying that FFPilot partially but not completely controls error in simulations of GTS. The anomalous dispersion decreases as the height of the barrier between 𝒜 and ℬ in probability space increases. This implies that the extra error is caused by a system dependent property and is not directly related to undersampling.

**FIG. 7.**
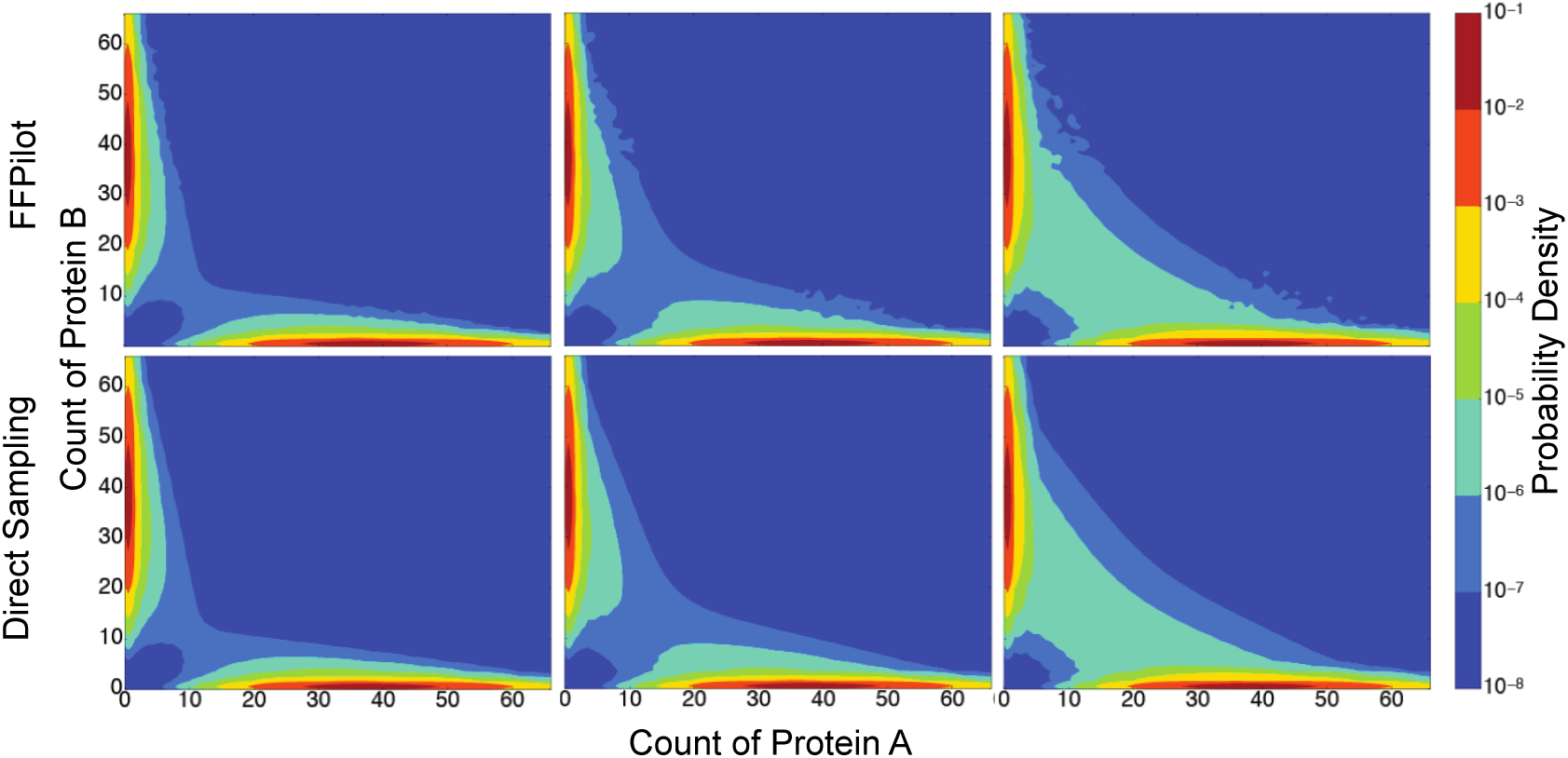
Stationary probability distributions for (left) GTS*_θ_*_=.1_, (center) GTS*_θ_*_=1_, and (right) GTS*_θ_*_=10_. The top row was calculated using FFPilot and the bottom row using direct sampling.

**FIG. 8.**
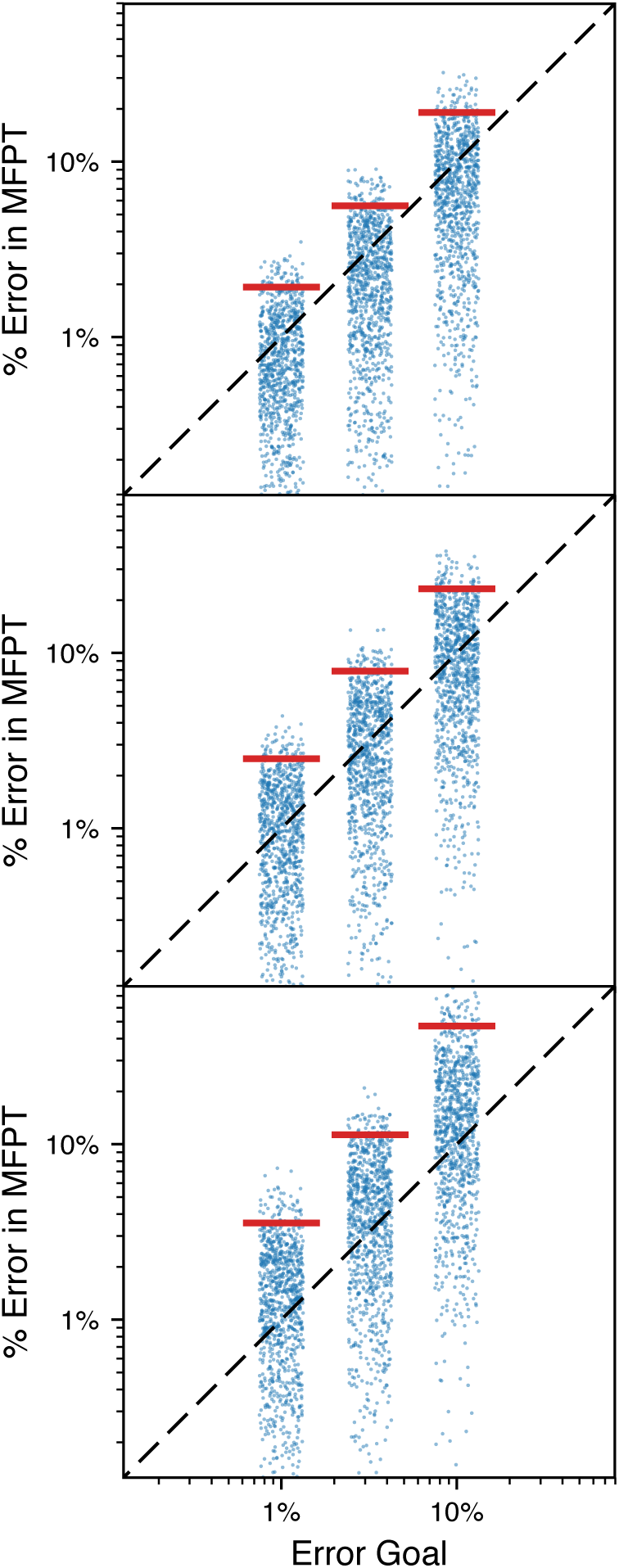
Actual error vs error goal of FFPilot simulations for (top) GTS*_θ_*_=.1_, (middle) GTS*_θ_*_=1_, and (bottom) GTS*_θ_*_=10_. Each strip shows (blue dots) 1000 independent FFPilot simulations and (red lines) the 95th percentiles of the errors.

We again found that phase 0 contributed significantly to the cost of GTS simulations. From parameters listed in Table V and Eq 21, we calculated the share of total simulation time consumed by phase 0, which was found to be 19%, 23%, and 33% for GTS*_θ_*_=.1_, GTS*_θ_*_=1_, and GTS*_θ_*_=10_, respectively.

##### F. Interface Landscape Error in Genetic Toggle Switch

We sought to understand the causes of the extra error in the GTS simulations. We chose the condition with the largest anomalous errors, GTS*_θ_*_=10_ executed with an error goal of 10%, and examined the phase weight estimates produced by each of the 1000 replicate simulations we had run with that condition. These phase weights are shown in the top half of Fig 9. The phase weights estimated by an equivalent FFPilot simulation run with an error goal of 0.1% are shown as dashed lines, and serve as a point of reference (there is no exact method for extracting the phase weights from a DS simulation). The dispersion of phase weight estimates around the reference weight is much greater in certain phases, especially phases 4 and 5. By itself, this is not an indication that FFPilot is failing to correctly estimate sampling counts for these phases. By design, FFPilot allows for different levels of dispersion in different phases when it is determining the optimal simulation plan.

**FIG. 9.**
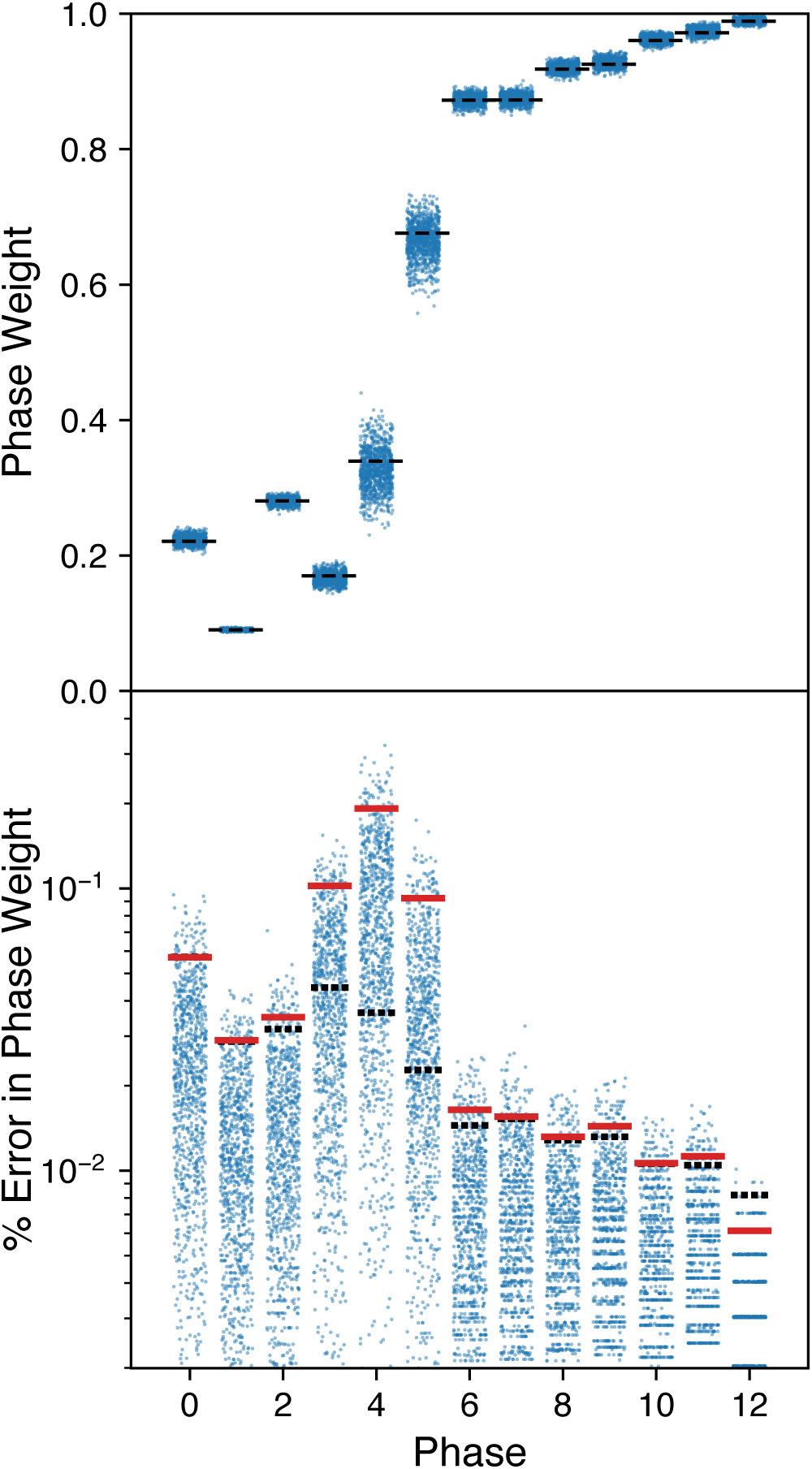
GTS*_θ_*_=10_ phase weight estimates and errors. (top) The phase weights as estimated by (blue dots) 1000 independent FFPilot simulations run to a 10% error goal. The dashed black lines show the phase weights from an FFPilot simulation with a 0.1% error goal. (bottom) The blue dots show the percent error of each phase weight estimate given in the top subplot, relative to the 0.1% error goal simulation. Also shown are (red lines) the observed 95th error percentiles, and (dashed black lines) the expected 95th error percentiles.

In order to determine how much of the phase weight dispersion represents FFPilot functioning as intended and how much of the dispersion is truly anomalous, we calculated an optimal set of error goal targets for each simulation phase. From the FFPilot optimizing equation (Eq 30) we derived analytic expressions for the per-phase error goals:

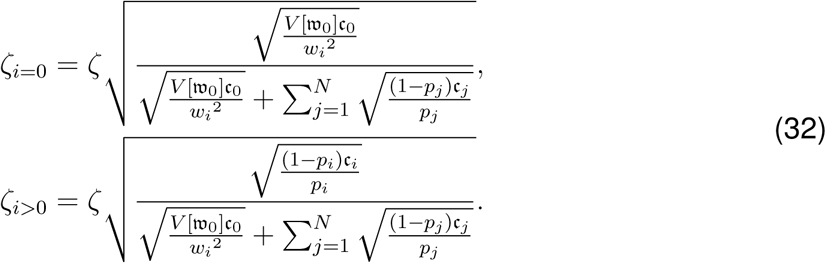

Just as with the overall error goal, in any given phase 100 *· α*% percent of simulations will have a level of sampling error in the phase weight estimate at or below the relevant per-phase error goal *ζ_i_*. We parameterized Eq 32 using the phase weight and phase cost estimates from the very high accuracy (0.1% error goal) FFPilot simulation of GTS*_θ_*_=10_ mentioned above.

The error goal targets we calculated are shown as the dashed lines in the bottom half of Fig 9. The phase weight percent errors (calculated relative to the estimates from the 0.1% error goal simulation) are shown as dots, and the red lines mark the 95th percentiles of the errors. If the red line and the dashed line overlap for a particular phase, it means that the error in this phase is dominated by sampling error, which FFPilot is able to completely account for. If the red line is above the dashed line, then there is more error occurring in that phase than was predicted by FFPilot, and the magnitude of the separation of the two lines represents the magnitude of the anomalous (as opposed to the predicted) error. Interestingly, FFPilot estimates the majority of phase weights to within the desired error goal. The extra error in the *MFPT* estimate appears to be due primarily to extra error in only three of the phase weight estimates, those from phases 3-5. Further, the bulk of the extra error is concentrated in just two of those phase weight estimates, those from phases 4 and 5. Interfaces *λ*_4_ and *λ*_5_ also happen to be the interfaces immediately preceding the transition midpoint.

We hypothesized that there must be some particular feature of the state space landscape of GTS that the simulations are exploring during phases 4 and 5 that is responsible for the anomalous error. We further reasoned that the same features that are responsible for what Allen and coworkers call landscape variance^48^ could be related to the increased error. Here we define a new source of error, which we call landscape error, that is due to two factors: (1) misrepresentation of some regions of the state space in the set of trajectory starting points at *λ_i_*, and (2) significant differences in *P* (*λ_i_|λ_i−_*_1_) as a function of trajectory starting state. The total probability factor *P* (*λ_i_|λ_i−_*_1_) that is measured in each phase *i >* 0 can be thought of as a mixture of many independent probabilities, one for each state along *λ_i−_*_1_, weighted by the (normalized) count of times the state is used as a starting point for a phase *i* trajectory. If either of factors (1) or (2) occurs alone, *P* (*λ_i_|λ_i−_*_1_) will still be correctly estimated. However, if the factors occur together they can lead to significant errors. In other words, if the landscapes assembled at *λ*_3_ and *λ*_4_ are heterogeneous across replicate simulations, and if differences in those landscapes can lead to differences in the effective value of *P* (*λ_i_|λ_i−_*_1_), then landscape error may be the cause of the anomalous simulation error we observe in our simulations of GTS. Of particular importance is the fact that the landscape error in phase *i* is due to errors in the landscape assembled from the endpoints of successful trajectories launched during phase *i −* 1. Thus, no amount of extra sampling performed during phase *i* alone can completely abolish landscape error.

We wanted to test if the conditions for landscape error were present in FFPilot simulations of GTS*_θ_*_=10_. To this end, we chose two simulations, which we will call replicate 6 and replicate 8, that had a large divergence in their *MFPT* estimates. Looking at the phase weights estimates produced by these two simulations (Fig 10), the divergence in the *MFPT* estimates can be seen to be mostly due to divergence in the phase weight 4 and 5 estimates (as expected). Exploring phase 5 in greater depth, we calculated the landscape occupancy along *λ*_4_ in terms of the orthogonal order parameter Σ. The *λ*_4_ occupancies for replicates 6 and 8 *P* (Σ|Ω*, λ*_4_) (which are binned by Σ and operator state Ω), are shown in the lower left-hand subplots of Fig 11 as lines colored blue or gold, respectively. Replicate 6 has somewhat higher occupancy in the *OA*_2_ and *O* states, and replicate 8 has higher occupancy in the *OB*_2_ states. Thus, condition (1) for landscape error in phase 5, heterogeneous occupancies along *λ*_4_, is indeed satisfied.

**FIG. 10.**
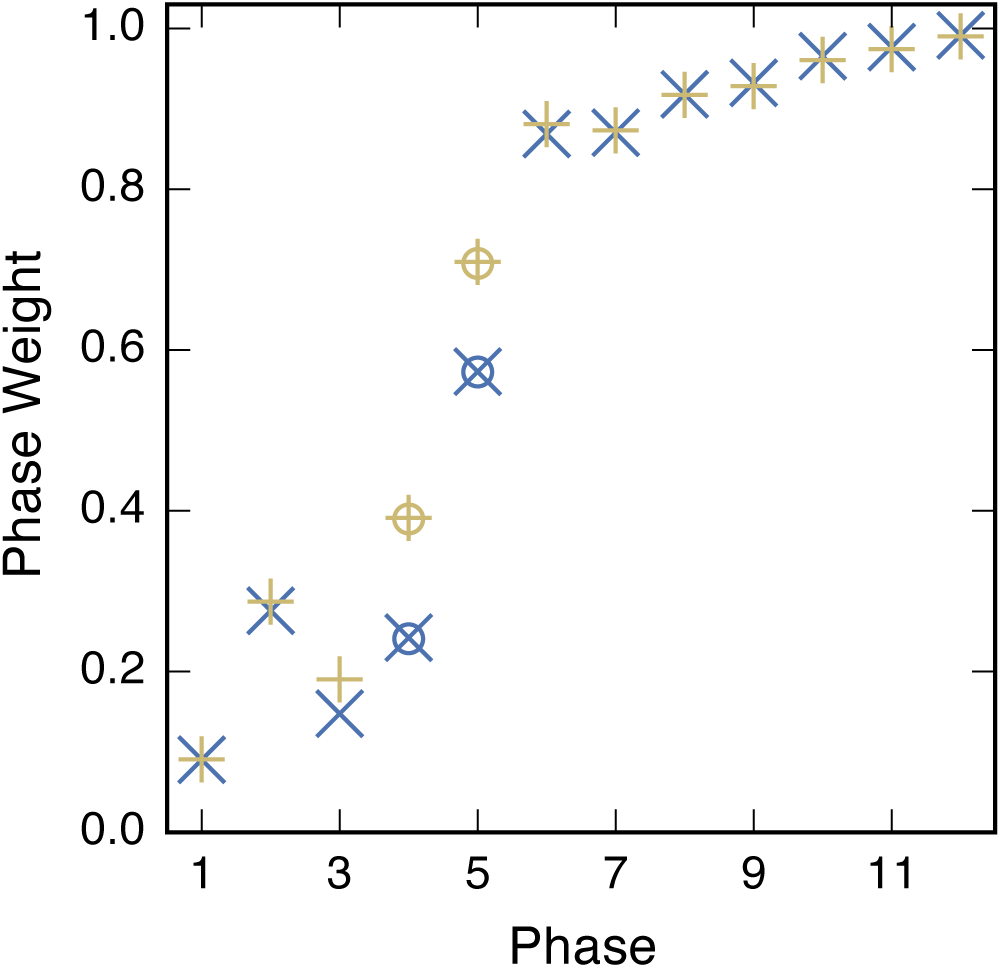
The phase weights of two replicates: (blue x) Replicate 6 and (gold +) Replicate 8, from a GTS*_θ_*_=10_ FFPilot simulation run to an error goal of 10%. Circles show weights for phases 4 and 5 calculated via an alternative method using Eq 33.

**FIG. 11.**
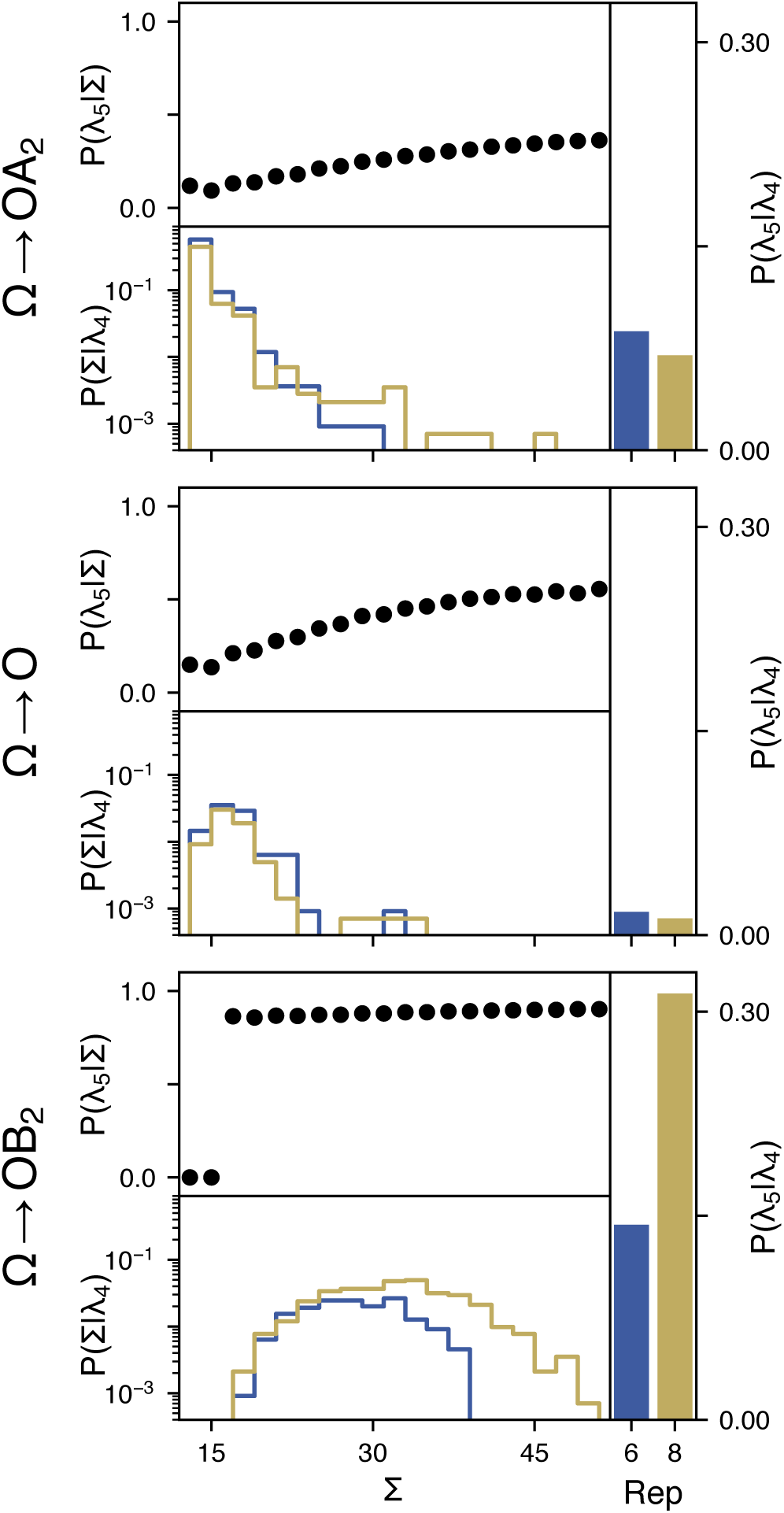
Breakout of the calculation of phase weight 5 for replicates 6 and 8. Each group of three plots show data from a slice of GTS state space with a different Ω, the state of the operator DNA. Each subplot within a group shows different probability data. (dots) The probability of a trajectory launched from a point on *λ*_4_ fluxing forward to *λ*_5_ vs the orthogonal order parameter Σ. (lines) The occupancy along Σ at interface *λ*_4_ for (blue) replicate 6 and (gold) replicate 8. (bars) The cumulative probability of fluxing forward across *λ*_5_ when starting at the specified Ω for (blue) replicate 6 and (gold) replicate 8.

Next, we launched 10^6^ independent trajectories from each starting state along *λ*_4_ that had non-zero occupancy in either replicate 6 or 8. Just as in a normal FFPilot simulation, we stopped each trajectory when it either reached the next interface or fell back into the initial basin, and we took note of the stopping states. This gave us a highly accurate estimate of *P* (*λ*_5_*|λ*_4_) as a function of trajectory starting state. These state-dependent probability values *P* (*λ*_5_|Σ, Ω*, λ*_4_), (which are also binned by Σ and operator state Ω), are shown as filled circles in Fig 11. *P* (*λ*_5_|Σ, Ω*, λ*_4_) varies a great deal across both Σ and Ω, meaning that condition (2) is also satisfied for our simulations of GTS*_θ_*_=10_, and that landscape error is indeed a possible explanation for the anomalous error.

In order to test if landscape error alone is a sufficient explanation for the observed anomalous error, we recalculated the phase 4 and 5 weights of replicates 6 and 8 from their landscapes alone, according to:

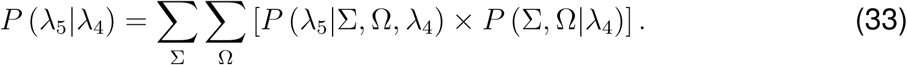

If landscape error is indeed the sole cause of the anomalous error, we expected that the phase weights derived from Eq 33 would closely match those originally estimated by replicates 6 and 8, even though these original estimates vary greatly between the replicates. In this view, the phase *i >* 0 weight estimate produced by each simulation converges (with increasing trajectory count) to a unique value of *P* (*λ_i_|λ_i−_*_1_), as determined by their heterogeneous samples of the landscape along *λ_i−_*_1_. The recalculated phase weight values are plotted as empty circles in Fig 10, and they do indeed closely agree with the original estimates. Thus, we conclude that sampling error is indeed being handled correctly by FFPilot, and the anomalous error in our GTS simulation results is due to landscape error.

##### G. Eliminating Landscape Error in GTS via Oversampling

Although a complete mathematical treatment of landscape error is beyond the scope of this paper, we wanted to investigate strategies for eliminating landscape error within the limited context of GTS. To this end we ran FFPilot simulations of GTS*_θ_*_=10_ under a variety of different oversampling schemes. As can be seen in Fig 9, for GTS*_θ_*_=10_ landscape error is mainly an issue in phases 3-5. Based on this, we initially we hypothesized that increasing sampling by 10X in phases 2-4 (*i.e.* increasing sampling in each of the phases preceding the problematic phases) would abolish landscape error.

The results from 1000 simulations of GTS*_θ_*_=10_ with 10X phase 2-4 sampling at an error goal of 10% are shown in the second column of Fig 12. Oversampling in these phases alone has only a minor effect on simulation error. Under these conditions the 95th percentile of error was 38%, whereas the error without oversampling is 47%. Based on these results, we tried out a more expansive oversampling strategy. In addition to over-sampling by 10X in phases 2-4, we oversampled by 20X in phase 0 and 10X in phase 1 as well. The results from 1000 simulations run with 20X phase 0, 10X phase 1-4 oversampling are shown in the last column of Fig 12. Error is reduced dramatically under these conditions, to just under 10%. We also tried 10X phase 0 oversampling, but found that it was not quite sufficient to eliminate landscape error (see supplemental Fig S5).

**FIG. 12.**
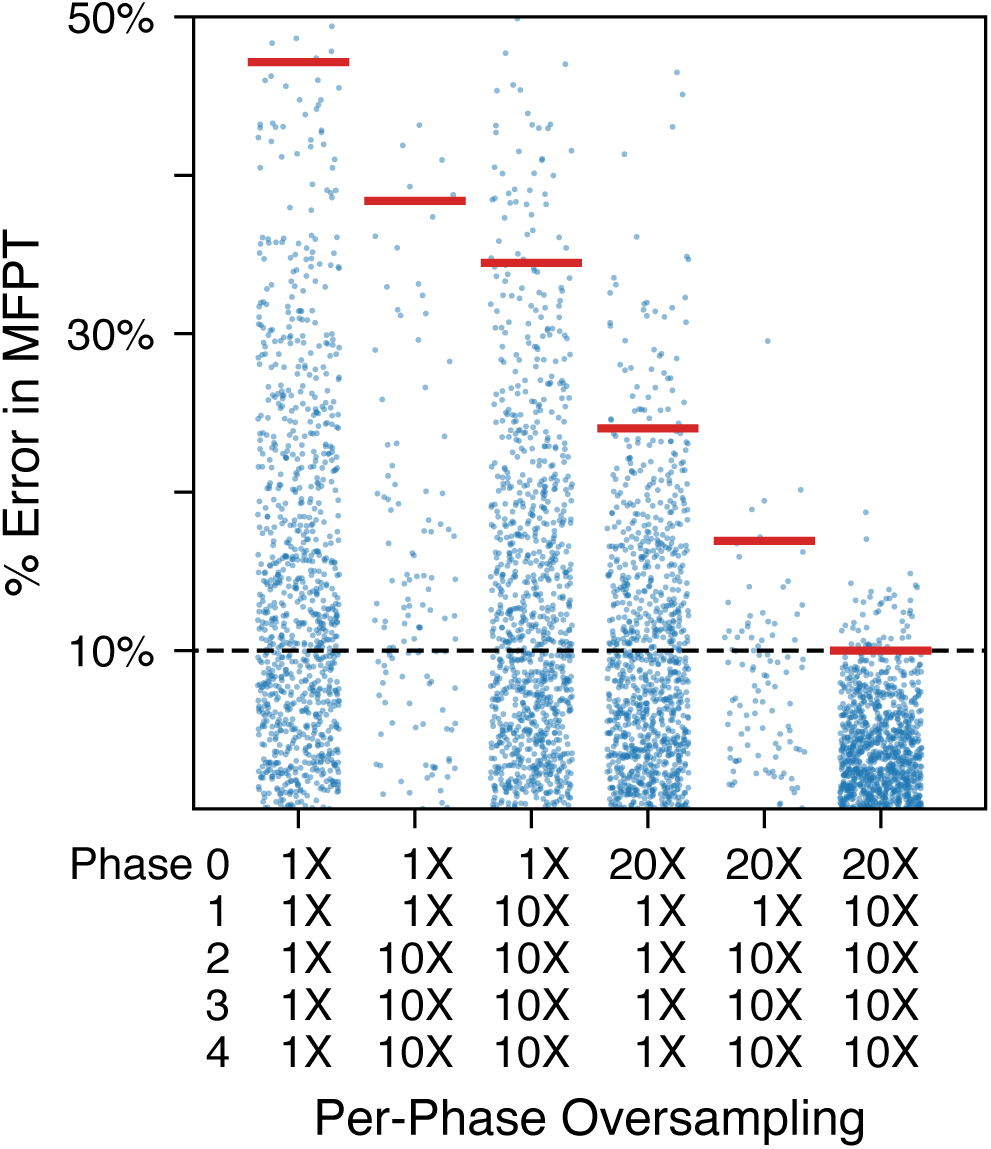
Errors from simulations executed with various oversampling schemes. The extent of oversampling in each phase for the separate schemes is indicated beneath each column of results. All simulations were of GTS*_θ_*_=10_ run to an error goal of 10%. Red lines show the 95th error percentiles.

Thus, oversampling in phases 2-4 alone had almost no effect, but oversampling in phases 0-4 was enough to eliminate the landscape error. In order to understand this difference, we eliminated oversampling in each phase individually. The results from these simulations are shown in supplemental Fig S6. The effect of skipping oversampling in phase 0 is particularly dramatic, leading to an increase in simulation error of nearly 25%. So long as phase 0 is being oversampled, the increase in simulation error from skipping oversampling in any of phases 1-4 is less dramatic (2%-7%) but still significant. This implies that landscape errors are correlated. Effectively, defects in the sampled landscape distribution at any *λ_i_* may carry over to *λ_i_*_+1_.

##### H. Relationship between Landscape Error and Phase Weight Covariance

To further investigate the origin of the landscape error, we considered whether covariance between the phase weight estimators could account for the extra error. Recall that in deriving Eq 18 we assumed that the phase weight estimators 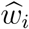 were independent and that the off-diagonal elements of the covariance matrix Σ were zero. When this assumptions does not hold there will be additional unaccounted for variance in the MFPT estimator 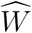.

To check this assumption for our GTS model, we numerically estimated the covariance matrix for the case where *θ* = 10. We performed 100 independent FFPilot and FFS simulations at a variety of accuracy levels. For FFPilot simulations we used error goals of 1%, 3.2%, and 10% and for FFS simulations a constant number of trajectories per interface from 1,000 – 25,000. Fig S9+S10 show the distribution of phase weight estimators obtained during the FFS runs. We next calculated the variance of each 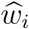 and for *i >* 0 compared to the expected variance obtained from Eq 14 and Eq 28:

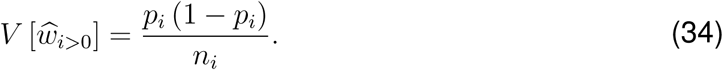

Fig 13a and S11+S12 show the comparison. For phases 3-5 the variance is much greater than expected, which agrees with our earlier analysis revealing which phases had the most uncontrolled error.

We then calculated the covariance matrix from the simulations. Fig 13b and S13 show the results. A clear cluster of high covariance is apparent between phase 3-5 and, additionally, phase 0 has high covariance with most of the other phases. However, phase 0 has higher absolute variance than the other phases and if one instead looks at the Pearson correlation coefficient (see Fig S14) that correlation between phase 0 disappears while the high correlation between phases 3-5 remains.

Given that the phases with high covariance correspond to the phases with high error and we were able to control the extra error using oversampling, we wanted to know how oversampling affected the covariance matrix. We applied the same oversampling strategy discussed in the previous section and recalculated the covariances. The bottom row in Fig 13 shows the covariance results with oversampling. When we applied oversampling in phases 0-4 the variances were near their expected levels and the magnitude of the covariances decreases substantially. It is interesting to note in Fig 13 how the variance for phase 5 moves to the expected value, even though no additional sampling was done in phase 5. As we discussed earlier, increasing the sampling in earlier phases can reduce the error in later phases through the landscape correlations.

**FIG. 13.**
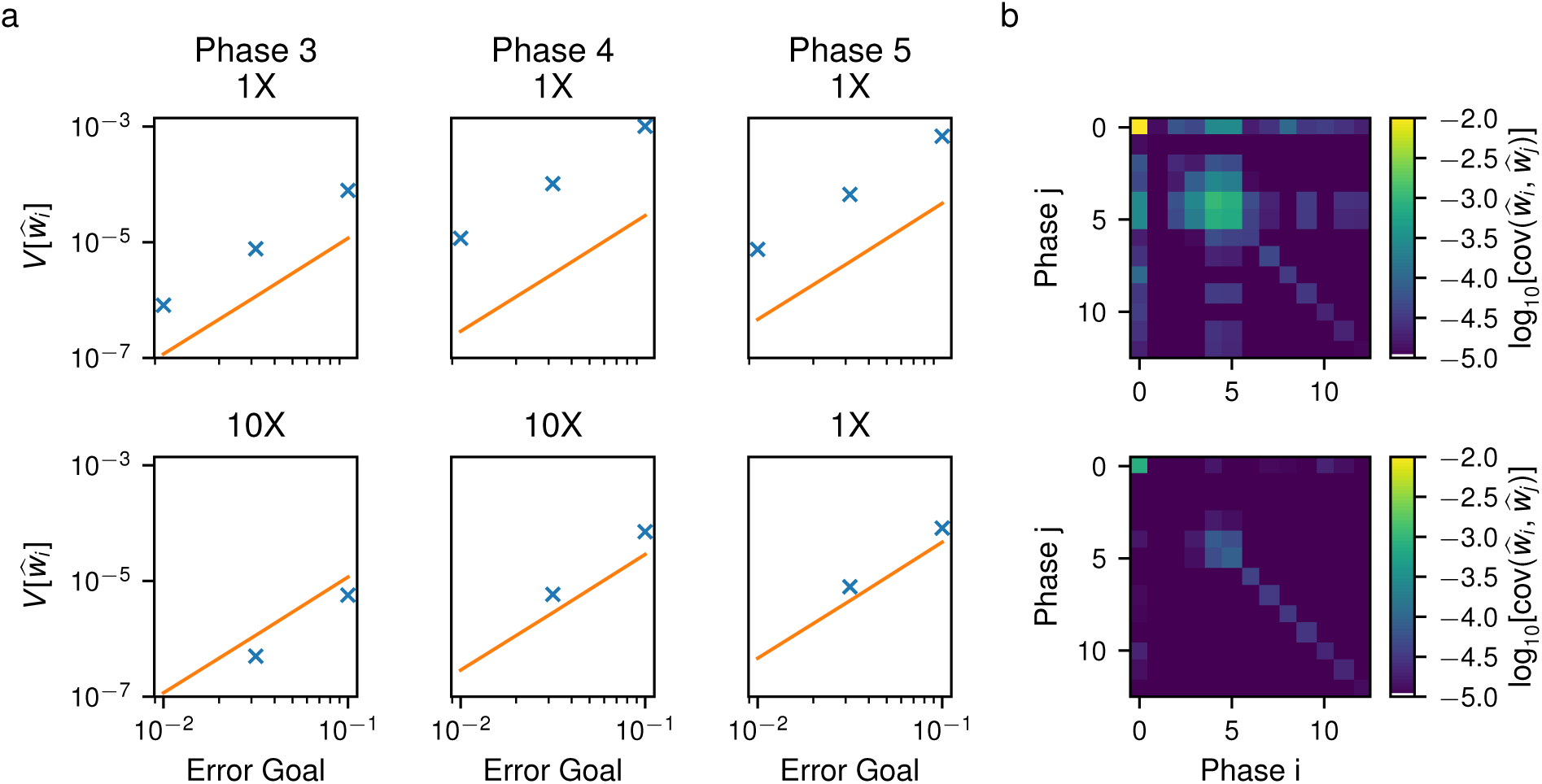
(a) Comparison of the phase weight estimator variances for the indicated phases. Shown are (x) the measured variance from 100 independent FFPilot simulations and (line) the expected variance from Eq 34. The top row shows the variances with no oversampling and the bottom row shows the variances with the optimal oversampling: 20X,10X,10X,10X,10X for phases 0–4, respectively. Note that phase 5 was not oversampled. (b) The covariance matrices for the phase weight estimators at 10% error goal (top) without and (bottom) with oversampling.

We also tried various combinations of oversampling. Fig S15 shows the results of two different experiments. First, we oversampled sequential phases one at a time, starting with phase 0. In this case the covariance gradually diminishes as more phases are oversampled. Second, we started with oversampling only phase 4 and then sequentially added prior phases, down to phase 0. In this case the covariance does not gradually change. It remains roughly the same until phase 0 is added to the oversampling and then it dramatically jumps lower. Again, this agrees with our previous conclusion that oversampling starting in phase 0 is necessary to reduce underrepresentation of trajectories likely to cross the barrier that might relax slowly during the crossing. It is this slow relaxation that drives the landscape correlation errors in our GTS model.

##### I. Theoretical Efficiency of DS, FFS, and FFPilot Simulations

Enhanced sampling is commonly assumed to be more efficient than DS. By controlling for simulation error, a direct comparison can be made between DS and FFPilot simulations, and the speedup of one simulation method versus the other can be assessed.

It is straightforward to derive an expression for the cost of a DS simulation 𝓒_ds_ as a function of the error goal. Plugging Eq 12 into Eq 11 yields:

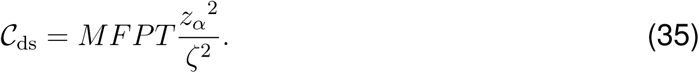

The cost of an FFS simulation 𝓒_ffs_ is given by Eq 21: 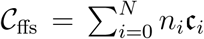. We can plug the FFPilot optimizing equation (Eq 30) into Eq 21 in order to expand the *n_i_* values. This yields an expression for 𝓒_ffs-opt_, the cost of an optimized FFS simulation given *a priori* knowledge of the parameters required for the optimizing equation:

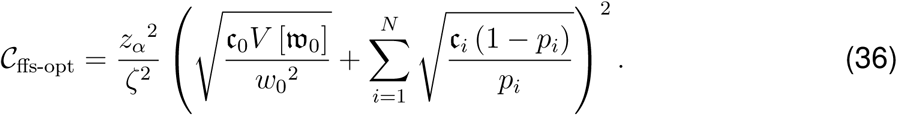

Next, we examined the theoretical efficiency of the FFPilot approach. Due to the FFPilot optimizing equation, the production stage of an FFPilot simulation can be thought of the most computationally efficient FFS simulation possible with respect to a given error goal. However, if the pilot stage is too expensive it may be possible that in general FFPilot is inefficient relative to the traditional FFS algorithm. Thus we wanted to determine if FFPilot simulation is reasonably efficient, and, if so, under what conditions.

The total cost of an FFPilot simulation 𝓒_ffpilot_ can be found by adding a second term to the RHS of Eq 36 that specifically accounts for the extra runs performed during the pilot stage:

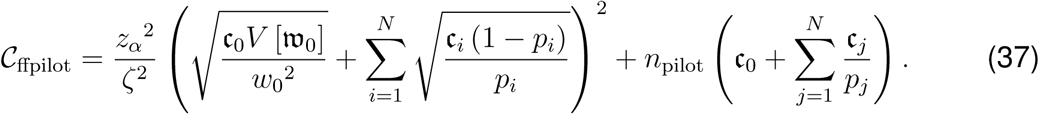

Eq 37 gives 𝓒_ffpilot_ as a function of error goal. The production stage cost increases with error goal 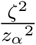 whereas the pilot stage cost remains fixed. For a low enough error goal, the pilot stage term in 𝓒_ffpilot_ will be negligible compared to the overall simulation cost. For simulations run with *n*_pilot_ = 10^4^ the approximation 𝓒_ffpilot_ *≈* 𝓒_ffs-opt_ holds when the error goal was *<*5% (see Fig 14).

**FIG. 14.**
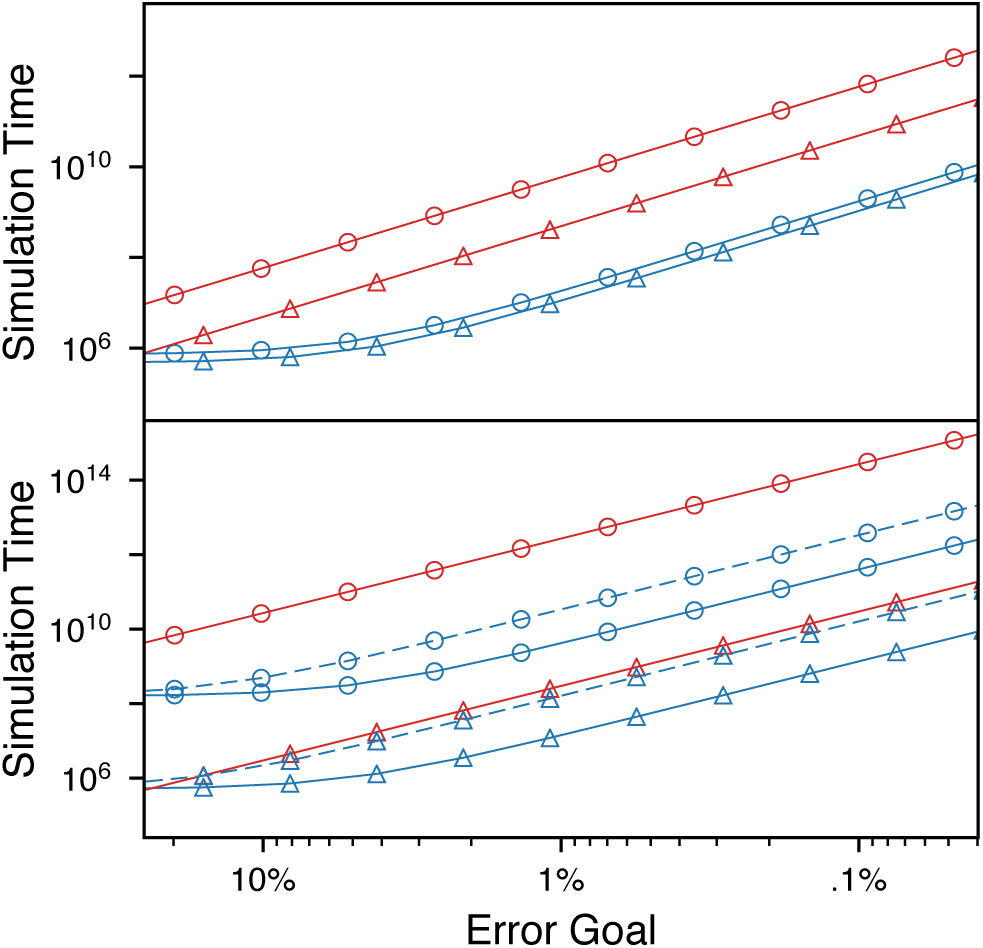
Simulation time vs error goal for several SRG and GTS models. Simulation time was calculated using Eq 35 for (red lines) DS, and Eq 37 for (blue lines) FFPilot and (dashed line) FFPilot with oversampling to correct the landscape error. Parameters were taken from Tables III and V. (top) Data from (circles) SRG_h=2.4_ and (triangles) SRG_h=2.2_. (bottom) Data from (circles) GTS*_θ_*_=.1_ and (triangles) GTS*_θ_*_=10_.

##### J. Speedup of FFPilot Compared to DS and FFS with Equivalent Error

To illustrate the practical performance of our method, we compared FFPilot against both DS and traditional FFS for a matched margin of error. It is straightforward to use Eq 11 to calculate the number of DS trajectories needed to make a comparison against FFPilot. For traditional forward flux, however, the comparison is more difficult. There is no general way (apart for our optimizing equation) to determine the distribution of trajectories to launch at each phase to achieve a specific error goal. We settled for a FFS scenario in which we executed an identical number of trajectories *N* at each phase and then calculated the margin of error versus *N* empirically. We ran a large number of FFS simulations at several relatively low values of *N* and then extrapolated this relationship to the desired error goal (see Fig S9).

Since GTS is the most interesting model we considered, we compared our oversampled FFPilot simulations against DS and FFS. We ran simulations at 1%, 3.2%, and 10% error for each of our GTS models using all three methods. We then calculated the total number of simulation steps that were required for each, which is a fair readout of simulation cost. Fig 15 shows the results of the comparison for 1% error and Fig S10 for all three error goals. FFPilot ranges from 3X–400X faster than DS, with the speedup increasing exponentially with the barrier height, as expected.

**FIG. 15.**
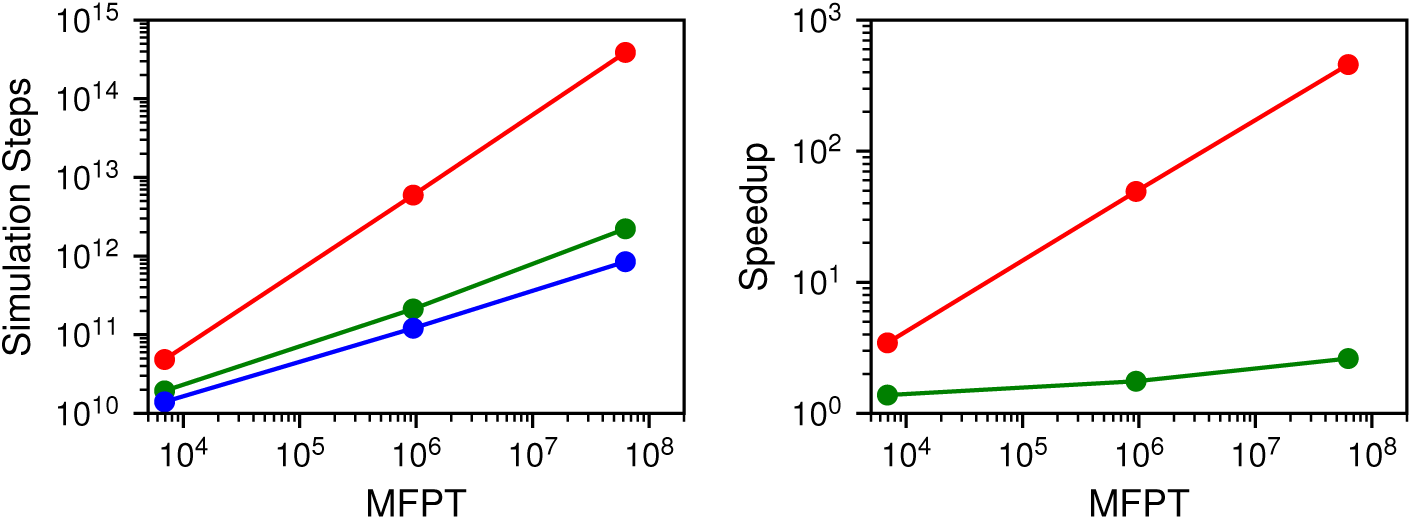
(left) Simulation steps vs *MFPT* for the three GTS models. The mean number of steps are shown for (red) DS, (green) traditional FFS with matched margin of error, and (blue) FFPilot corrected for landscape error. (right) The speedup of FFPilot relative to (red) DS and (green) traditional FFS. Connecting lines added for illustration.

Compared to traditional FFS, FFPilot has an average performance improvement of *∼*2X. This speedup means that by increasing the number of cheap trajectories and decreasing the number of expensive trajectories to be launched, FFPilot can offer a significant improvement in runtime while maintaining the same error. However, this analysis overlooks the fact that without running a sample set of simulations to calculate the optimal *N*, it is impossible for a user to know how many trajectories to specify for a traditional FFS simulation. Thus, the real utility of FFPilot over traditional FFS is not the runtime improvement, but the fact that it automatically calculates the best way for a FFS simulation to be configured to achieve a desired level of statistical error.

## IV. DISCUSSION

Above, we presented our FFPilot approach for automatically parameterizing a FFS simulation to both minimize simulation time and constrain sampling error at a user specified margin of error and confidence interval. By performing an inexpensive pilot simulation we obtain estimates of the weights and computational costs at each FFS interface, which are used to optimize a more thorough production simulation. Our pilot simulation approach also provides the advantage that individual interface placement, weights, and costs can be quickly evaluated before a potentially costly simulation is performed, if desired.

Unlike previous FFS optimization techniques, our method accounts for the statistics of phase 0 and also for varying computational cost along the order parameter. Both of these features are important for ensuring that error is controlled while minimizing simulation time. Particularly, optimizing the time spent in phase 0 is important as our results indicate that these calculations can consume a significant fraction of the total simulation time.

Our results show that FFPilot correctly controls the sampling error in FFS simulations. For one-dimensional systems, the *MFPT* estimates fall precisely within the specified error bounds. For higher dimensional systems, our testing reveals that while sampling error is well controlled by FFPilot, error due to the system dependent landscape pushes the total error outside of the specified bounds.

In the genetic toggle switch (GTS), substantial oversampling (from 10X–20X) in some phases is required to achieve the desired margin of error. As revealed by a detailed analysis, the anomalous error is due to underrepresentation of some parts of phase space in the crossing sets of early interfaces, especially phase 0. In the case of GTS, the system has a much higher probability of switching 𝓐 *→* 𝓑 from the Ω = OB_2_ state, but this state is rarely occupied during early phases and is thus subject to greater statistical variation. Additionally, the system relaxes much more slowly in the Ω dimension than in the other degrees of freedom, causing the crossing probability distributions at successive interfaces to be correlated in Ω. Errors in the estimation of early crossing distributions therefore lockin and cannot relax during later phases. These combined effects cause greater variability in the estimation of the phase weights than predicted and an increase in the total error.

Since all multi-dimensional systems of significant complexity likely relax more quickly in some degrees of freedom than others, a general approach is needed to control for landscape error in FFS while minimizing simulation time. Our analysis of the covariance matrix suggest that it may be possible to control for the landscape error in a revised formulation of our optimizing equation Eq 30 that includes the covariance matrix. However, it is not straightforward to estimate the covariances from the pilot simulation due to the lack of direct correspondence between individual trajectories in the various phases. Using the approach we followed here, calculating the covariances from many independent simulations, would be computationally prohibitive. At present we recommend first calculating the covariance matrix with a set of inexpensive low-accuracy FFPilot simulations and then using it to determine the optimal oversampling strategy for a more accurate production simulation.

Studying stochastic biochemical systems with metastable states will become increasingly important as more regulatory and developmental networks are elucidated in sufficient detail to permit quantitative modeling. Our findings raise the important issue of what other obstacles exist for efficient rare event sampling of these systems using FFS. For example, how to control landscape error in systems with many metastable states, with multiple transition paths, or with metastable intermediates? One possibility would be to study each transition path separately, using a branching approach, and then recombine the results based on the branching probabilities.

In summary, our FFPilot method provides a significant speedup compared to direct simulation of systems with rare event dynamics. Even with oversampling to control landscape error, speedups on the order of 100X for systems with long first passage times can be expected. The automatic optimization of simulation parameters to achieve a desired level of sampling error through the use of our optimization equation makes FFS simulations more robust and efficient.

## SUPPLEMENTARY MATERIAL

See supplementary material for additional derivations and figures.

## SOFTWARE AVAILABILITY

Code implementing the FFPilot algorithm is available as part of our LMES software package for high-performance simulation of stochastic biological models (https://www.robertslabjhu.info/home/software/lmes/). We also have created a tutorial describing in detail how to use LMES to execute FFPilot simulations (https://www.robertslabjhu. info/home/tutorials/tutorials/#ffpilot).

## ACKNOWLEDGMENTS

The authors thank the members of Roberts lab for discussions. This work was supported by the National Science Foundation under grant number PHY-1707961, and by the National Institutes of Health under grant T32 GM008403.

## Appendix A: The Upper Bound of the Variance of a Sum of Bernoulli Distributions

The overall process of launching trajectories in order to determine the phase weight during each phase of FFS simulation can be thought of as a single draw from a sum of many independent Bernoulli distributions, also called a Poisson binomial distribution (PBD). This viewpoint emphasizes the statistical equivalence between the outcome of the *j*th trajectory in an FFS phase, which will either fall back into its starting basin or flux forward to the next interface, and the outcome of the *j*th Bernoulli random variable in a PBD, which will take on a value of either 0 or 1. In both cases “success” (fluxing forward, drawing a 1) occurs with some probability *p_j_* inherent to the individual process, while “failure” (returning to basin, drawing a 0) occurs with probability (1 *− p_j_*). The expected value and the variance of a draw from a PBD of *n* terms (*i.e.* the sum over an independent draw from each of its constituent Bernoulli random variables) is:

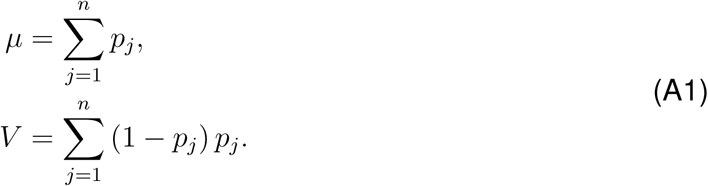

Equivalently, Eq A1 is also the mean and variance of the total count of successful trajectories *n^s^* in FFS phase *i >* 0, given that *n* = *n_i_* trajectories were run.

The variance of a PBD is maximized when the probability parameter of each of its Bernoulli random variables are all the same *p*:

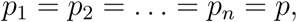

and so the upper bound on the variance of a sum of Bernoulli random variables is:

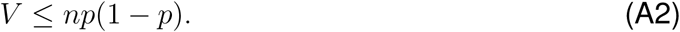

*Proof.* For a given value of *µ*, the method of Lagrange Multipliers can be used to maximize *V* with respect to the choice of particular values of the *p_i_* terms. Appropriate constraint and target equations can be taken from Eq A1:

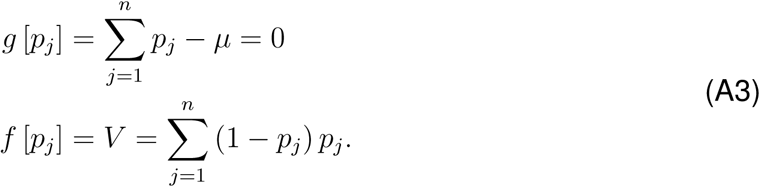

A Lagrangian can be formed from Eq A3:

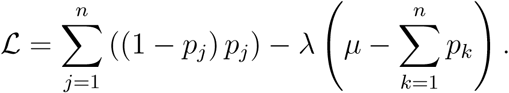

Next we find the gradient of the Lagrangian:

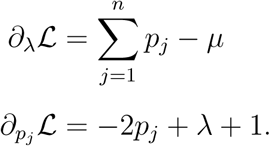

Now we set each part of the gradient to zero and solve the resulting set of equations. We start by solving for *p_i_* in terms of *λ*:

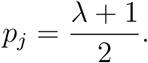

Next we solve for *λ* alone:

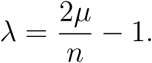

Then finally we plug the solution for *λ* into the gradient of *p_i_* and solve for *p_i_* alone:

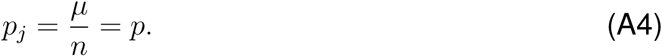

Thus, the spot at which every *p_i_* is equal to 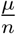 is a critical point for the variance of a Bernoulli mixture distribution. It can further be shown that the above critical point is a maximum for the constrained variance using the Bordered Hessian variation of the classical second derivative test. The Bordered Hessian^67^ of a Lagrangian can be defined as:

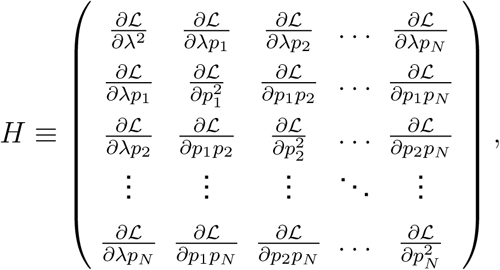

which for our Lagrangian works out to:

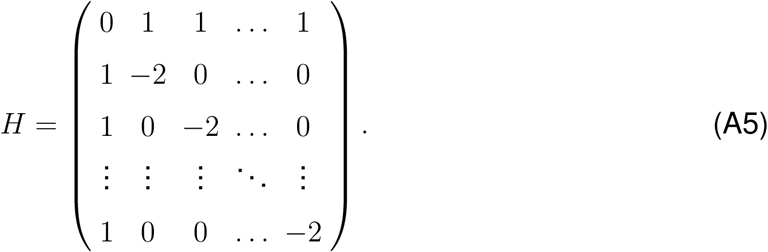

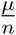 is a maximum if and only if *H* is negative definite. The negative definiteness of *H* can be demonstrated by showing that the signs of its leading principal minors demonstrate the appropriate alternating pattern^67^. For any finite value of *n*, this Hessian can be diagonalized using the Gaussian Elimination technique. This makes it easy to calculate the determinants of the various *x× x* upper-left submatrices and to show that the signs of said determinants do indeed follow the pattern of (*−*1)*^x−^*^1^ for *x ≥* 3, satisfying the condition for negative definiteness. This conclusion can be generalized to arbitrary values of *n* using a proof by induction (the details of which are omitted for brevity). Thus, *H* is always a negative definite matrix, and so setting each 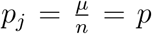 does indeed maximize *V* for a given *µ*.

From Eq A2, the upper bound on the variance of the count of successful trajectories 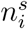 in FFS phase *i >* 0 is:

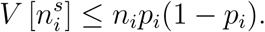

From Eq 3, the phase weight estimator 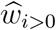 can be given in terms of the success count 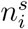:

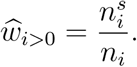

The variance of a quotient *V* [*x/y*] is *V* [*x*] */y*^2^ given that *y* is a constant^58^. Thus, the upper bound on the variance of 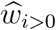 is:

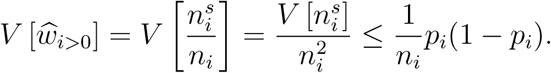

Finally, Eq 14 can be used to derive an upper bound on 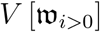 from the bound on 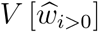:

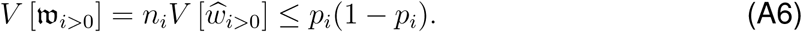

Parameterizing the FFPilot optimizing equation (Eq 30) with a value of *V* [𝔴*_i>_*_0_] that is at least as large as the true value helps to ensure that the calculated *n_i_* is at least sufficient to achieve the given error goal. The upper bound on the variance given in Eq A6 is equivalent to the variance of a single Bernoulli random variable with parameter *p_i_*. Thus, for the purposes of parameterizing the FFPilot optimizing equation we treat each 𝔴*_i>_*_0_ as a single Bernoulli random variable without any loss in accuracy.

## Appendix B: Blind Optimization Method

Taken as a whole, the FFPilot approach to optimizing simulations can function reliably only if the results of the pilot stage are highly accurate (at least in terms of the individual parameter estimates) and computationally inexpensive (relative to the production stage). No prior knowledge of the system under study is used in the setup of the pilot stage. Thus, a “blind” optimization method, one that uses no information about the current phase or any other, must be used during this initial stage.

The blind optimization method that we use during the FFPilot pilot stage works by altering the conditions under which a simulation phase is terminated. During a standard FFS simulation, simulation phase *i >* 0 is terminated once a fixed number of trajectories *n_i_* have launched from *λ_i−_*_1_ and have ended, regardless of where (in state space) those trajectories have ended. During a pilot stage, we instead terminate phase *i >* 0 only once a fixed number 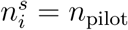 of *successful* trajectories (*i.e.* the ones that reach *λ_i_*) have been observed.

The advantage of using our blind optimization method is that it is able to produce estimates of the phase weight *w_i>_*_0_ = *p_i_* with constrained maximum error. The margin of error for a single phase *i >* 0 can be calculated from Eqs 20 and 28:

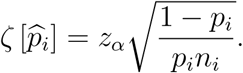

The total count of trajectories *n_i_* required to produce *n*_pilot_ successful trajectories in phase *i >* 0 converges to 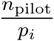. Thus, with respect to the pilot stage the above equation can be rewritten as:

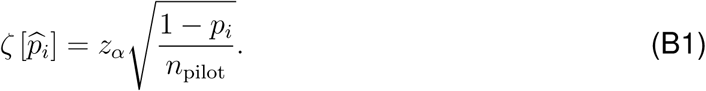

The error increases as *p_i_* becomes smaller, but it remains within a finite bound even as *p_i_* goes to zero (see supplemental Fig S10). By default and throughout this paper we use the fixed values of *n*_pilot_ = 10^4^ and *z_α_* = *z.*_95_*≈* 1.96 for all phases of the pilot stage. These values of *n*_pilot_ and *z_α_* give a maximum 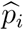 margin of error of 2%.

During phase 0, the distribution of samples taken from the underlying random variable w_0_ (*i.e.* the set of observed waiting times in between *λ*_0_ forward crossing events) is model dependent. This means that no formulation equivalent to Eq B1 is possible for the phase weight *w*_0_ = *τ_A_*. However, the estimator 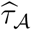 can be in general assumed to be 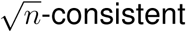 (as described in Secs II D and III A 1). Plugging Eqs 9 and 14 into Eq 6 gives the asymptotic margin of error for phase 0 of the pilot stage:

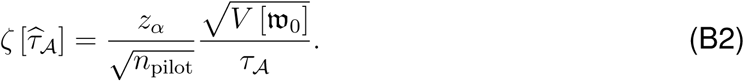

It can be ensured that 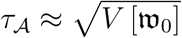 through appropriate placement of 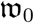 (*i.e.* away from a basin of attraction). Given that *τ_A_* and 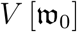 are appropriately matched, the margin of of error of phase 0 will be roughly proportional to 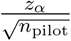. For *n*_pilot_ = 10^4^ and *z_α_ ≈* 1.96, this also works out to a 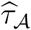 margin of error of about 2%. Thus, the blind optimization approach used in the FFPilot pilot stage controls error in 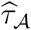, though not in such a conveniently bounded fashion as 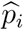.

## S1. ALTERNATIVE DERIVATION OF VARIANCE OF THE *MFPT* ESTIMATOR

Instead of using the delta method as in Sec III.A.1 of the main text, the variance of the forward flux sampling (FFS) *MFPT* estimator can also be derived from the fundamental properties of variance (though no explicit information about the underlying distribution of 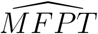 is gained this way). From the general properties of variance it is known that for independent random variables *X*_0_*, X*_1_*, …, X_i_*:

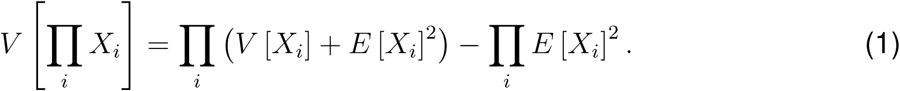

The righthand side of the above Eq 1 can be rewritten in terms of a generalized variance *G*^1^:

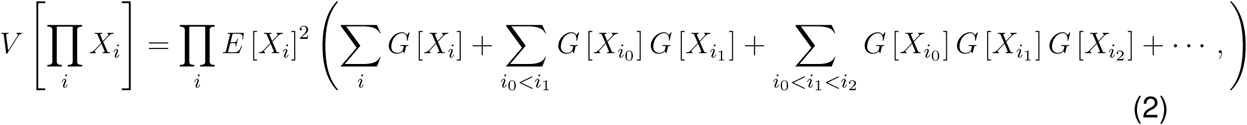

where 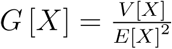. If the expected values are much larger than the variances, the higher order terms of 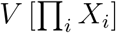 can be ignored without a large loss of accuracy. The condition:

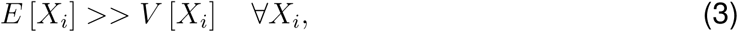

implies that:

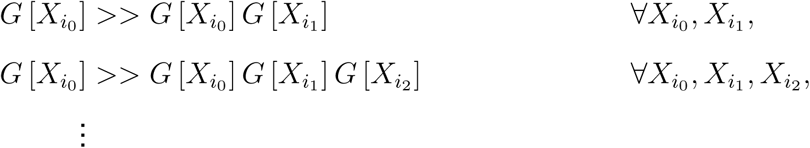

This regime yields an easy to work with approximation of Eq 2 as a series of of independent terms that only depend on a single *X_i_*:

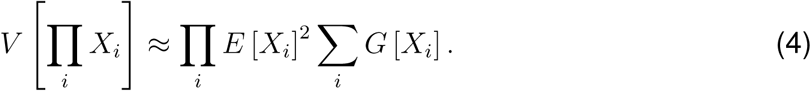

Each FFS simulation phase *i* can be conceptualized as taking a series of samples from a random variable 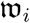 (see Sec II.A in the main text). The phase weight estimator 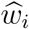 is then the mean of the 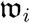 samples. The moments of each 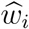 can be derived using the standard expected value and variance identities^2^, giving:

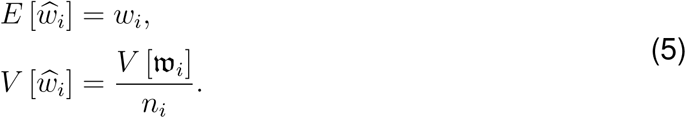

Since *V* [*w_i_*] is a fixed value, the value of each *V* [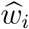] term trends monotonically downward as *n_i_* increases. If *n_i_* is assumed to be set to a large value, then it is reasonable to assume that 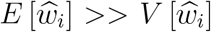 as well. In this regime the condition in Eq 3 is satisfied, and so we can apply the simplified formula for the product variance to the phase weights. Plugging the moments in Eq 5 into Eq 4 yields:

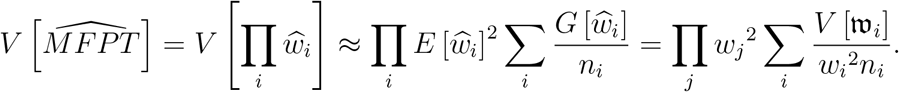

## S2. SUPPLEMENTAL FIGURES

**FIG. S1.**
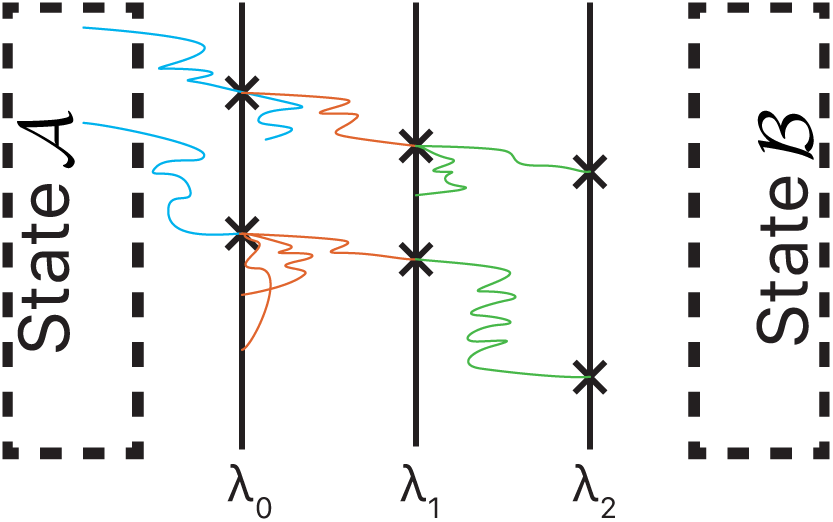
Schematic example of a pilot stage from a FFPilot simulation, with total phase count *N* = 3 and *n*_pilot_ = 2. Trajectories from phases (blue) 0, (red) 1, and (green) 2 shown in different colors.

**FIG. S2.**
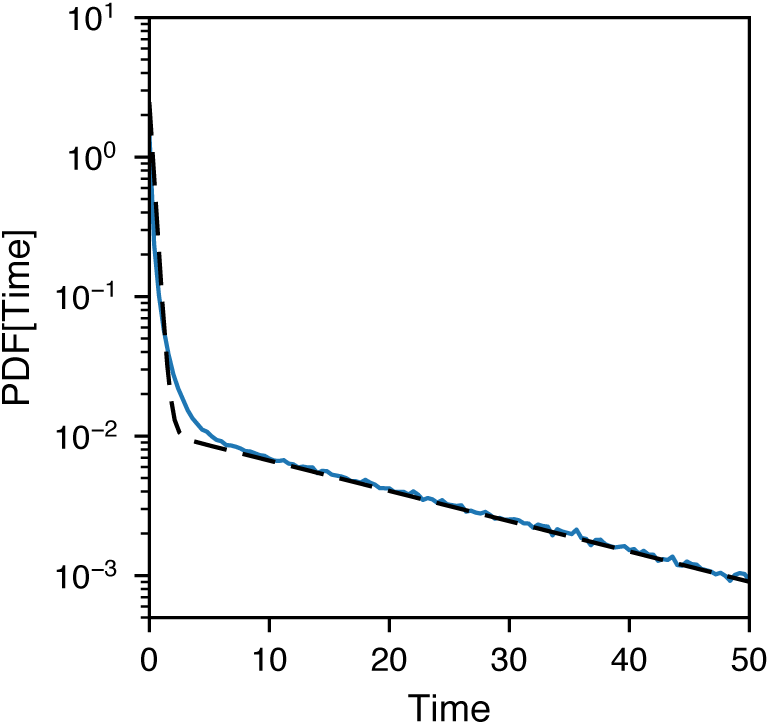
Distribution of waiting times in between phase 0 forward flux events for SRG_h=2.2_, 10^6^ samples. Dashed line is a fit of a mixture of two exponential distributions.

**FIG. S3.**
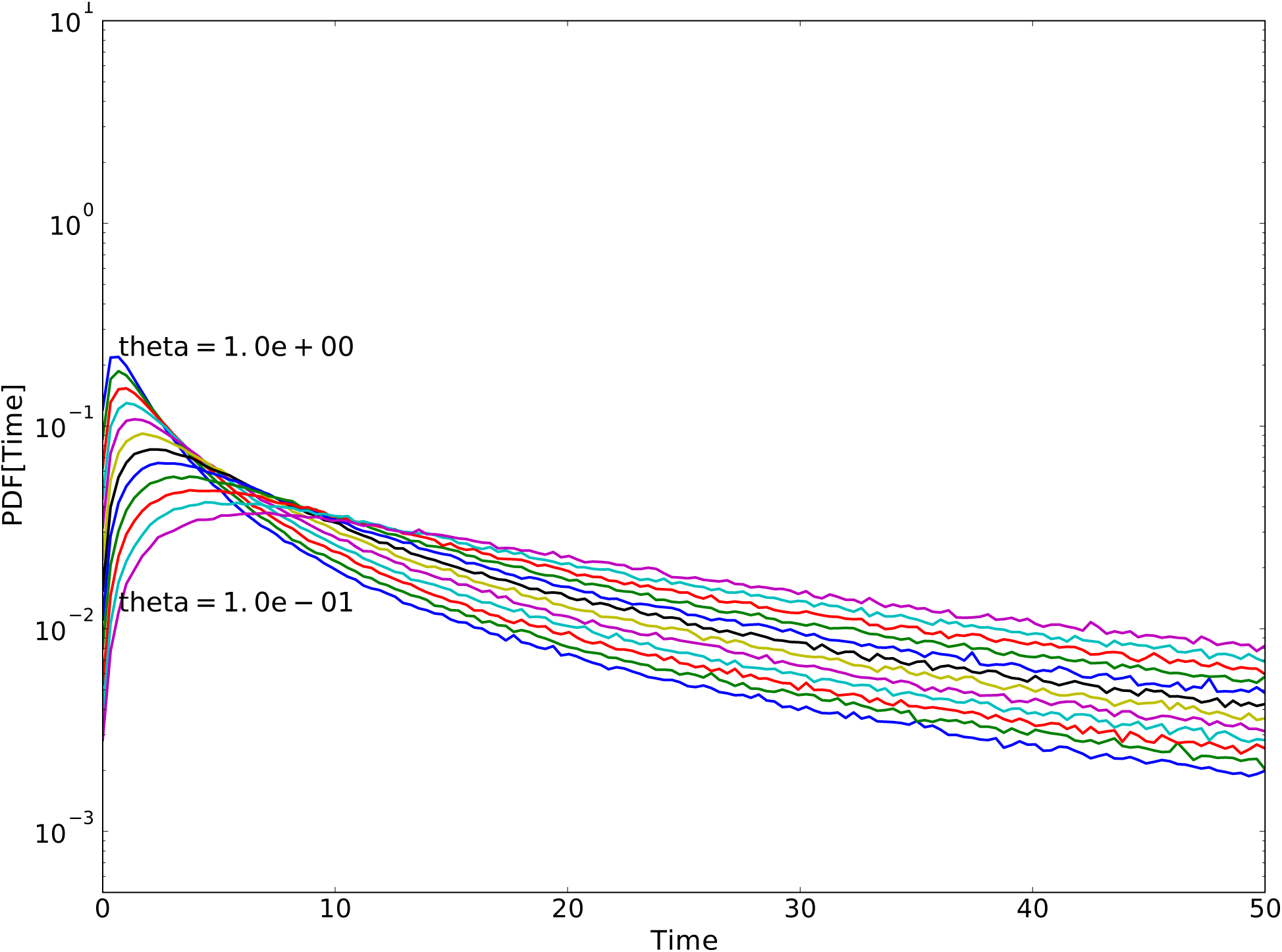
Genetic Toggle Switch phase zero inter-event time distribution for values of *θ* in between .1 and 10 (inclusive). 10^6^ samples in each distribution.

**FIG. S4.**
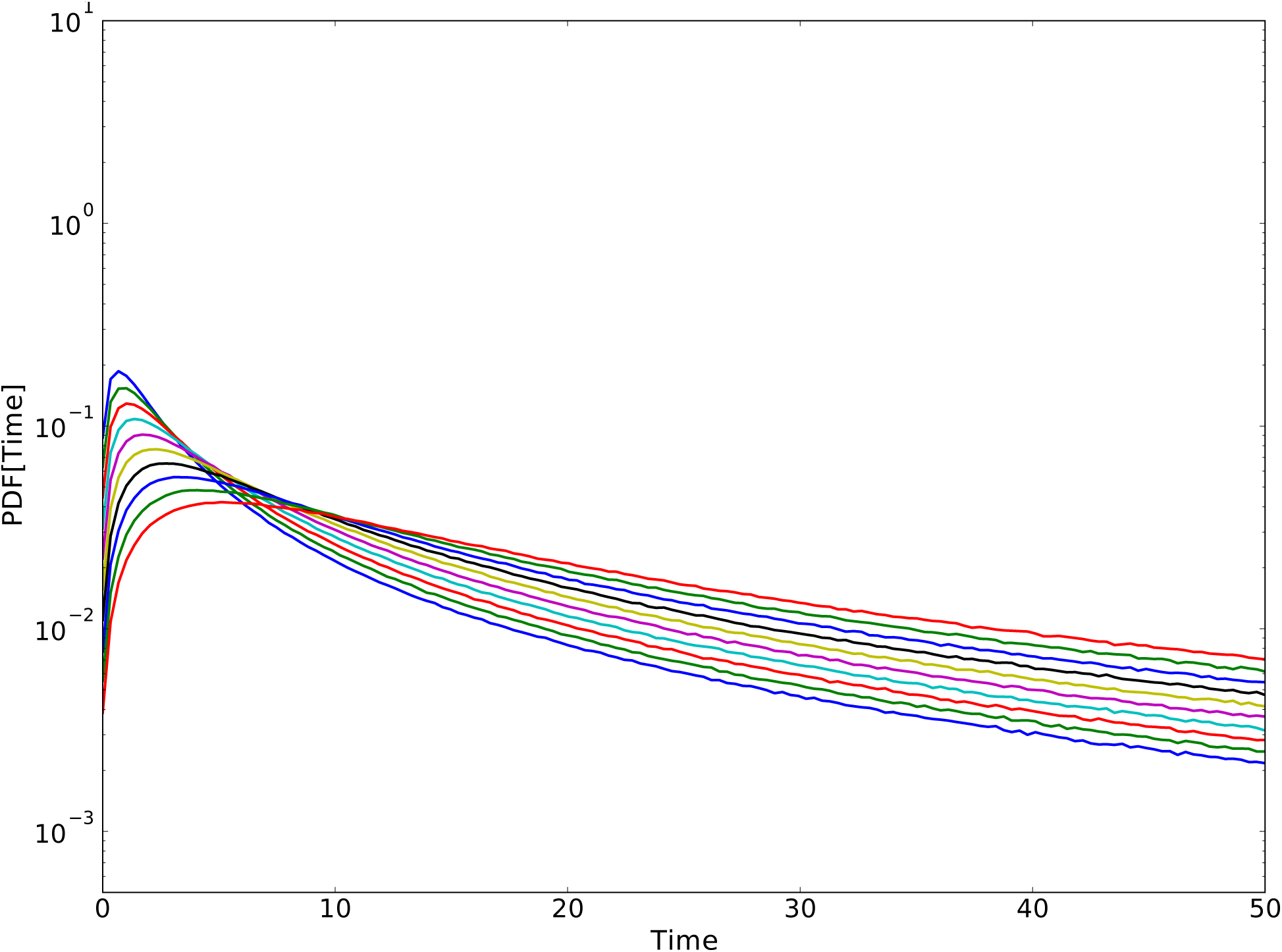
Genetic Toggle Switch phase zero inter-event time distribution for values of *θ* in between. 1and 10 (non-inclusive). 10^7^ samples in each distribution.

**FIG. S5.**
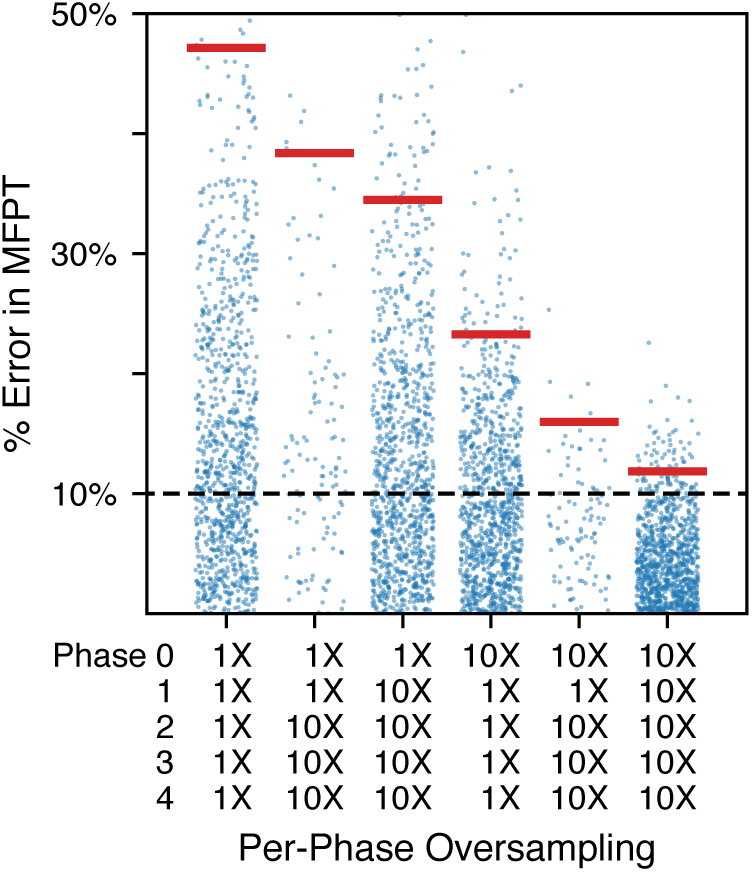
More results from simulations executed with various oversampling schemes. Aside from the use of 10X phase 0 oversampling instead of 20X, these simulations were equivalent to those presented in Fig 12 in the main text. As can be seen in the righthand column, 10X oversampling in every phase from 0 to 4 is almost, but not quite, enough to eliminate landscape error. The extent of oversampling in each separate scheme is indicated beneath each column of results. 1X sampling implies that standard number of FFPilot trajectories were sampled in that phase. For purposes of comparison, results from simulations run with no oversampling are included in the first column, and results from simulations run with 10X phase 0, 10X phase 1-4 oversampling are also shown. All simulations were of GTS*_θ_*_=10_ at an error goal of 10%.

**FIG. S6.**
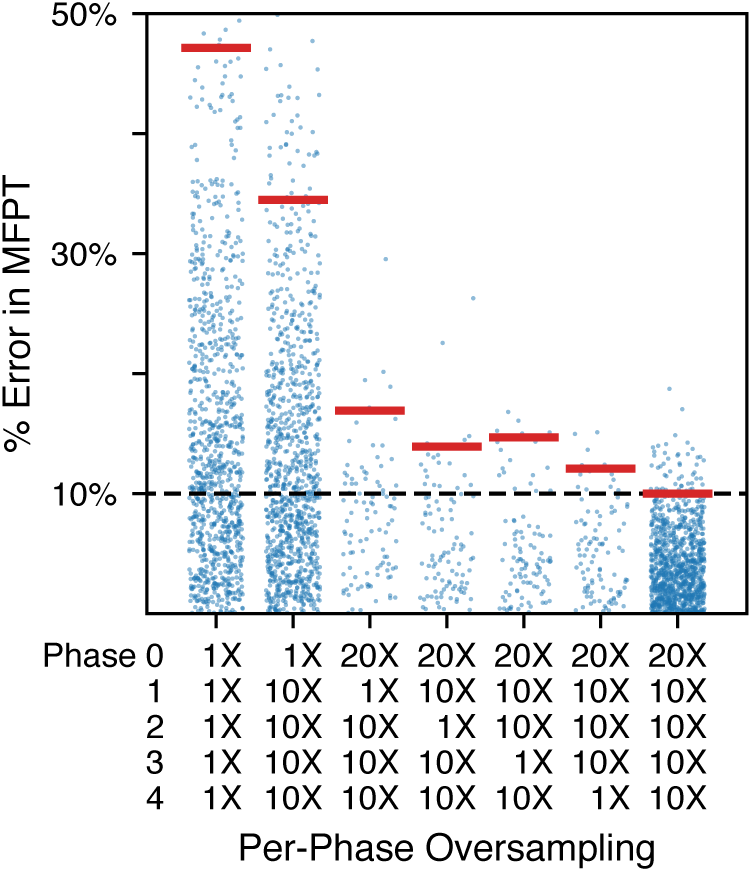
More results from simulations executed with various oversampling schemes. We wanted to investigate the effects of skipping oversampling in a single phase. Starting at the second column, results are shown from simulations in which we skipped oversampling in phase 0, then in the next column from simulations in which we skipped oversampling in phase 1, and so forth. The extent of oversampling in each separate scheme is indicated beneath each column of results. 1X sampling implies that standard number of FFPilot trajectories were sampled in that phase. For purposes of comparison, results from simulations run with no oversampling are included in the first column, and results from simulations run with 20X phase 0, 10X phase 1-4 oversampling (which is enough to eliminate landscape error) are also shown. All simulations were of GTS*_θ_*_=10_ at an error goal of 10%.

**FIG. S7.**
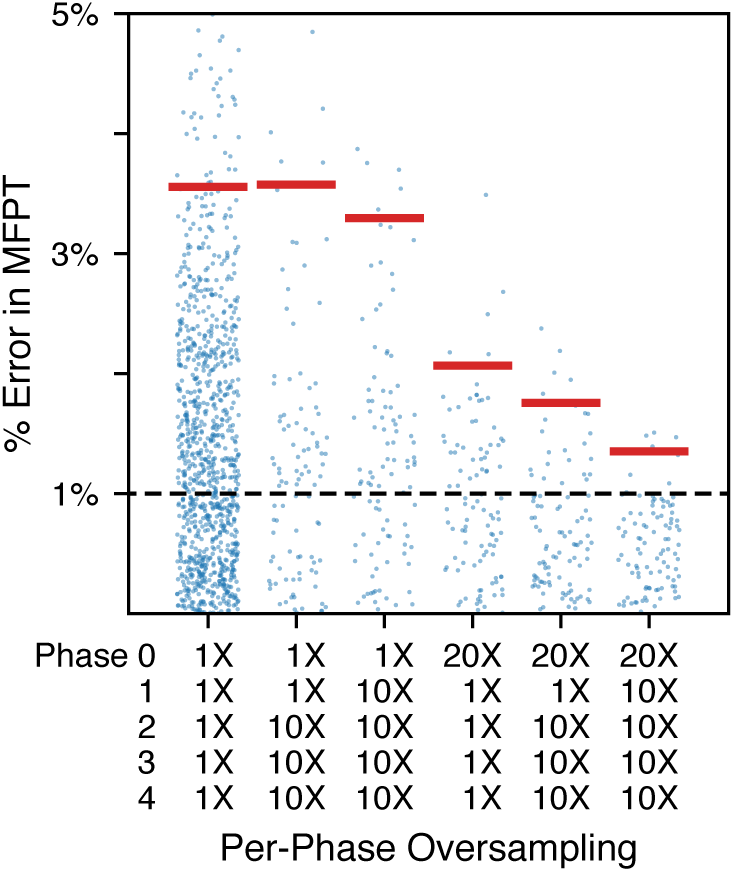
*MFPT* percent errors from a large set of simulations executed with various oversampling schemes. The simulations are similar to those presented in Fig 12 from the main text, except that all simulations were of GTS*_θ_*_=10_ at an error goal of 1% instead of 10%. All *MFPT* percent errors are calculated relative to the *MFPT* value estimated by a high accuracy DS simulation (.62% error goal).

**FIG. S8.**
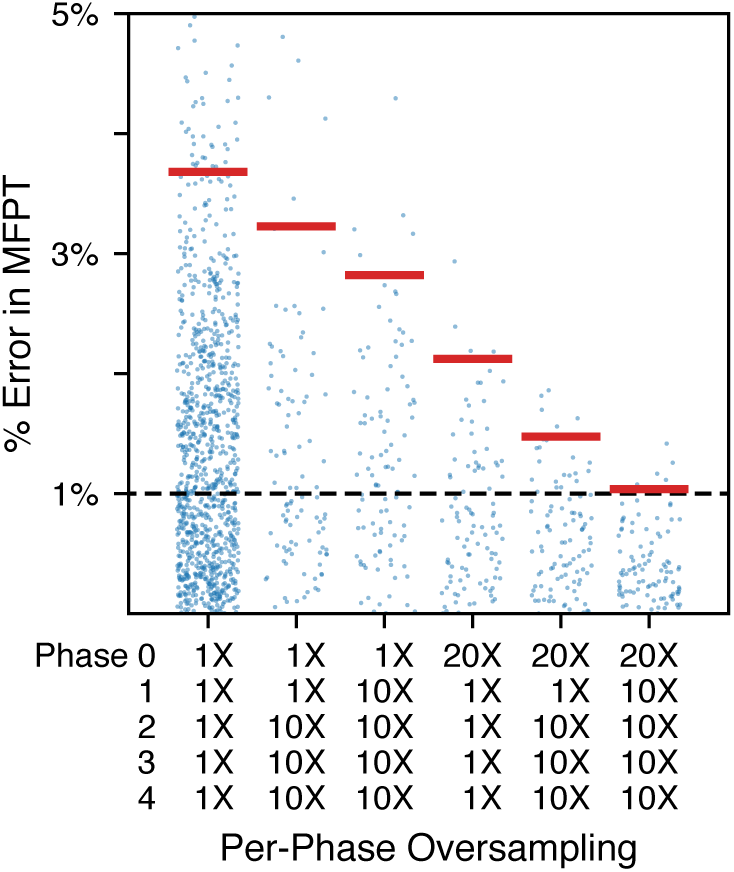
*MFPT* percent errors from a large set of simulations executed with various oversampling schemes. The simulations are similar to those presented in Fig 12 from the main text, except that all simulations were of GTS*_θ_*_=10_ at an error goal of 1% instead of 10%. All *MFPT* percent errors are calculated relative to the *MFPT* value estimated by a high accuracy FFPilot simulation (.1% error goal).

**FIG. S9.**
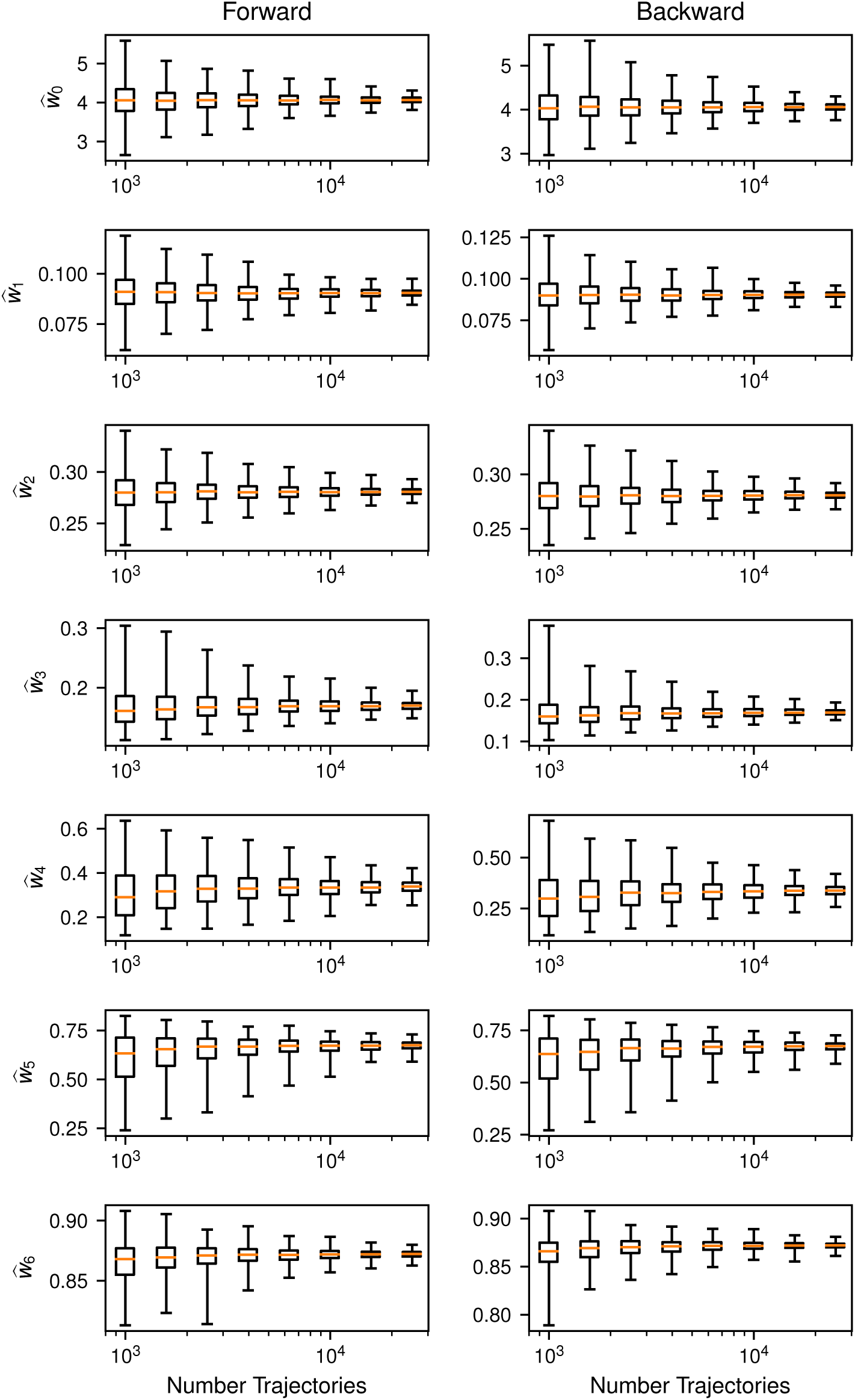
Phase weight estimators distributions for the GTS model with *θ* = 10 by the number of trajectories launched for each phase. Distributions were calculated from 100 independent FFS simulations.

**FIG. S10.**
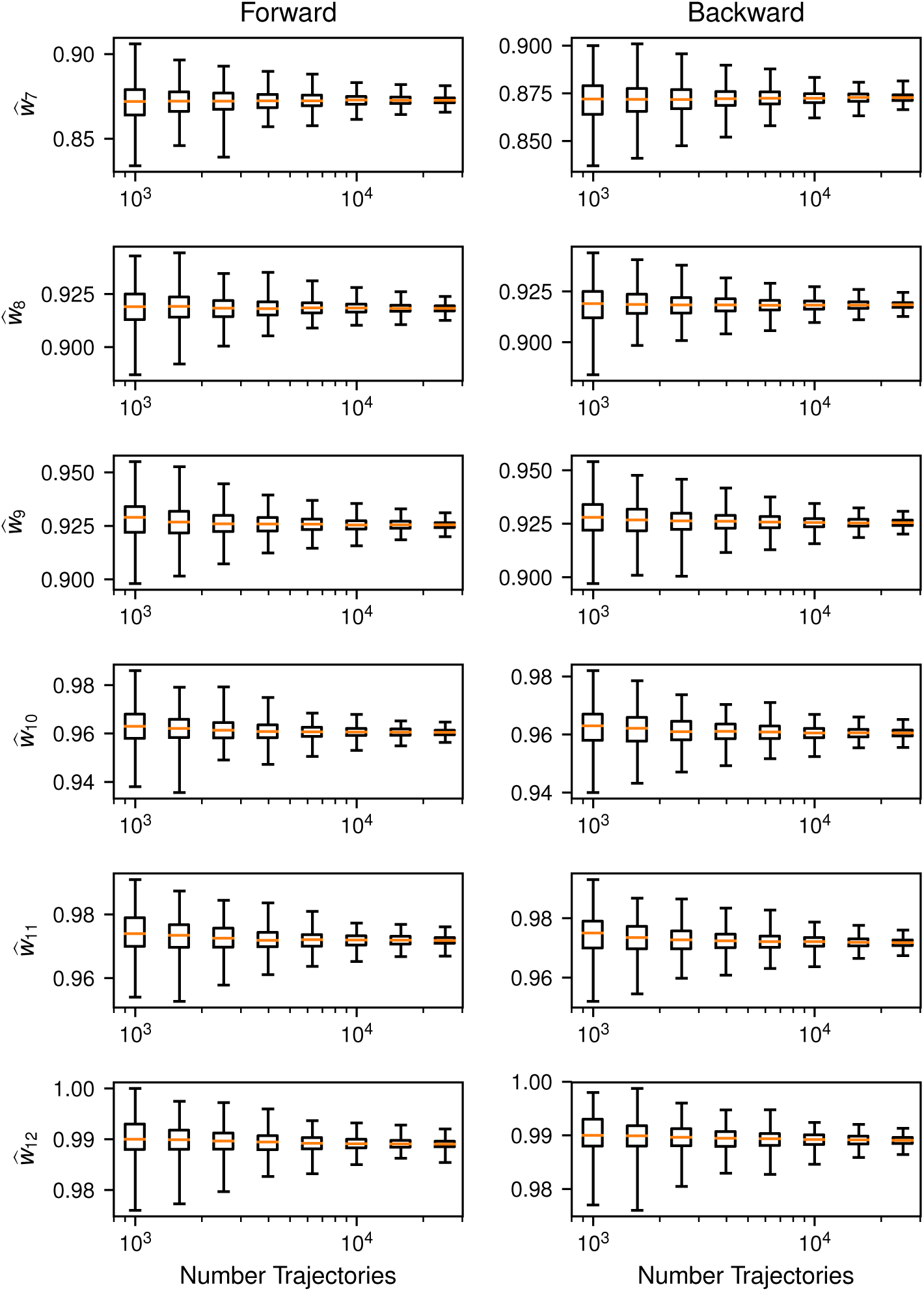
(cont) Phase weight estimators distributions for the GTS model with *θ* = 10 by the number of trajectories launched for each phase. Distributions were calculated from 100 independent FFS simulations.

**FIG. S11.**
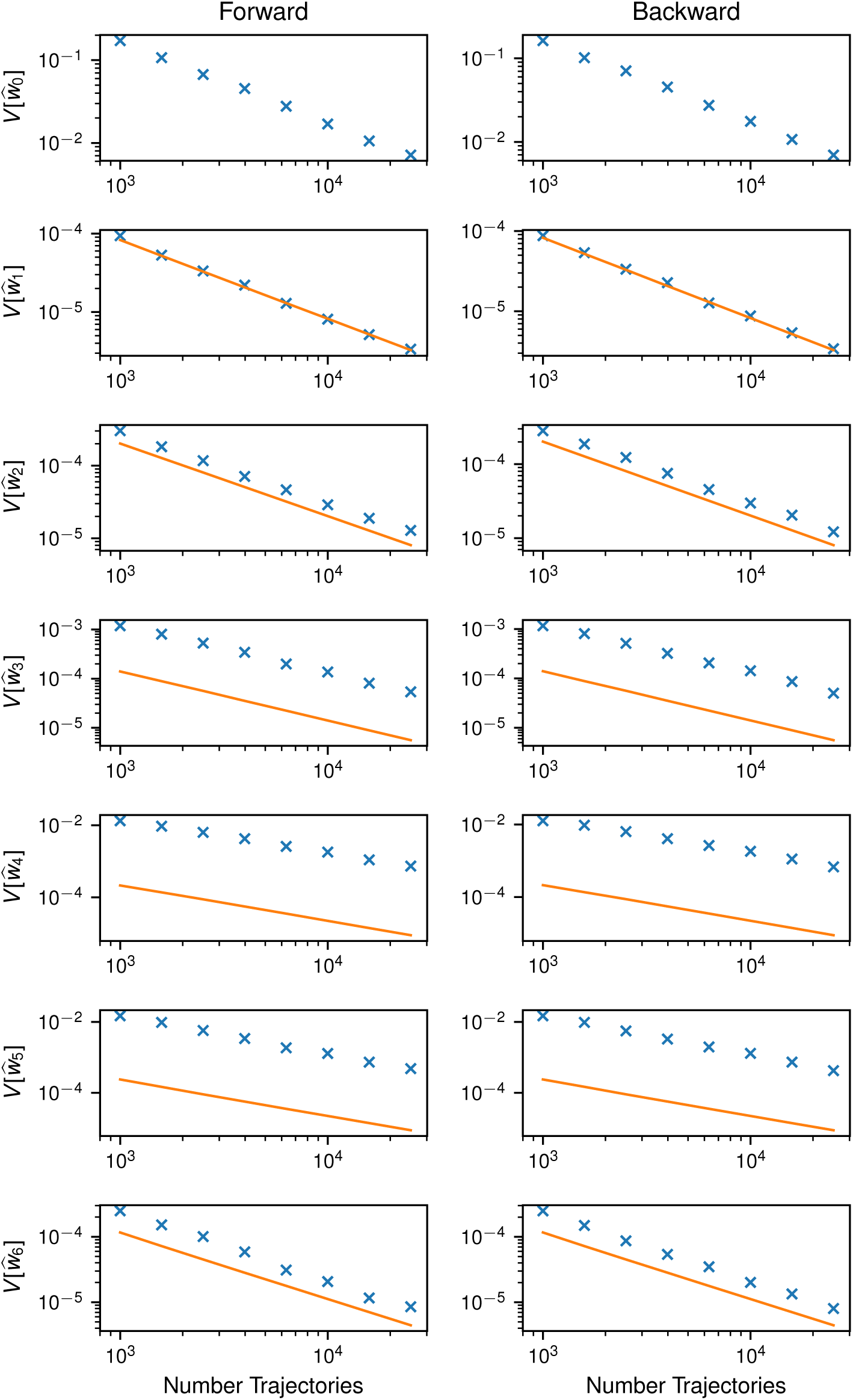
Variance of the phase weight estimator for the GTS model with *θ* = 10 by the number of trajectories launched for each phase. Variances were calculated from 100 independent FFS simulations. The solid line shows the expected variance from Eq 34.

**FIG. S12.**
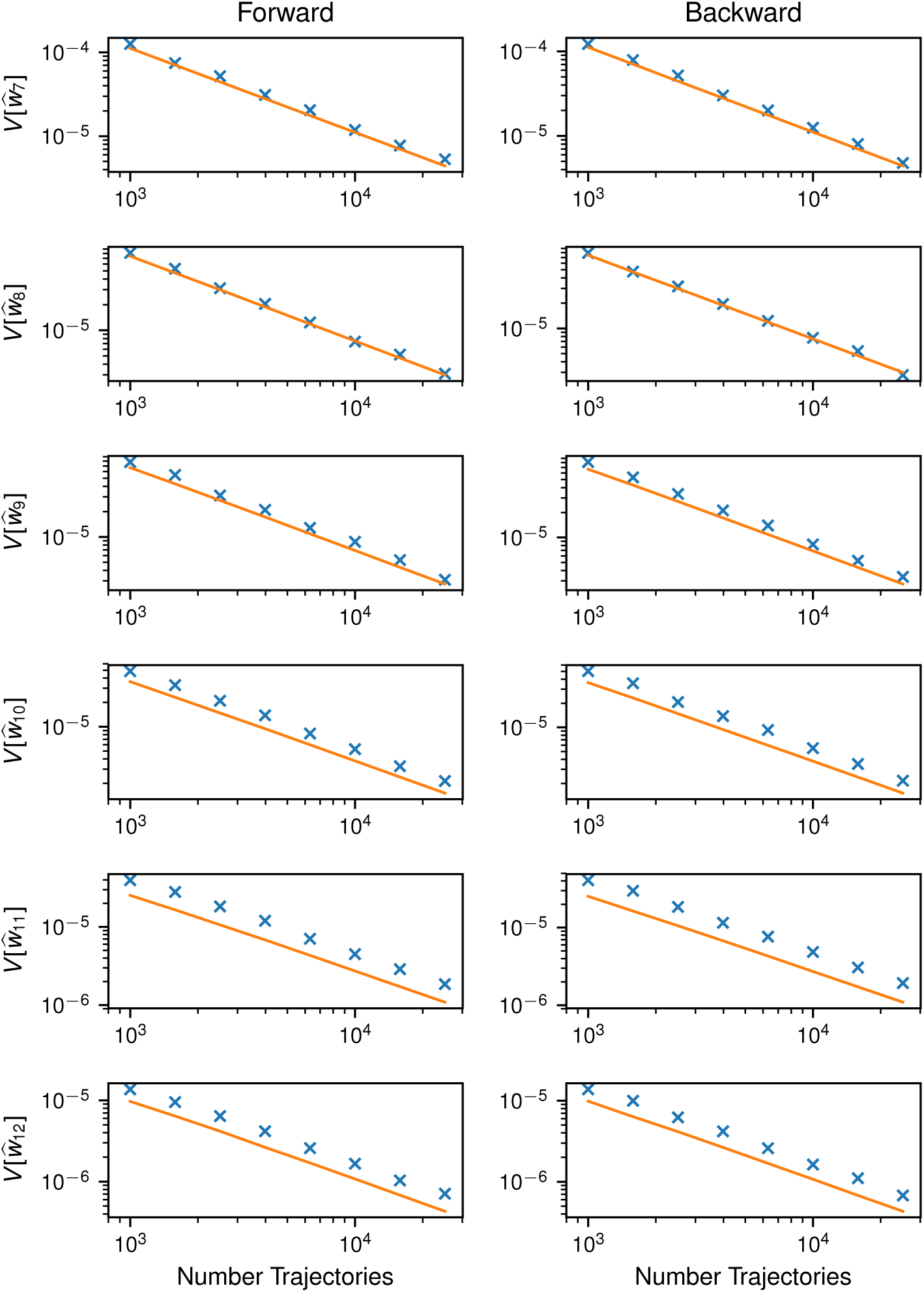
(cont) Variance of the phase weight estimator for the GTS model with *θ* = 10 by the number of trajectories launched for each phase. Variances were calculated from 100 independent FFS simulations. The solid line shows the expected variance from Eq 34.

**FIG. S13.**
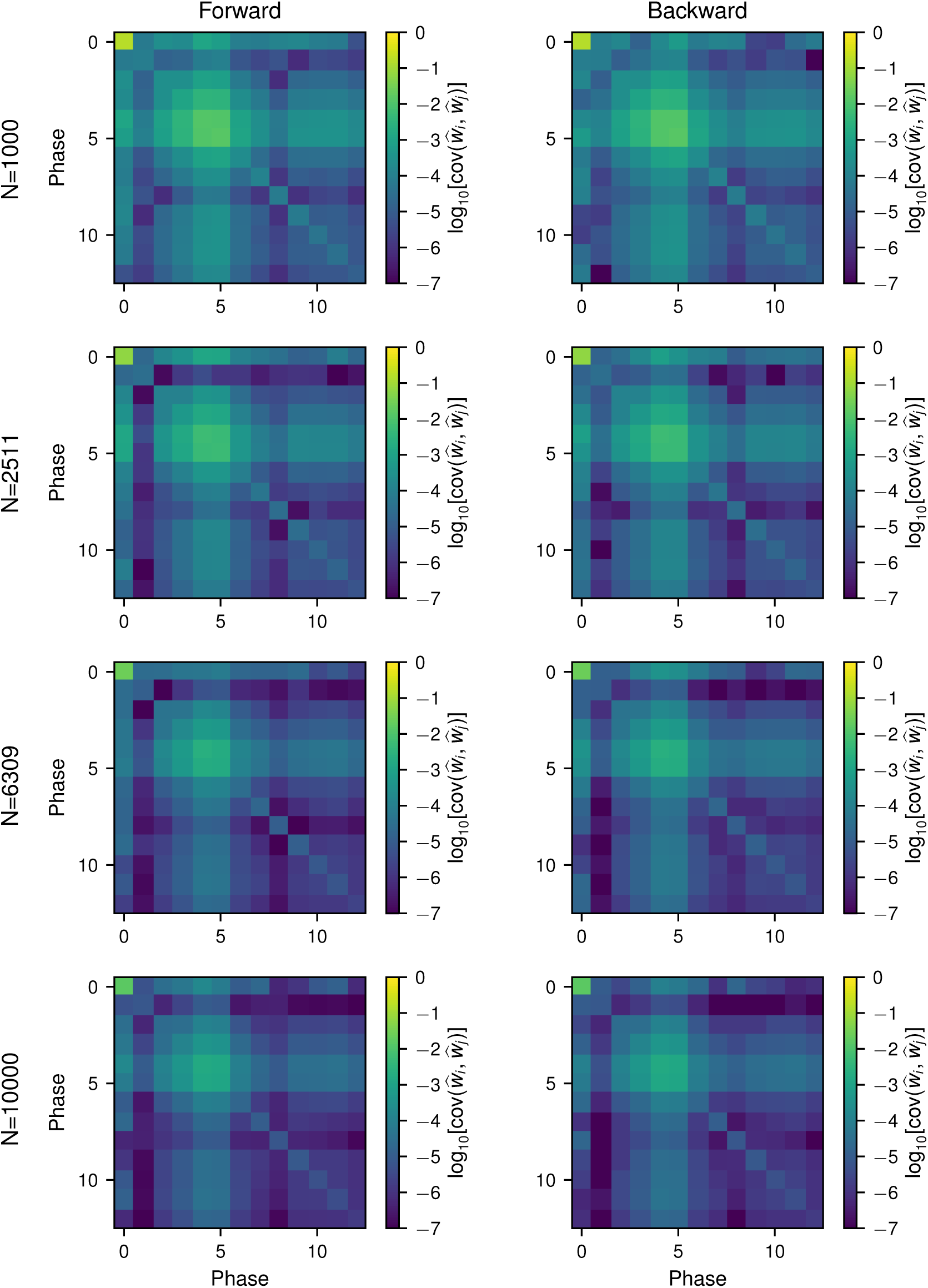
Covariances between the phase weight estimators for the GTS model with *θ* = 10. Rows show covariances for different numbers of trajectories N launched for each phase. Covariances were calculated from 100 independent FFS simulations.

**FIG. S14.**
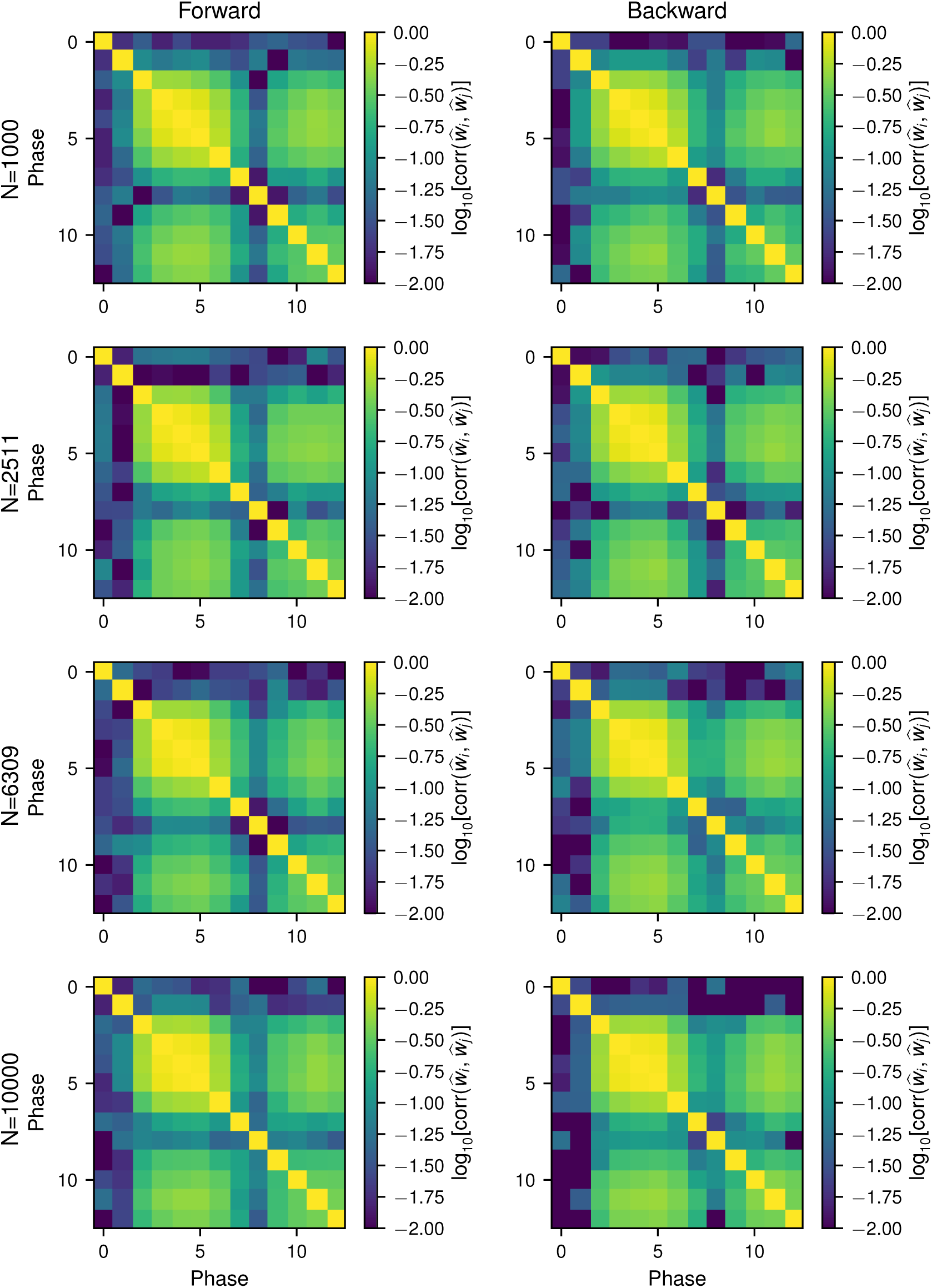
Correlations between the phase weight estimators for the GTS model with *θ* = 10. Rows show correlations for different numbers of trajectories N launched for each phase. Correlations were calculated from 100 independent FFS simulations.

**FIG. S15.**
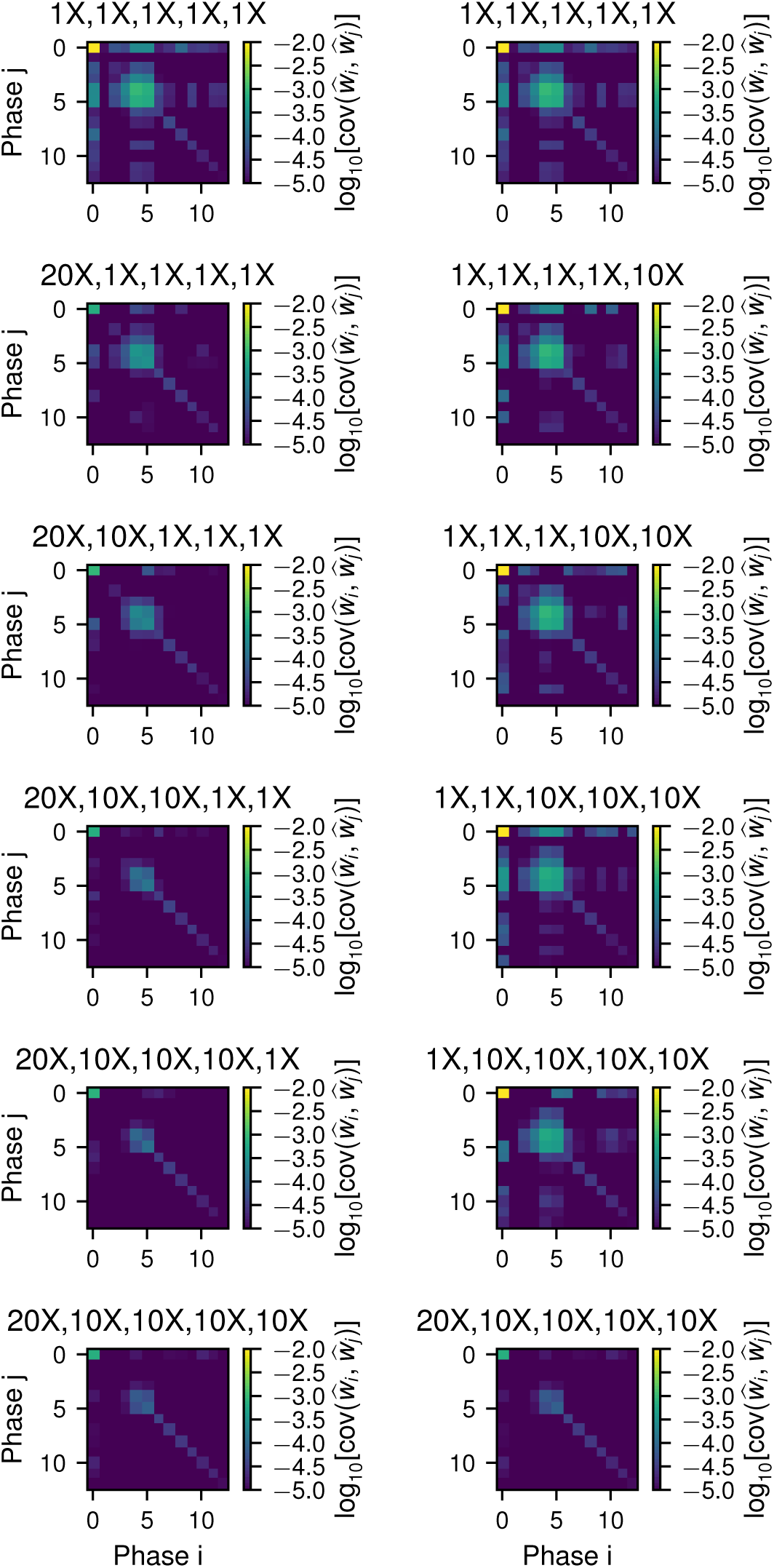
Effect of oversampling on the covariances between the phase weight estimators for the GTS model with *θ* = 10 and a 10% error goal. (left) Adding oversampling starting at phase 0 and working upwards. (right) Adding oversampling starting at phase 4 and working downwards. The oversampling factor for phases 0-4 is given in the title of each plot. Oversampling for phases 5+ was 1X.

**FIG. S16.**
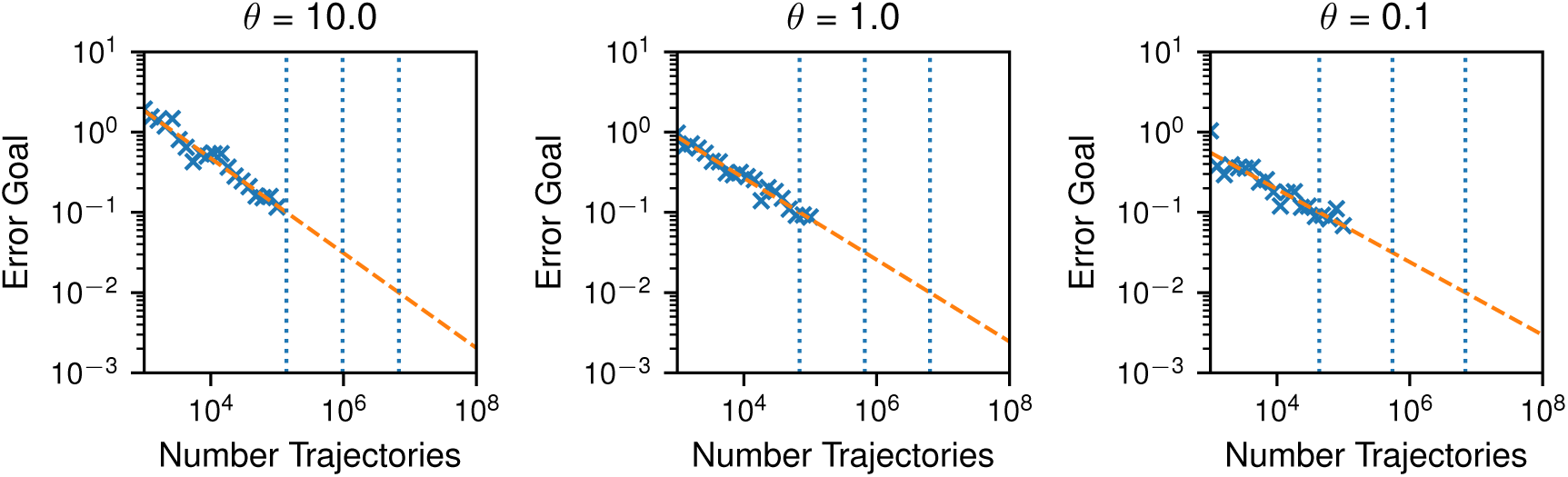
Number of trajectories launched per phase of traditional FFS vs margin of error, for the indicated GTS models. Symbols are values calculated from sets of FFS simulation. Dashed lines are linear fits in log-log space. The vertical dotted lines show the extrapolated optimal number of trajectories *N* to achieve a margin of error of 10%, 3.2%, and 1%, from left to right, respectively.

**FIG. S17.**
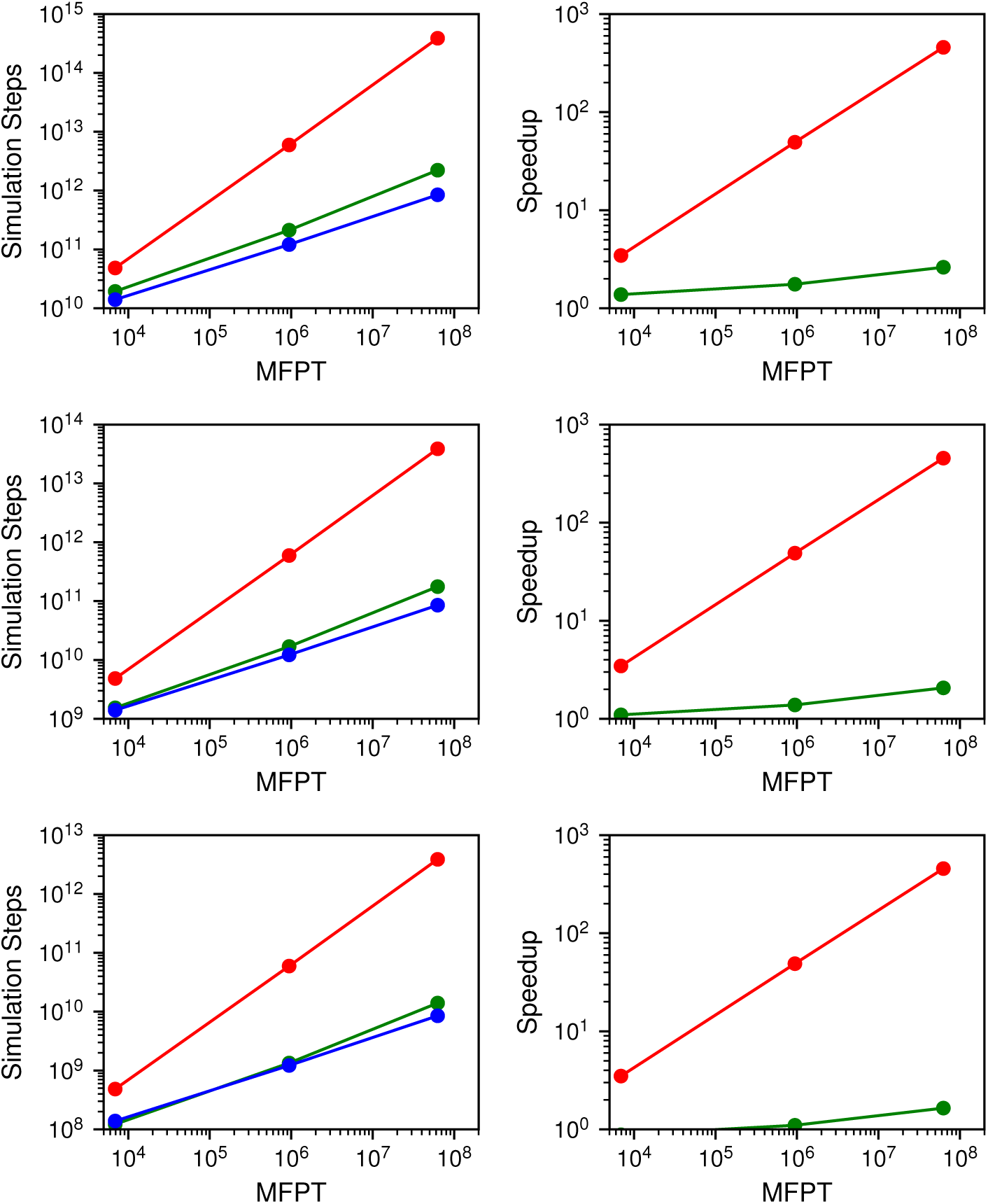
Performance comparison of (red) DS, (green) FFS, and (blue) FFPilot with matched margin of error for the three GTS models by MFPT. Rows show (top) 1%, (middle) 3%, and (bottom) 10% error goals.

**FIG. S18.**
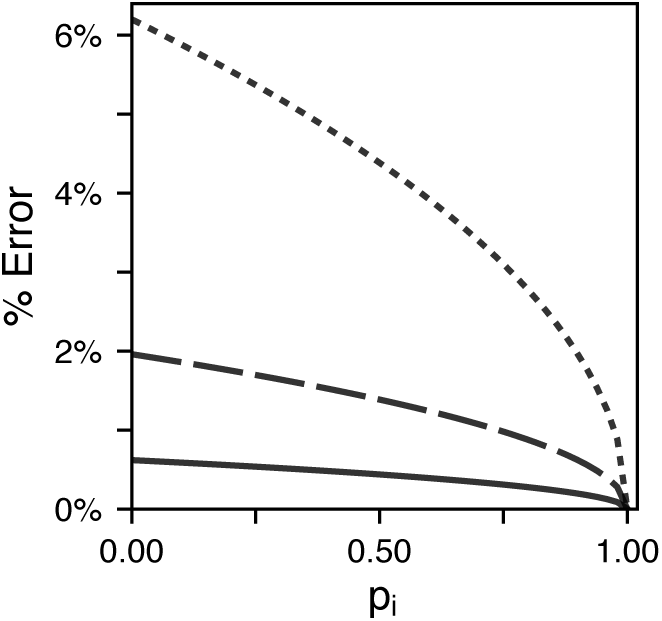
Maximum error (with respect to a single phase weight, 95% confidence) vs phase weight for a phase *i >* 0 when using the blind optimization method from the FFPilot pilot stage. Each of the lines shows the error for a different fixed value of the successful trajectory count, *n^s^* = *n*_pilot_. (solid line) *n*_pilot_ = 10^5^, (dashed line) *n*_pilot_ = 10^4^, (dotted line) *n*_pilot_ = 10^3^.

**FIG. S19.**
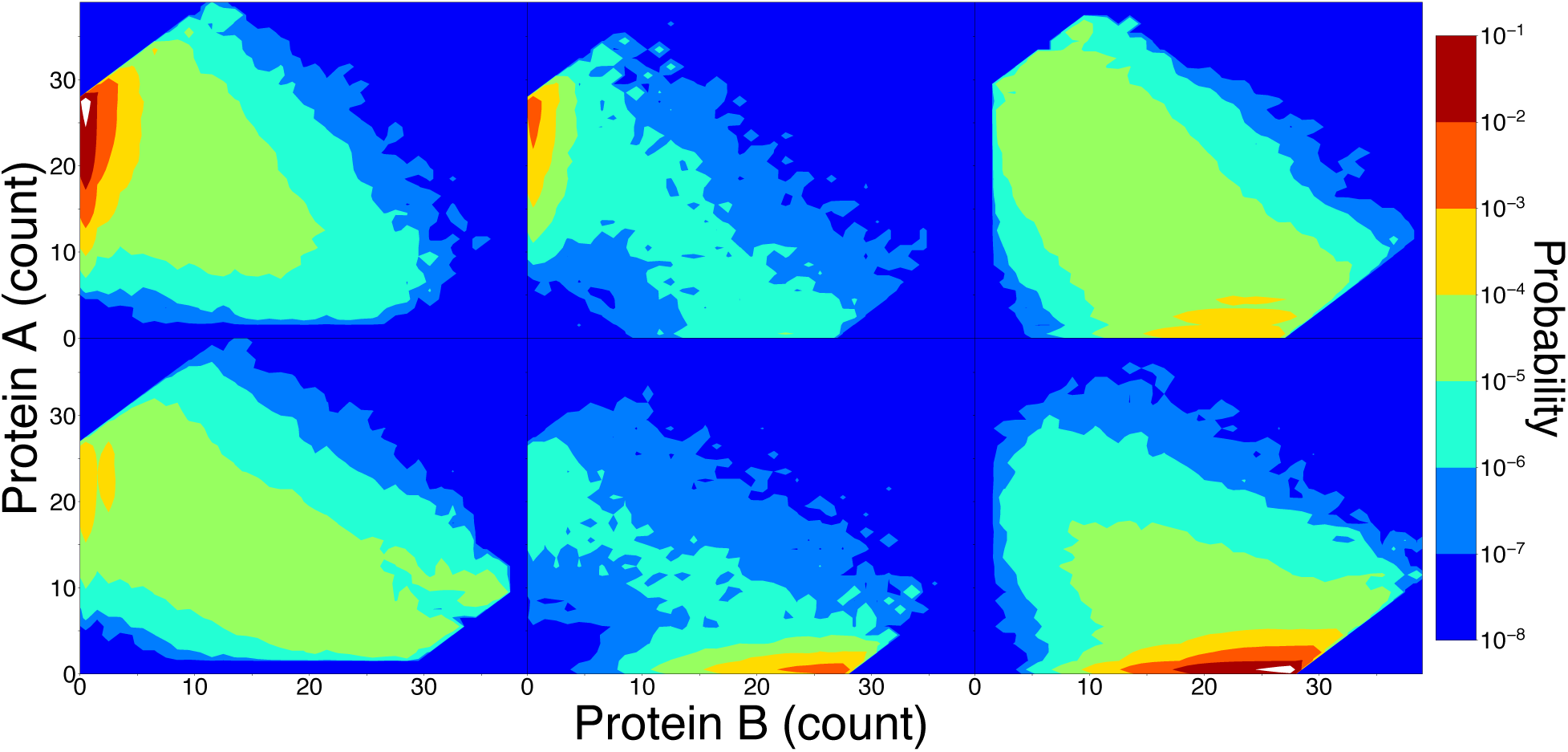
The state landscape of GTS*_θ_*_=10_, sliced by operator occupancy. The lefthand column shows the slice of the landscape in which an A dimer is bound to the operator DNA, the middle column shows the slice in which the operator is unbound, and the righthand column shows the slice in which the operator is bound to a B dimer. The top row is from a simulation of GTS*_θ_*_=10_ in which it switched 𝒜 → ℬ, and the bottom row is from a simulation in which it switched ℬ → 𝒜. Complex, high dimensional, and reasonably accurate landscapes can be produced in a straightforward fashion from relatively low cost FFPilot simulations. The 3 landscapes in each row represent the output of a single FFPilot simulation run to a 10% error goal, which ran to completion on a laptop in 5 minutes. The landscapes were analyzed and plotted using LMA, an analysis package written in Python and designed to work alongside the Lattice Microbes simulation suite.

## S3. SUPPLEMENTAL TABLES

**TABLE S1.**
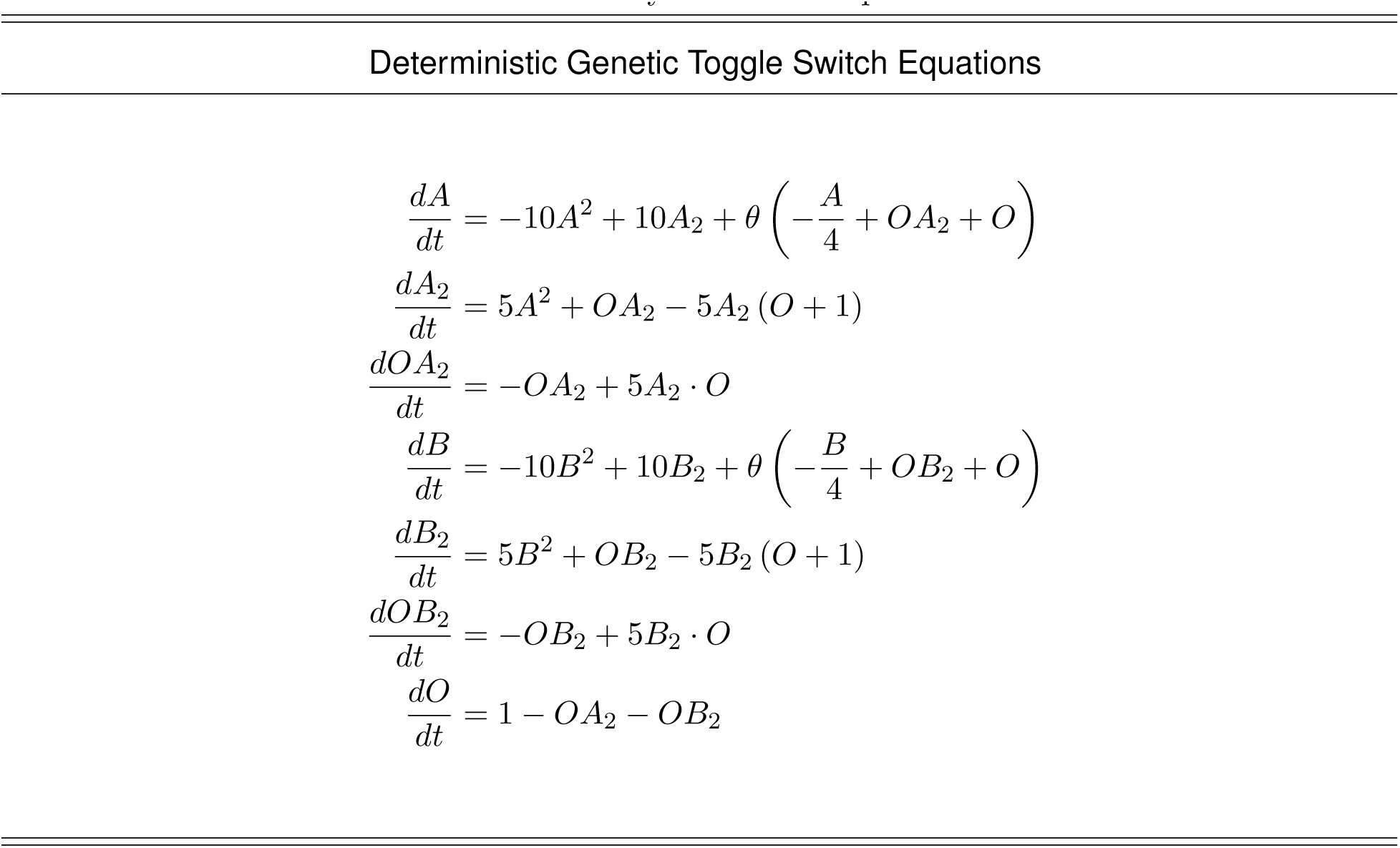
Deterministic ordinary differential equation model of the GTS.

| Deterministic Genetic Toggle Switch Equations |
| --- |
| $\frac{dA}{dt} = -10A^2 + 10A_2 + \theta \left( -\frac{A}{4} + OA_2 + O \right)$ $\frac{dA_2}{dt} = 5A^2 + OA_2 - 5A_2(O + 1)$ $\frac{dOA_2}{dt} = -OA_2 + 5A_2 \cdot O$ $\frac{dB}{dt} = -10B^2 + 10B_2 + \theta \left( -\frac{B}{4} + OB_2 + O \right)$ $\frac{dB_2}{dt} = 5B^2 + OB_2 - 5B_2(O + 1)$ $\frac{dOB_2}{dt} = -OB_2 + 5B_2 \cdot O$ $\frac{dO}{dt} = 1 - OA_2 - OB_2$ |

